# The T766M-EGFR lung cancer mutation promotes tumor growth by exploiting newfound mechanisms assembling ligand-free EGFR oligomer structures

**DOI:** 10.1101/2023.06.16.545259

**Authors:** R. Sumanth Iyer, Sarah R. Needham, Ioannis Galdadas, Benjamin M. Davis, Selene K. Roberts, Rico C.H. Man, Laura C. Zanetti-Domingues, David T. Clarke, Gilbert O. Fruhwirth, Peter J. Parker, Daniel J. Rolfe, Francesco L. Gervasio, Marisa L. Martin-Fernandez

## Abstract

Epidermal growth factor receptor (EGFR) is central to cell growth in physiology and pathophysiologies, including non-small cell lung cancer (NSCLC). EGFR has been successfully targeted with tyrosine kinase inhibitor generations, but the missense secondary T766M mutation is a common cause of resistance. Overcoming this therapeutic challenge has been hindered by poor understanding of how T766M dysregulates EGFR function leading to tumor progression. Here we show that T766M amplifies tumor growth *in vivo* by exploiting newly discovered oligomer assembly mechanisms employed by wild type (WT)-EGFR to maintain ligand-independent basal phosphorylation. These mechanisms, also shared by drug-resistant exon 20 EGFR insertions, reveal tumor growth promoting functions for hitherto orphan transmembrane and kinase interfaces and for the ectodomain tethered conformation of EGFR. Placing our findings into the context of a ligand-free oligomer structure model, we provide a framework for future drug discovery directed at tackling EGFR mutations in cancer by disabling oligomer-assembling interactions.

## Introduction

EGFR is a transmembrane tyrosine kinase receptor at the heart of signals for cell growth and division.^1^ EGFR is regulated on the cell surface by the binding of cognate growth factor ligands,^2^ which promote the assembly of a two-liganded back-to-back ectodomain dimer (B2B^ect^_dimer_) that underpins the formation of an asymmetric kinase dimer (Asym^kin^_dimer_)^3^ via structural coupling across the plasma membrane^4^ (**Figure 1A)**. The Asym^kin^_dimer_ is fundamental to catalyze EGFR auto­phosphorylation in C-terminal tyrosine residues, the trigger for EGFR-dependent downstream signaling pathways.^5^

**Figure 1:**
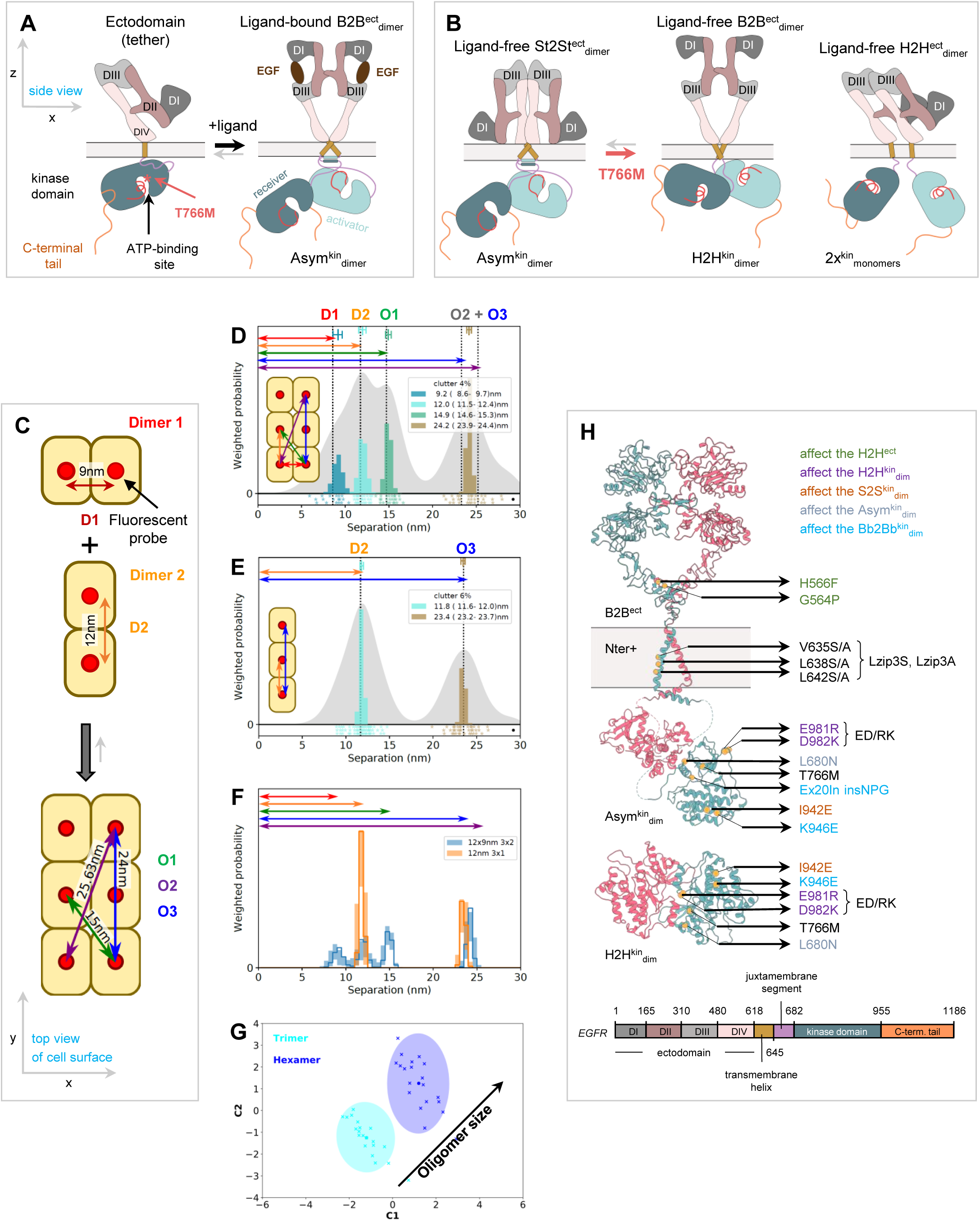
FLImP measurement of pairwise separations. **(A)** (*Left*) Cartoon of a ligand-free tethered ectodomain (subdomains numbered) linked to a kinase monomer; the ATP-binding site, the T766M mutation, and the C-terminal tail are marked. (*Right*) A two-liganded extended back-to-back ectodomain dimer (B2B^ect^_dimer_) and asymmetric kinase dimer (Asym^kin^_dimer_); an “activator” kinase stabilizes a “receiver” kinase in the active conformation.^3^ **(B)** (*Left*) A stalk-to-stalk ectodomain dimer (St2St^ect^_dimer_)^43^ coupled with the Asym^kin^_dimer_.^43^ (*Middle*) An extended B2B^ect^_dimer_^72^ coupled with a head-to-head kinase dimer (H2H ^kin^_dimer_)^45^. (*Right*) A head-to-head ectodomain dimer (H2H^ect^_dimer_) linked to two kinase monomers.^43^ **(C)** Cartoon of two imaginary dimers assembling a hetero^conf^-hexamer. Dimer and oligomer separations marked. **(D)** FLImP analysis of 100 synthetic pairwise separations for the hetero^conf^-hexamer (inset). (*Gray background*) Sum of posteriors of individual separations between fluorophores. (*Colored peaks*) Probability distributions of individual components (peaks) of decomposed separation distribution. The area under each peak is weighted according to the estimated proportion of measurements attributed to that peak (“abundance”). Results show agreement between ground truth and calculations for resolved separations. **(E)** As (D) for the homo-trimer formed by Dimer 2. **(F)** Comparisons between decomposed separation probability distributions between datasets. The continuous lines show the marginalized separation posterior for each condition, the sum of the abundance-weighted peaks in (D). The fluctuations around each continuous line arise from variations derived from FLImP decompositions for 20 bootstrap-resampled datasets to assess errors due to finite number of measurements. **(G)** Wasserstein MDS analysis. Similarities or dissimilarities between the 21 separation sets of different conditions (one main FLImP decomposition plus 20 bootstrap-resampled decompositions) are compared; in this case for the hetero^conf^-hexamer (D) and homo-trimer (E). The plot axes are Component 1 (C1) and Component 2 (C2). C1 represents the dimension that captures the largest amount of variance in the data, while C2 represents the second-largest amount of variance that is orthogonal to C1. The ellipse centers (95% confidence range) mark the positions of the main FLImP decompositions. The crosses mark the positions of individual bootstrap-resampled separation sets. **(H)** (*Top*) Map of mutations and treatments superimposed on the structure of a B2B^ect^_dimer_,^72^ an Asym^kin^_dimer_,^3^ and an H2H^kin^_dimer_^43, 61^. Mutations are colored according to the effect they are expected to have on the different dimers. (*Bottom*) EGFR sequence diagram.

Lung cancer is responsible for most cancer deaths worldwide.^6^ Somatic *EGFR* mutations in exons 18-21 are identified with high frequency (40%–60% in South-East Asian and 10%–20% in Caucasian patients with lung adenocarcinomas^7–9^) among a specific group of NSCLC patients with distinct etiology, prognosis and treatment approaches. In these patient groups EGFR has been successfully targeted with first-generation, ATP-competing tyrosine kinase inhibitors (TKIs),^10^ but resistance emerges with a frequency of about 50%-60%^11–13^ by the development of a dominant secondary T766M mutation in exon 20 at the entrance of the ATP-binding site^14^ (**Figure 1A**).

The T766M-EGFR mutation renders first-generation TKIs ineffective, in part by increasing ATP-binding affinity.^15^ Partial sensitivity is maintained with second-generation irreversible TKIs,^16^ like Afatinib,^17^ that form covalent bonds with a cysteine residue in the ATP-binding pocket, but their potency against WT-EGFR induces severe epithelium-based toxicity.^18^ This limitation was overcome with highly-selective, third-generation irreversible TKIs, such as Osimertinib (∼200-fold more potent towards T766M than WT-EGFR).^19, 20^ While the on-target (i.e., EGFR-dependent) resistance mechanism dominates during early-generation TKI treatment, it only accounts for 10–20% of patients treated with Osimertinib, whereas the C773S mutation represents the most common on-target resistance mechanism.^21–23^

Approaches to overcome resistance to third-generation TKIs rely on discoveries describing various off-target (i.e., EGFR-independent) as well as on-target resistance mechanisms. Strategies to overcome off-target resistance mechanisms are many-fold and include combining Osimertinib with MET inhibitors,^24, 25^ with Cyclin-dependent kinase 4/6 inhibitors,^26, 27^ with EGFR downstream signaling inhibitors (e.g. MEK inhibitors^28–30^), and with other TKIs (e.g. against AXL^31^ or FGFR1^32^), the latter remaining so far in the preclinical setting. Strategies to overcome on-target resistance mechanisms include combining first with third-generation TKIs, combining Brigatinib (dual ALK and EGFR inhibitor) with the anti-EGFR antibody Cetuximab,^33, 34^ and mutant-selective allosteric drugs, including EAI045,^35^ JBJ-09-063,^36^ CH7233163,^37^ and BLU-945.^38^ Notably, EAI045 prevents the kinase domain from adopting its active conformation and shows promise against the triple L834R/T766M/C773S mutant if combined with Cetuximab, which blocks EGFR dimerization.^35, 39^ However, toxicity-related concerns from off-target effects of Cetixumab limit the therapeutic potential.^40^

Despite the diversity in approaches currently under investigation, reliable therapeutic solutions against drug resistance remains elusive. One in-built weakness derives from the blanket effect on ATP-catalysis, which has developed into a cat-and-mouse chase because of the propensity of the kinase to mutate around the ATP-binding pocket. Another is that current approaches do not discriminate between homeostatic Asym^kin^_dimer_ signals regulated by ligand-binding and those sustained independently of ligand by somatic EGFR mutations that promote tumor growth.^41^ This lack of specificity has resulted in severe off-target effects.^42^ It is therefore paramount to investigate options that deliver a longer-lasting benefit with fewer side effects. One yet untested strategy is to interfere with specific events leading to the EGFR mutation-dysregulated ligand-independent state. In previous work we progressed towards characterizing the three different dimer conformers formed by ligand-free T766M-EGFR and WT-EGFR.^43^ This work exploited a super-resolution imaging method, Fluorophore Localization Imaging with Photobleaching (FLImP),^44^ guided by protein structures and MD simulations. We reported that in the absence of ligand the Asym^kin^_dimer_ is instead structurally coupled on the cell surface to a ligand-free stalk-to-stalk ectodomain dimer (St2St^ect^_dimer_)^43^ (**Figure 1B**) and that this active dimer coexists with two additional ligand-free inactive dimers: a back-to-back ectodomain dimer structurally coupled to a symmetric head-to-head kinase dimer (B2B^ect^/H2H^kin^_dimer_),^45^ and a head-to-head ectodomain dimer linked to two non-interacting kinase monomers (H2H^ect^/2x^kin^_monomers_)^43^ (**Figure 1B**). The counterintuitive finding that the T766M mutation, despite inducing constitutive signaling, destabilizes the catalytic Asym^kin^_dimer_ in favor of the inactive H2H^kin^_dimer_^43^ suggests that ligand-free dimer conformers must somehow interact to assemble hetero-conformational-oligomers (hetero^conf^-oligomers). However, the previous ∼5 nm resolution of FLImP prevented us from determining if or how. The Asym^kin^_dimer_ is also disfavored by the WT-EGFR kinase.^46, 47^ Moreover, the Asym^kin^_dimer_, when artificially joined to a ligand-free ectodomain dimer, is disordered in electron micrographs.^48, 49^ Consequently, we speculated that the St2St^ect^/Asym^kin^_dimer_ may form only within a hetero^conf^-oligomer whilst reinforced by interactions with the B2B^ect^/H2H^kin^_dimer_. If true, we conjectured that understanding the underlying oligomer-assembling interactions may reveal a potential Achilles heel in T766M-driven tumors that could be therapeutically targeted.

Here we implement a higher-resolution FLImP version, enabling us to probe interactions between the three ligand-free EGFR dimers and uncover the assembly mechanisms of dysregulated ligand-free hetero^conf^-oligomers. Combined with large-scale simulations of various membrane-embedded dimer interfaces, we build an experimentally-backed model of all the relevant interactions, showing that WT-EGFR and T766M-EGFR share a ligand-free hetero^conf^-oligomer structure. WT-EGFR and T766M-EGFR exploit the ectodomain tethered conformation^50^ to form H2H^ect^/2x^kin^_monomers_, and a transversal transmembrane interface^51^ to form tetramer-scaffolds between the H2H^ect^/2x^kin^_monomers_ and B2B^ect^/H2H^kin^_dimer_. These tetramer scaffolds cantilever into position the two halves of the extracellular portion of the St2St^ect^_dimer_ enabling the coupled Asym^kin^_dimer_ to form. Within these hetero^conf^-oligomers, the Asym^kin^_dimer_ is positively and negatively regulated via two currently functionally-orphan kinase interfaces revealed by X-ray crystallography (PDB IDs:3VJO^52^ and 5CNO^53^). Crucially, we show that some of these newly-found protein-protein interfaces, specific to the ligand-free state, are critical for T766M-EGFR-dependent growth *in vivo*.

## Results

### Quantitative fingerprinting of dimers and oligomers

FLImP measures the lateral pairwise separations on the scale of 0-70 nm between fluorescent probes bound to cell surface dimers and oligomers and their relative abundance (**STAR methods**).^44^ Mimicking the viewpoint of FLImP microscopy, we can assume the orthogonal projection onto the cell surface (xy-plane) of different conformation transmembrane Dimers 1 and 2, specifically labelled with a fluorescent probe, which form a hetero^conf^-hexamer (**Figure 1C)**. We created a dataset of synthetic FLImP separation probability distributions simulating measurements of individual separations between these fluorophores, including spurious fluorophore localizations (clutter), and noise, which we summed (**Figure 1D**; gray background). FLImP decomposes pairwise probe distances separated from each other by <3 nm, namely 1^st^-order (dimer interfaces) (D1, D2) and the short diagonal (O1), but not those separated by 1.6 nm, namely the long diagonal (O2) and 2^nd^-order vertical (O3) (**Figure 1D**).

We also simulated synthetic FLImP separations for the homo-trimer that forms when Dimer 1 is disrupted (**Figure 1E**, gray background). FLImP decomposes from these data the remaining 1^st^ and 2^nd^-order vertical separations from the homo-trimer (D2, O3). Reflecting the structural change, the D2 and O3 peaks change in intensity, and O3 shifts position because the unresolved contribution from O2 is no longer present (**Figure 1E)**. Changes in peak intensity and position like these are markers for protein-protein interactions in oligomers and those that we aim to resolve and assign to structures using FLImP on cells.

FLImP samples a finite population of separations, and this introduces errors. We, therefore, used bootstrap-resampling^54^ to estimate how this affects the decomposition (**STAR methods**). **Figure 1F** illustrates that bootstrap-resampling can capture changes caused by the transition from hetero^conf^-hexamer to homo-trimer and evaluate significance above finite sampling errors.

Given enough resolution and with the help of mutations/treatments, separation peaks from dimers/1^st^ order interfaces are typically amenable to be assigned, but higher order peaks can be harder. We interrogate unassigned higher order peaks by folding them into a complementary multidimensional scaling (MDS) Wasserstein metric.^55^ This measures the work needed to convert a decomposed separation set into another, thereby estimating similarities and differences between whole FLImP separation decompositions (**STAR methods**). As shown in the example (**Figure 1G**), we also include the bootstrap-estimated errors associated with finite sampling, and calibrate changes to report oligomer growth direction.

The mutations and treatments that we used to dissect the protein-protein interactions assembling ligand-free oligomers are mapped-out in **Figure 1H**. Given the number of mutations, automating acquisition and data was paramount (**STAR methods**).

### The B2B^ect^/H2H^kin^_dimer_ underpins the formation of the St2St^ect^/Asym^kin^_dimer_

The color peaks in **Figure 2A** show the most likely positions and intensities of pairwise separations between CF640R fluorophores specifically conjugated to anti-EGFR Affibodies bound to DIII of cell surface WT-EGFR ectodomains. All FLImP separation sets have hereafter the median positions of the WT-EGFR separations superimposed (dashed lines) to facilitate comparisons. FLImP measurements require immobilizing receptors on the cell surface by chemical fixation using a method previously demonstrated not to introduce detectable artefacts.^44^ Nevertheless, predictions arising from results of chemically fixed cells were validated in live cells using single particle tracking (**Figure S1A**).

**Figure 2:**
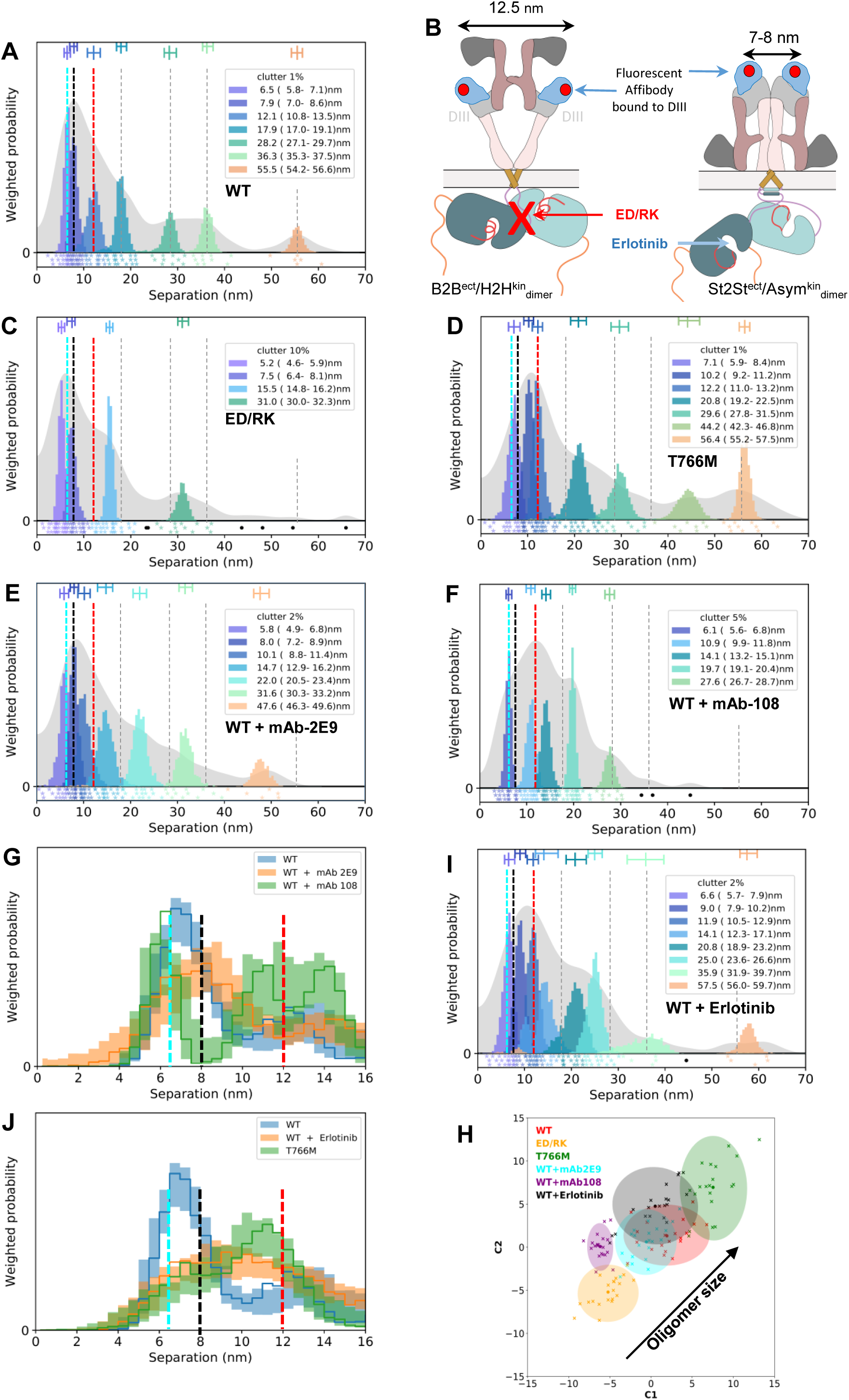
The B2B^ec^7H2H^kin^_dimer_ underpins the formation of the St2St^ec^7Asym^kin^_dimer_. **(A, C-F, I)** FLImP analysis of 100 separation probability distributions between Affibody-CF640R pairs in the conditions indicated: (*Gray background*) As in Figure 1(D). (*Colored peaks*) As in Figure 1(D). The median peak positions marked by dashed lines are hereafter also superimposed on all the FLImP separation diagrams to facilitate comparisons with the WT. **(B)** (*Left*) Cartoon of a labeled B2B^ect^/H2H^kin^_dimer_ showing two fluorescent anti-EGFR Affibody bo und to the two DIII of the ectodomains. The ED/RK mutations break the H2H^kin^_dimer_. (*Right*) A labelled St2St^ect^/Asym^kin^_dimer_. Erlotinib binds the ATP pocket of the kinase stabilizing the Asym^kin^_dimer_. **(G, J)** Comparisons as in Figure 1(F). **(I)** Wasserstein MDS analysis comparing the 21 separation sets (main FLImP set + 20 bootstrap-resampled) of the conditions marked in the plot. Axis labels and 95% confidence ellipses as in caption Figure 1G.

Interpretation of FLImP separation sets requires assigning peak positions to dimer and/or protomer interaction interfaces in hetero^conf^-oligomers. We interrogate this (see below) with the help of mutations and treatments designed to break or reinforce previously reported dimers. In previous work,^43^ we linked the 10.8-13.5 nm peak (**Figure 2A**, red dashed line) to the B2B^ect^/H2H^kin^_dimer_ (**Figure 2B**). This link is supported here by the observation that FLImP no longer indicates separation density between 8.1-14.8 nm when a previously proposed double E981R/D982K (ED/RK) mutation in the C-terminus^56^ that destabilizes electrostatic interactions at the heart of the H2H^kin^_dimer_ is introduced (**Figure 2C**; red dashed line). By contrast, separation density at 9.2-13.2 nm increases when the B2B^ect^/H2H^kin^_dimer_ is stabilized by the T766M mutation (**Figure 2D**; red dashed line), further supporting the assignment of the 10.8-13.5 nm peak in the WT-EGFR separation set to the B2B^ect^/H2H^kin^_dimer_.

We previously measured a <9 nm separation for the St2St^ect^/Asym^kin^_dimer_.^43^ Our enhanced FLImP method suggests two <9 nm components in the separation set of the WT-EGFR (**Figure 2A**; cyan and black dashed lines). To determine if one of these components corresponds to the St2St^ect^/Asym^kin^_dimer_, we pre-treated WT-EGFR-expressing cells with the conformation-selective monoclonal antibodies mAb-2E9^57^ or mAb-108,^58^ which select for high and low-affinity Epidermal Growth Factor (EGF) binding, respectively. As the selectivity of these mAbs is bona fide against EGF binding, mAb-treated cells were next probed with an EGF-CF640R derivative. Here cells were fixed after mAb treatment but before probing with EGF-CF640R to avoid ligand-induced conformational changes. Results show that EGF-CF640R binds well to fixed cells at similar sites to Affibody-CF640R (**Figure S1B, S1C**).

Previous work^59^ together with our results suggest that EGF binds St2St^ect^/Asym^kin^_dimer_ with high affinity (**Figure S1D)**. The separation set obtained from mAb-2E9-treated cells suggests two components of <9 nm (4.9-6.8 nm and 7.2-8.9 nm) (**Figure 2E**; cyan and black dashed lines), consistent with those in the WT-EGFR separation set. In contrast, no separation density at 6.8-9.9 nm is apparent in the set from cells treated with mAb-108, which blocks high-affinity binding, (**Figure 2F**; black dashed line). To assess the robustness of this result, because incompletely resolved components cannot be perfectly separated, we compared the evidence for separations in bootstrap-resampled datasets after pooling the individual components (marginalized probability) (**STAR methods**). This analysis indicates that the absence of separation density in the vicinity of ∼8 nm is robust to finite sampling errors (**Figure 2G**). We therefore assigned the 7.0-8.6 nm peak in the WT-EGFR separation set to the St2St^ect^/Asym^kin^_dimer_. Further validation of this assignment is provided below.

Despite the T766M mutation destabilizing the Asym^kin^_dimer_,^43^ the separation density at ∼8 nm is similar between WT-EGFR and T766M-EGFR (**Figure 2A, 2D)**. We hypothesized that the T766M mutation indirectly underpins formation of the St2St^ect^/Asym^kin^_dimer_ by stabilizing the inactive B2B^ect^/H2H^kin^_dimer_. To test this, WT-EGFR expressing cells were treated with Erlotinib,^60^ a TKI that binds to EGFR’s kinase ATP-binding pocket stabilizing the St2St^ect^/Asym^kin^_dimer_.^59^ Erlotinib treatment was found to recapitulate effects of stabilizing the B2B^ect^/H2H^kin^_dimer_ via the T766M mutation, as reflected by the similarity between the separations of <20 nm associated with Erlotinib-treated WT-EGFR and T766M-EGFR (**Figure 2I, 2D, 2A**). Quantified in **Figure 2J**, both induce a decrease in the 5.8-7.1 nm component, the assignment of which is provided below, and an increase at ∼9-12 nm. We will discuss later that this reflects a conformational change. These results argue that stabilizing either the B2B^ect^/H2H^kin^_dimer_ or the St2St^ect^/Asym^kin^_dimer_ has the same effect on <20 nm separations, and therefore that these dimers interact.

Further evidence of the joint participation of the B2B^ect^/H2H^kin^_dimer_ and St2St^ect^/Asym^kin^_dimer_ in hetero^conf^-oligomer assembly is shown in **Figure 2H**. This includes the increase (decrease) in oligomer size induced by the T766M (ED/RK) mutations combined with the differences in oligomer size associated with mAb-2E9 and mAb-108, which suggest that B2B^ect^/H2H^kin^_dimer_ units and St2St^ect^/Asym^kin^_dimer_ ligand-binding sites are distributed along oligomers. Notably, Erlotinib does not significantly increase oligomer size for reasons discussed later (**Figure 7**). These results assign the role of underpinning oligomer growth to the H2H^kin^_dimer_.

### The H2H^ect^_dimer_/2x^kin^_monomers_ participates in hetero^conf^-oligomer assembly

In a previous study, we proposed a third ligand-free dimer on the cell surface based on a lattice contact in an X-ray structure of the tethered ectodomain monomer (PDB ID:4KRP^61^), in which the monomers are held by ectodomain interactions (H2H^ect^_dimer_). Because the distance between transmembrane domains in the H2H^ect^_dimer_ is too large to facilitate intracellular interactions, we proposed that the H2H^ect^_dimer_ is linked to two non-interacting kinase monomers (H2H^ect^_dimer_/2x^kin^_monomers_)^43^ (**Figure 3A**). This suggestion was supported by the FLImP results associated with ΔC-EGFR, a mutant in which the intracellular domains are deleted. Application of the higher resolution decomposition to that data shows peaks at almost a fixed interval (**Figure 3B**), consistent with the previously proposed homo-oligomers of repeating extracellular H2H^ect^_dimer_ units.^43^ ΔC-EGFR results exemplify the need for mutational studies to assign FLImP separations. Note how the 2^nd^-order peak from the truncated H2H^ect^_dimer_ of ΔC-EGFR appears close to the position assigned to the B2B^ect^/H2H^kin^_dimer_ (**Figure 3B**; red dash line). The presence of an H2H^ect^_dimer_/2x^kin^_monomers_ is further supported by the finding that the 1^st^-order separation of the truncated H2H^ect^_dimer_ of ΔC-EGFR shares position with the shortest component in the separation set for WT-EGFR (**Figure 3B**; cyan dashed line). Taken together, previous and current evidence confirms the presence of H2H^ect^_dimer_/2x^kin^_monomers_ as a third ligand-free dimer on the cell surface, along with the B2B^ect^/H2H^kin^_dimer_ and St2St^ect^/Asym^kin^_dimer_.

**Figure 3:**
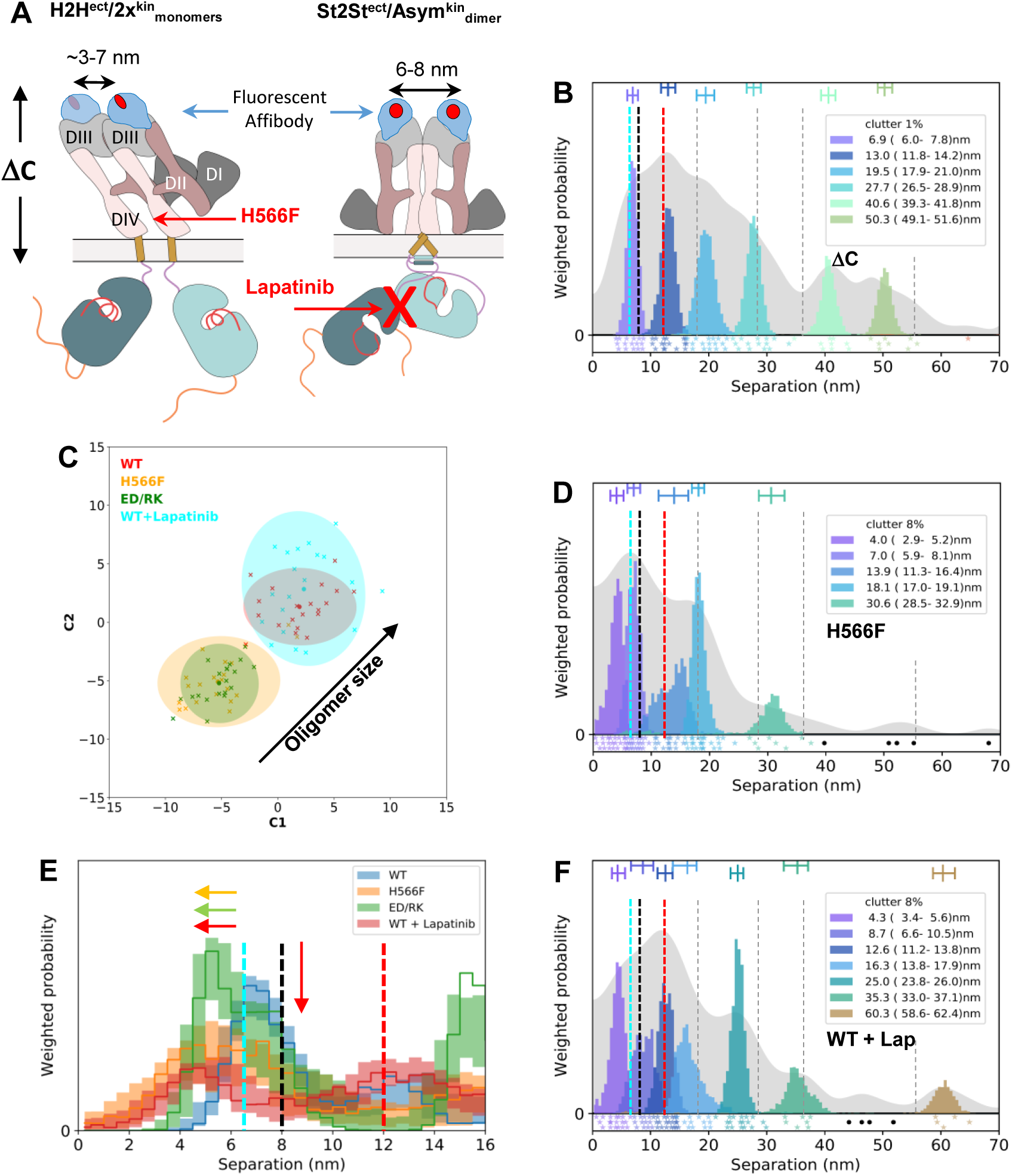
The H2H^ect^_dimer_/2x^kin^monomers participates in hetero^conf^-oligomer assembly. **(A)** (*Left*) Cartoon of labeled full-length H2H^ect^/2x^kin^_monomers_ with two fluorescent anti-EGFR Affibody bound to the two DIII of the ectodomains. The length of the ΔC-EGFR deletion mutant is marked. DIV-binding tether-disruptive mutations interfere with the H2H^ect^_dimer_ conformation. (*Right*) A labeled St2St^ect^/Asym^kin^_dimer_. Lapatinib binds the kinase ATP-binding pocket disrupting the Asym^kin^_dimer_. **(B, D, F)** FLImP analysis as in Fig 2. **(C)** Wasserstein MDS analysis comparing the 21 separation sets (main FLImP set + 20 bootstrap-resampled) of the conditions marked in the plot. Axis labels and 95% confidence ellipses (as in caption Figure 1G). **(E)** Comparisons as in Figure 1(F).

To understand the role of the H2H^ect^_dimer_/2x^kin^_monomers_ we introduced mutations that disrupt the H2H^ect^_dimer_. Our MD simulations suggest that disruption of the tether via two well-understood mutations in DIV of EGFR’s ectodomain, H566F and G546P^62^, disrupt the conformation of the H2H^ect^_dimer_ (**Figure S2A-S2C)**. Revealing a much sought-after biological function for EGFR’s ectodomain tether these mutations induce a shift in the shortest separation component above assigned to the H2H^ect^_dimer_/2x^kin^_monomers_ from a median position of 6.5 nm to 4 nm, which implicates the ectodomain tethered conformation in the formation of H2H^ect^_dimer_/2x^kin^_monomers_ (**Figure 3D, 3E**; blue and orange). Results for G564P are shown in **Figure S3A-S3D**. Note that the component that includes an 8 nm separation, corresponding to the St2St^ect^/Asym^kin^_dimer_, is not significantly decreased by the H566F mutation (**Figure 3E)**. This is consistent with the tether-disrupting mutations not inhibiting phosphorylation^62^ (**Figure S3E**).

Because MD simulations suggest that the WT-EGFR H2H^ect^_dimer_ can explore a separation range of 3-7 nm between the center of mass of the two DIIIs (**Figure S2B)**, the shift in the separation component annotated to the H2H^ect^_dimer_/2x^kin^_monomers_ towards short separations introduced by the ED/RK mutation (**Figure 3E**; blue and green) suggests that inhibiting the B2B^ect^/H2H^kin^_dimer_ changes H2H^ect^_dimer_/2x^kin^_monomers_ conformation. The ED/RK mutations also induce an oligomer size reduction comparable to that induced by the H566F mutation (**Figure 3C**). Together these results suggest that the B2B^ect^/H2H^kin^_dimer_ and H2H^ect^_dimer_/2x^kin^_monomers_ interact to assemble ligand-free hetero^conf^­oligomers.

To investigate whether H2H^ect^_dimer_/2x^kin^_monomers_ interacts with the St2St^ect^/Asym^kin^_dimer_, WT-expressing cells were treated with Lapatinib, a TKI that binds to EGFR’s kinase ATP-binding site breaking the Asym^kin^_dimer_.^63^ As expected, Lapatinib induced a significant reduction around the 8 nm position assigned to the St2St^ect^/Asym^kin^_dimer_ (**Figure 3E**; blue and red). This is accompanied by a shift from a median position of 6.5 nm, the component assigned to H2H^ect^_dimer_/2x^kin^_monomers_, to 4.2 nm (**Figure 3E, 3F**; cyan dashed line), arguing that inhibiting the St2St^ect^/Asym^kin^_dimer_ changes the H2H^ect^_dimer_/2x^kin^_monomers_ conformation and hence the H2H^ect^_dimer_/2x^kin^_monomers_ and St2St^ect^/Asym^kin^_dimer_ interact. We showed above that stabilizing the St2St^ect^/Asym^kin^_dimer_ has the opposite effect stabilizing the H2H^ect^_dimer_/2x^kin^_monomers_ (**Figure 2J)**. The appearance of a ∼25 nm peak suggests that Lapatinib induces an oligomer conformational change (**Figure 3F**). Interestingly, Lapatinib does not decrease oligomer size (**Figure 3C**). This will be discussed later with more data (**Figure 7**).

### Ligand-free dimers interact via transmembrane contacts

The ligand-free dimer geometries suggest that hetero^conf^-oligomer assembly might be mediated by transmembrane interactions. Therefore, we next considered the *Lzip* transmembrane dimer named after its leucine zipper-like interactions (V^635^xxL^638^xxxL^642^)^51^ (**Figure 4A)**. *Lzip* dimer residues are distinct from those forming the other transmembrane dimers and could therefore in principle mediate interactions between the H2H^ect^_dimer_/2x^kin^_monomers_ and the other ligand-free dimers (**Figure 4B, S3F**). To investigate this possibility, we mutated all three *Lzip* dimer transmembrane helix amino acids (V635, L638, L642) to either serine or alanine, named *Lzip*3S and *Lzip*3A, respectively. Based on previous literature, *Lzip*3S mutations would be expected to strongly inhibit the *Lzip* interaction, unlike the more conservative *Lzip*3A.^64^

**Figure 4:**
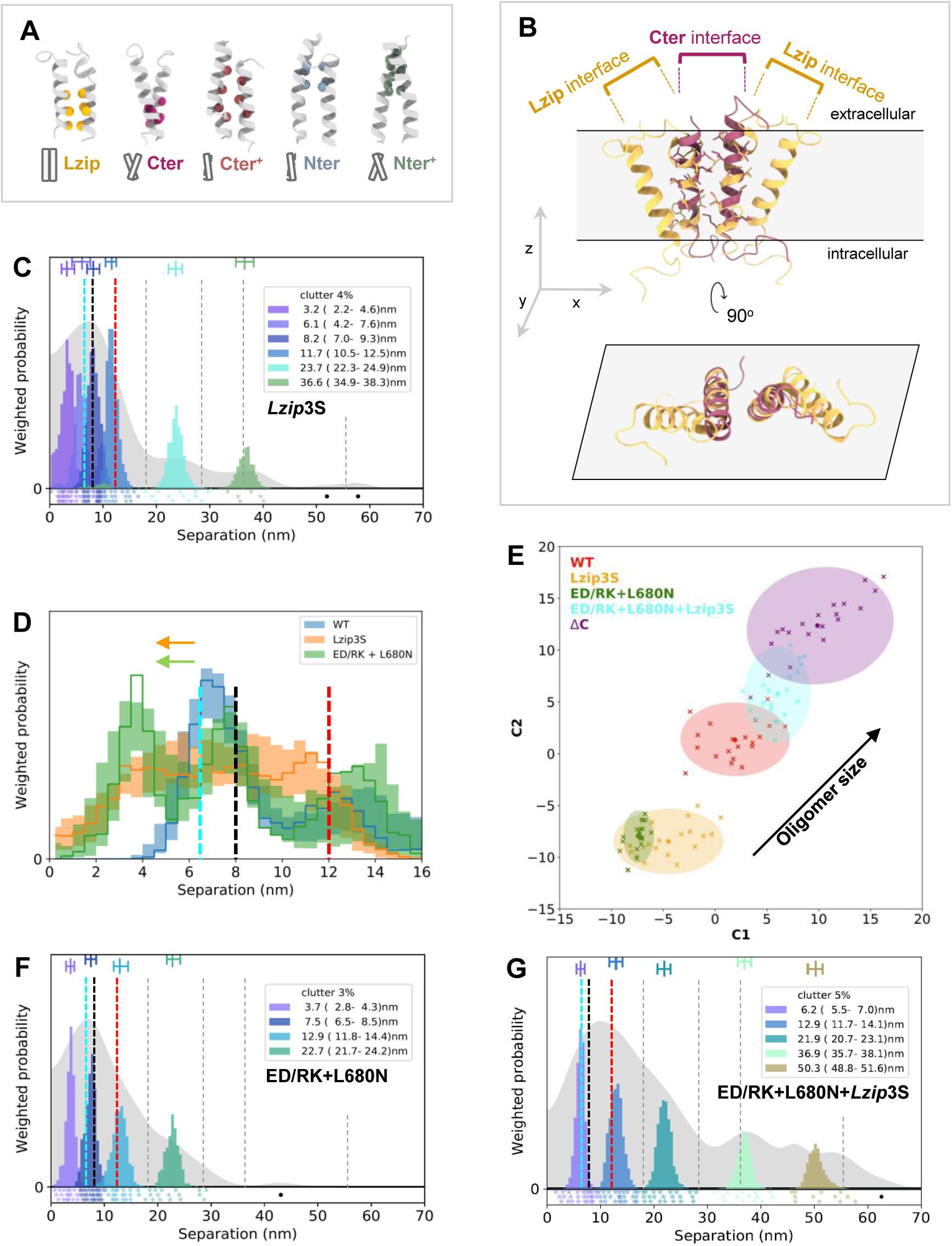
Ligand-free dimers interact via transversal transmembrane contacts. **(A)** Transmembrane dimers simulated by Lelimousin et al.^51^ *Lzip* is the only left-hand dimer. *Nter*^+^ is linked to ligand-bound dimers.^51^ *Cter*^+^, *Nter* and *Cter* might be associated to ligand-free dimers^45^. **(B)** (*Top*) A speculative tetramer formed by two monomers interacting through two *Lzip* contacts and one *Cter* interface. (*Bottom*) Orthogonal projection on xy-plane. **(C, F, G)** FLImP analysis as in Fig 2. **(D)** Comparisons as in Figure 1(F). **(E)** Wasserstein MDS analysis comparing the 21 separation sets (main FLImP set + 20 bootstrap-resampled) of the conditions marked in the plot. Axis labels and 95% confidence ellipses as in caption Figure 1G.

We found that the short separations in the *Lzip3S*-EGFR set are poorly resolved (**Figure 4C**). Notable effects include increased separation density at 2.2-4.6 nm (**Figure 4C, 4D)**, suggesting the conformation of H2H^ect^_dimer_/2x^kin^_monomers_ has changed, alongside a reduction in oligomer size (**Figure 4E**). The *Lzip*3A mutations did not show these effects (**Figure S3G-S3I)**. Together, these results indicate that *Lzip* contacts are involved in hetero^conf^-oligomer assembly.

To evaluate the consistency of the FLImP results, we reasoned that simultaneously inhibiting the B2B^ect^/H2H^kin^_dimer_ and St2St^ect^/Asym^kin^_dimer_, thus all intra-dimer intracellular interactions, together with *Lzip* contacts should recapitulate the results for ΔC-EGFR. This hypothesis was evaluated in two stages. First, by combining the ED/RK mutations with L680N, a kinase N-lobe mutation that prevents the kinase domain from acting as receiver,^56^ thus inhibiting the St2St^ect^/Asym^kin^_dimer_ and phosphorylation. Then the *Lzip*3S mutations were added. We found that the ED/RK+L680N mutations induce a shift in the peak associated to the H2H^ect^_dimer_/2x^kin^_monomers_ and an oligomer size reduction comparable to the *Lzip*3S mutations alone (**Figure 4E-4F)**. Reassuringly, combining ED/RK+L680N+*Lzip*3S mutations recapitulated the results for ΔC-EGFR (**Figure 4G, 4E**), e.g., the loss of the 2.8-4.3 nm peak in **Figure 4F**, pseudo-periodic separations, and the oligomer size increase, suggesting that *Lzip* contacts inhibit the formation of the homo-oligomers of repeating extracellular H2H^ect^_dimer_ interfaces.

### The extracellular structure of ligand-free hetero^conf^-oligomers

Based on the above data, we constructed a model of the orthogonal projection on the cell surface of the hetero^conf^-oligomer based on the known shape and dimensions of the three ligand-free dimers (**Figure 1B, S4A-4C)**. The *Lzip* interface is related to the transmembrane dimers in the B2B^ect^/H2H^kin^_dimer_ and St2St^ect^/Asym^kin^_dimer_ by a rotation of the helix along the long axis^51^ (**Figure 4B, S3F**). Attempts to join an H2H^ect^_dimer_/2x^kin^_monomers_ and a St2St^ect^/Asym^kin^_dimer_ via *Lzip* contacts led to steric clashes (**Figure S4**). In contrast an H2H^ect^_dimer_/2x^kin^_monomer_ and a B2B^ect^/H2H^kin^_dimer_ could form a tetramer (**Figure 5A)**. Remarkably, in this tetramer the *Lzip* interface positions one of the ectodomains of the H2H^ect^_dimer_/2x^kin^_monomers_ at an orientation that can facilitate the St2St^ect^/Asym^kin^_dimer_.

**Figure 5:**
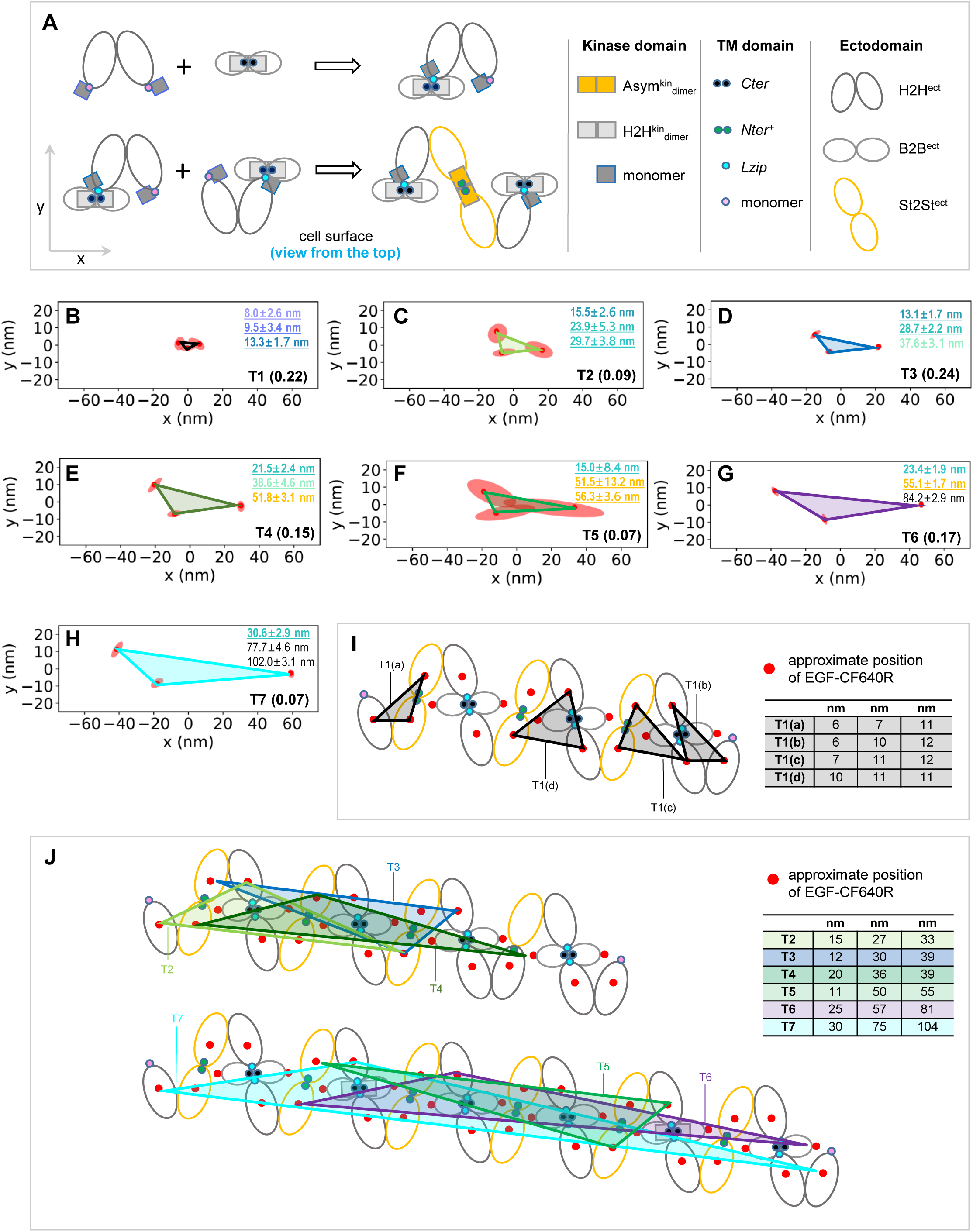
The extracellular structure of ligand-free hetero^conf^-oligomers. **(A)** (*Top Left*) Tetramer assembled by an H2H^ect^*/*2x^kin^_monomers_ and a B2B^ect^/H2H^kin^_dimer_ via an *Lzip* interface. (*Bottom left*) Two tetramers form a St2St^ect^/Asym^kin^_dimer_. (*Right*) Annotated interfaces in the model. More details in **Figure S4.** (**B-H**) Optimally grouped seven distinct triangles from 2D FLImP separations determined between EGF-CF640 probes bound to cell surface T766M-EGFR (**Figure S5**). The side lengths of each triangle annotated in the colors used in 1D FLImP decompositions when found in the separation sets of either T766M-EGFR or WT-EGFR, and colored and underlined if found in both. (Note that 2D FLImP considers larger separations than 1D FLImP). The abundance of each triangle is also annotated (*Bottom right*, black). **(I)** Ligand-free hetero^conf^-oligomer model extended as described in (A). The approximate positions where EGF-CF640R would bind DIII of the ectodomains are marked (red circles). Four versions of the experimentally-optimized triangle 1 (T1a-T1d) are superimposed. (*Inset*) Table of the lengths of the superimposed triangle sides. **(J)** (*Top*) Ligand-free hetero^conf^-oligomer model with the triangle groups T2-T4 (C-E) superimposed. (*Bottom*) Hetero^conf^-oligomer model with the largest triangles T5-T7 (F-H) superimposed. (*Inset*) Table of the side lengths of the superimposed triangles.

To test this model and its relevance to T766M-EGFR, we implemented a 2D version of FLImP that reports triangular arrangements between probes bound to EGFR structures (**STAR methods**). The 3-fold higher probe concentration required for 2D FLImP was better suited to the less sticky EGF-CF640R derivative, as Affibody-CF640R began at this higher concentration began to show signs of non-specific binding on the glass supporting the cells.^65^

We probed cells expressing T766M-EGFR with EGF-CF640R after chemical fixation to avoid ligand-induced conformational changes, as discussed above. The resulting 2D FLImP triangle dataset was optimally grouped into seven distinct triangles (**Figure 5B-5H, S5)**. Reassuringly, the 1D separations in the triangles are found as components decomposed by 1D FLImP for T766M-EGFR and WT-EGFR (**Figure 2D, 2A)**, arguing that T766M-EGFR and WT-EGFR share hetero^conf^-oligomer structure.

To evaluate whether the 2D FLImP data supports the proposed model, the model was expanded to the size required by the triangles and the positions we expected for EGF-CF640R probes bound to EGFR’s ectodomain DIII marked (**Figure 5I)**. We found that the smallest triangle (T1) accounts, within errors, for four triangular probe motifs in the model (**Figure 5I**). Triangles T2-T7 each account for one motif (**Figure 5J)**. To further validate the applicability of the hetero^conf^ oligomers to T766M-EGFR, we verified that combining the T766M mutation with the tether-disrupting H566F mutation or the *Lzip*3S mutations disrupts the hetero^conf^-oligomers (**Figure S6A-6D**). This, together with the excellent results of superimposing the triangles from 2D FLImP data collected from CHO cells expressing T766M-EGFR indicate that the model is an accurate representation of the extracellular portion of the ligand-free T766M-EGFR hetero^conf^-oligomers.

### The St2St^ect^/Asym^kin^_dimer_ is buttressed by a symmetric kinase interface

The model predicts that inhibiting the B2B^ect^/H2H^kin^_dimer_ would result in the tetramer in **Figure 6A**. Inhibiting the *Lzip* interface should have the same effect, but separations found when the B2B^ect^/H2H^kin^_dimer_ is inhibited by the ED/RK mutations are inconsistent with those found when *Lzip* contacts are inhibited via *Lzip*3S mutations (**Figure 6B**). This hinted at the possibility that we had not yet accounted for all interactions.

**Figure 6:**
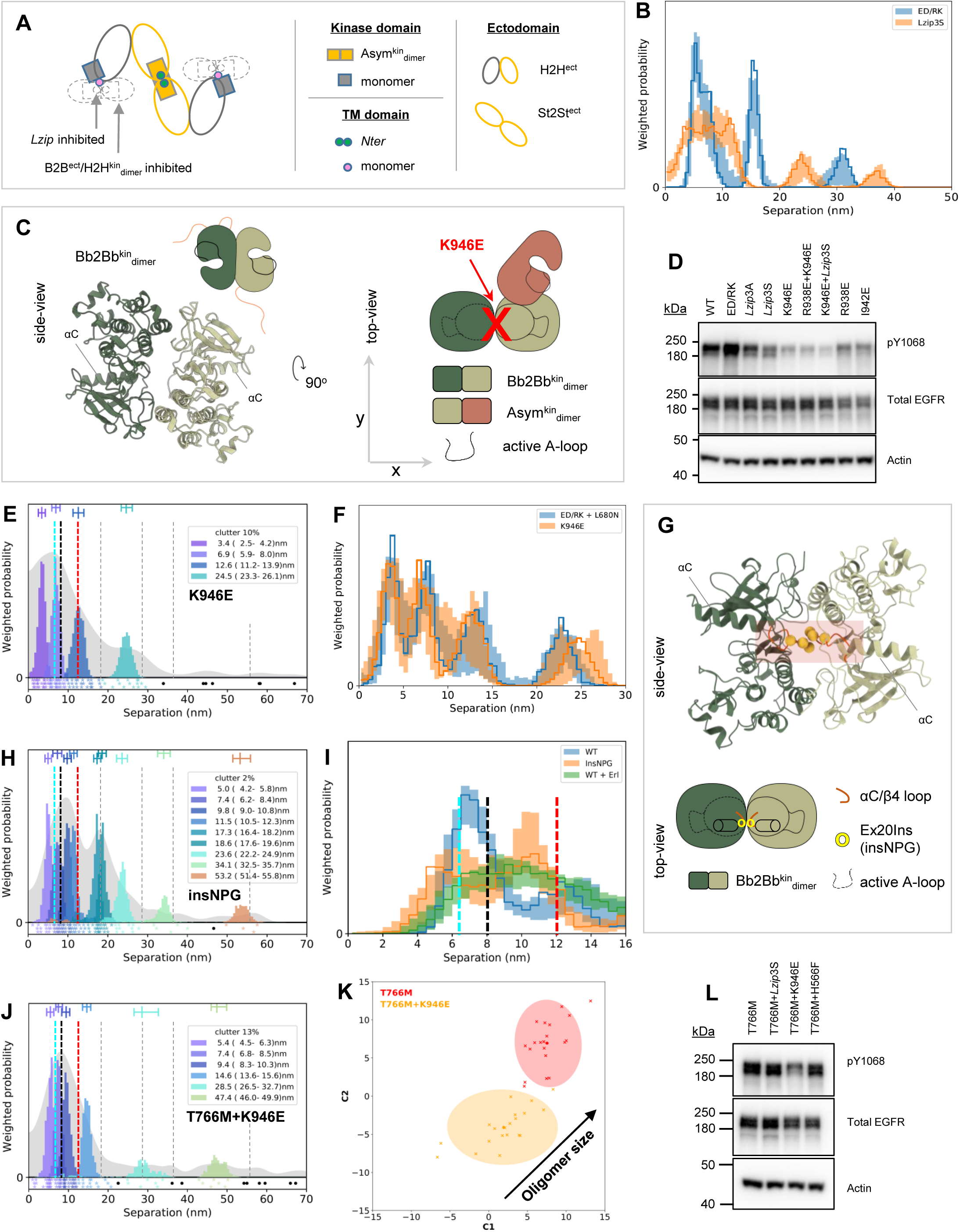
A Bb2Bb^kin^_dimer_ underpins activation in ligand-free oligomers. **(A)** (*Left*) Tetramer that would form if the B2B^ect^/H2H^kin^_dimer_ or *Lzip* contacts are inhibited. (*Right*) Interfaces annotated. **(B, F, I)** Comparisons as in Figure 1(F). **(C)** Structure and cartoon model of the Bb2Bb^kin^_interface_ formed mainly through N-to-C lobe interactions (PDB ID 3VJO).^52^ A kinase monomer can dock in the Bb2Bb^kin^_interface_ to form an Asym^kin^_dimer_, as shown. The position of the A-loop in the active configuration of the Asym^kin^_dimer_ is marked. The K946E mutation breaks the Bb2Bb^kin^_interface_ interface (**Figure S7A**). **(D)** Phosphorylation of the annotated mutant-EGFR in the absence of ligand in transfected CHO cells. **(E, H, J)** FLImP analysis as in Fig 2. **(G)** Side and top-view of the Bb2Bb^kin^_dimer_ (described in^52^) formed mainly through N-to-C lobe interactions (PDB ID: 3VJO).^52^ The inserted residues of the Ex20Ins mutation, which lies on the αC/β4 loop, are shown in spheres. **(K)** Wasserstein MDS analysis comparing the 21 separation sets (main FLImP set + 20 bootstrap-resampled) of the conditions marked in the plot. Axis labels and 95% confidence ellipses (as in caption Figure 1G). **(L)** Phosphorylation of the annotated mutant-EGFR in the absence of ligand in transfected CHO cells.

We considered a functionally orphan symmetric backbone-to-backbone kinase interface (Bb2Bb^kin^_interface_), revealed by X-ray crystallography^52^ (**Figure 6C**). Because a kinase monomer could dock into a Bb2Bb^kin^_dimer_ to form an Asym^kin^_dimer_, we speculated that the Bb2Bb^kin^_interface_ might be involved in oligomer assembly. Giving credence to this possibility, two non-naturally occurring charge-reversal R838E and K946E mutations, which compromise the Bb2Bb^kin^_dimer_, as reported by MD simulations (**Figure S7A-S7B**), decrease receptor phosphorylation (**Figure 6D**).

Remarkably, we found that the separation set associated with K946E-EGFR is almost indistinguishable from that of ED/RK+L680N-EGFR (**Figure 6E, 6F)**. This reveals that disrupting the Bb2Bb^kin^_interface_ via the K946E mutation recapitulates the joint effect of inhibiting both the St2St^ect^/Asym^kin^_dimer_ and the B2B^ect^/H2H^kin^_dimer_ via the combined ED/RK+L680N mutations. Given **Figure 6C** and the reduction in phosphorylation (**Figure 6D)**, these results suggest that the Bb2Bb^kin^_interface_ underpins the St2St^ect^/Asym^kin^_dimer_, and thereby its interaction with the B2B^ect^/H2H^kin^_dimer_ in the hetero^conf^-oligomers.

If the above interpretation is correct, one would expect that stabilizing the Bb2Bb^kin^_interface_ should mirror the effects of stabilizing the St2St^ect^/Asym^kin^_dimer_. We conjectured that EGFR exon 20 insertions (Ex20Ins) might increase the number of contacts between the kinase domains of the Bb2Bb^kin^_interface_, stabilizing that interface (**Figure 6G**). This was supported by MD simulations of the WT and D770-N771insNPG (insNPG), a mutant chosen because structural data is available^66^ (**Figure S7C-7D**). Furthermore, over the course of the simulation, one of the monomers of the insNPG-EGFR in the Bb2Bb^kin^_interface_ is stabilized in the active conformation characteristic of the Asym^kin^_dimer_, and the other samples it as well. In support of the notion that the Bb2Bb^kin^_interface_ underpins St2St^ect^/Asym^kin^_dimer_ stability, the separation set for insNPG-EGFR recapitulates the effects of the Erlotinib treatment in WT-EGFR in short and long separations (**Figure 6H, 6I**, **S6E**).

Adding the K946E mutation to T766M-EGFR also reduces oligomer size and decreases receptor phosphorylation (**Figure 6J-6L)**, in turn implicating the Bb2Bb^kin^_interface_ in the assembly and phosphorylation of T766M-EGFR hetero^conf^-oligomers.

### A mechanism of T766M-induced oligomer activation

After incorporating the Bb2Bb^kin^_interface_ (**Figure 7A**), another prediction is that disrupting *Lzip* contacts should have an analogous effect on phosphorylation to inhibiting the B2B^ect^/H2H^kin^_dimer_ via the ED/RK mutations, but only the latter increases phosphorylation (**Figure 6D**). We speculated this might be explained if the B2B^ect^/H2H^kin^_dimer_ sequestered kinase monomers thereby competing with the formation of the Bb2Bb^kin^_interface_ and destabilizing the Asym^kin^_dimer_ (**Figure 7B**). In the crystal lattice of the activator-impaired V924R-H2H^kin^_dimer_^53^ we noticed a side-to-side interface (S2S^kin^_interface_) that could in principle play such a role (**Figure 7C**).

**Figure 7:**
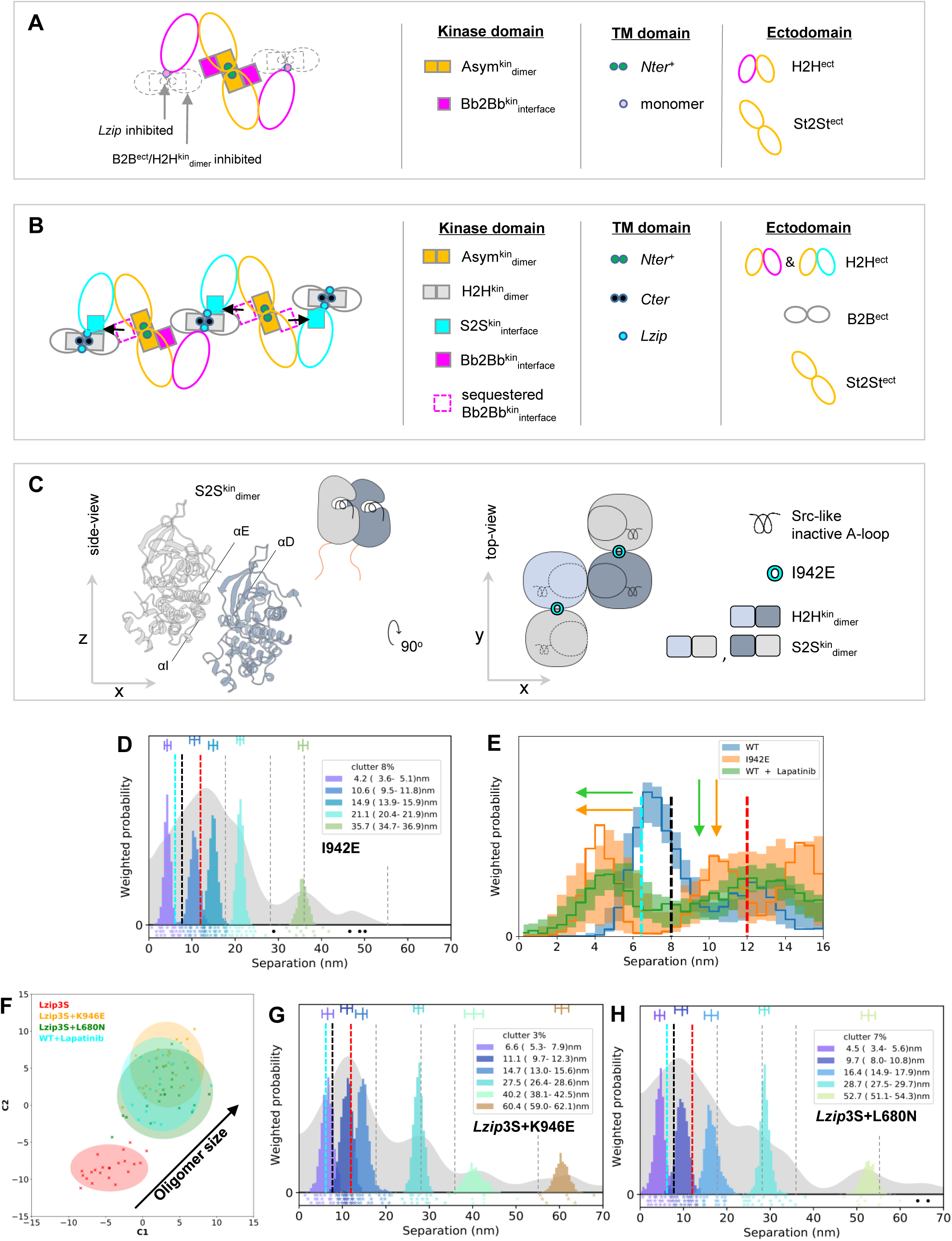
An autoinhibitory S2S^kin^_dimer_ promotes hetero^conf^-oligomer growth. **(A)** (*Left*) Tetramer that would form if the B2B^ect^/H2H^kin^_dimer_ or *Lzip* contacts are inhibited, incorporating two Bb2Bb^kin^_interfaces_ buttressing an Asym^kin^_dimer_. (*Right*) Interfaces annotated. **(B)** (*Left*) Model of the ligand-free hetero^conf^-oligomer incorporating three S2S^kin^_interfaces_. (*Right*) Assembling interfaces annotated. **(C)** (*Left*) Side-view of the S2S^kin^_dimer_ maintained mainly through the β2-sheet and αD-helix of the one monomer and the αE- and αΙ-helices of the other monomer (PDB ID: 5CNO)^53^. The position of the A-loop in the inactive configuration is marked. (*Right*) Cartoon representation of the top-view of an H2H^kin^_dimer_ flanked by two monomers via S2S^kin^_interfaces_. **(D, G, H**) FLImP analysis as in Fig 2. **(E)** Comparisons as in Figure 1F. **(F)** Wasserstein MDS analysis comparing the 21 separation sets (main FLImP set + 20 bootstrap-resampled) of the conditions marked in the plot. Axis labels and 95% confidence ellipses (as in caption Figure 1G).

We previously reported that the non-naturally occurring I942E mutation inhibits ligand-bound complex formation.^43^ Here we show that I942E decreases ligand-free EGFR phosphorylation (**Figure 6D**). The I942 residue lies at the heart of the S2S^kin^_interface_, and modelling of the WT and I942E mutant suggested that the latter stabilizes the S2S^kin^_interface_ (**Figure S7E)**. Supporting the notion that the S2S^kin^_interface_ plays an autoinhibitory role, the separation set associated with I942E-EGFR is consistent with the results obtained for WT-EGFR-expressing cells treated with Lapatinib, which inhibits the Asym^kin^_dimer_ (**Figure 7D-7E**). These results indicate that S2S^kin^_interface_stabilization via the I942E mutation also inhibits the St2St^ect^/Asym^kin^_dimer_. Combining the T766M and I942E mutations support this notion (**Figure S6F-S6G**).

If the B2B^ect^/H2H^kin^_dimer_ sequestered some kinase monomers via the S2S^kin^_interface_ preventing these from reinforcing the St2St^ect^/Asym^kin^_dimer_ via the Bb2Bb^kin^_interface_, then inhibiting the Bb2Bb^kin^_interface_ should make more kinase monomers available to stabilize the B2B^ect^/H2H^kin^_dimer_ via the S2S^kin^_interface_, thereby growing larger oligomers. This was observed for the T766M mutation that stabilizes directly B2B^ect^/H2H^kin^_dimer_ (**Figure 2H)**. To test this possibility, we need to inhibit the Bb2Bb^kin^_interface_ whilst preserving the St2St^ect^/Asym^kin^_dimer_ and B2B^ect^/H2H^kin^_dimer_ so they can compete for kinase monomers. However, when the Bb2Bb^kin^_interface_-inhibitory K946E mutation is introduced in WT-EGFR, the mutation also disassembles the St2St^ect^/Asym^kin^_dimer_ and B2B^ect^/H2H^kin^_dimer_ (**Figure 6G**). In contrast, partially disassembled *Lzip*3S-EGFR oligomers preserve their phosphorylation (**Figure 6D**) and some density around 12 nm (**Figure 6B**), suggesting that *Lzip*3S-EGFR oligomers retain some interacting St2St^ect^/Asym^kin^_dimer_ and B2B^ect^/H2H^kin^_dimer_ units. Thus, we reasoned that adding K946E to *Lzip*3S mutations should stabilize the B2B^ect^/H2H^kin^_dimer_ and increase oligomer size. The results are consistent with this notion (**Figure 7F**). We also found that the separation sets of *Lzip*3S+K946E and *Lzip*3S+L680N are similar below ∼35 nm (**Figure 7G-7H)**, suggesting that inhibiting the St2St^ect^/Asym^kin^_dimer_ and the Bb2Bb^kin^_interface_ are almost equivalent, and explaining why disrupting the St2St^ect^/Asym^kin^_dimer_, either directly via Lapatinib or indirectly via the I942E mutation, does not decrease oligomer size, as both stabilize the S2S^kin^_interface_ and thereby the H2H^kin^_dimer_ counterbalancing the effect of disrupting the St2St^ect^/Asym^kin^_dimer_ (**Figure 7F**).

Results, therefore, show that destabilizing the St2St^ect^/Asym^kin^_dimer_, either directly by the L680N mutation or indirectly by inhibiting the Bb2Bb^kin^_interface_, increases oligomer size. In WT-EGFR this depends on the S2S^kin^_interface_ at the expense of phosphorylation. The T766M mutation directly stabilizes the B2B^ect^/H2H^kin^_dimer_, thus promoting the formation of larger oligomers without detrimental effects on phosphorylation, thereby conferring a cellular growth advantage to mutated cells.

### Tumor growth depends on hetero^conf^-oligomerisation

To test the relevance of the proposed ligand-free oligomerization model *in vivo*, we carried out cellular transformation assays using the IL3-dependent murine lymphoid Ba/F3 cell system with the aim of using these cells to establish tumor xenografts. Ba/F3 cells fail to survive and multiply in the absence of IL3^67, 68^ but this phenotype can be rescued by the ectopic expression of a constitutively active receptor tyrosine kinase such as T766M-EGFR, which allows survival signaling in the absence of IL3^67^.

We generated Ba/F3 cell lines stably expressing WT-EGFR (Ba/F3+WT), T766M-EGFR (Ba/F3+T766M), and EGFR mutants at near equal levels using the PiggyBac system and tested the ability of the transformed cells to grow in the absence of IL3 (**Figure S8A**). To minimize mice number, from mutations that disrupt hetero^conf^-oligomer structure and impair the ability of Ba/F3 cells to grow we focused on the T766M+H566F and omitted the equivalent T766M+*Lzip*3S (**Figure S6A-S6D**). We also chose T766M+K946E over T766M+I942E because unlike the I942E mutation, K946E does not interfere with the ligand-bound state (**Figure S8B**), and therefore potential drugs should carry fewer side effects.

Next, we compared the ability of these Ba/F3 cell lines to establish and grow tumors. Notably, all these cell lines also stably expressed GFP, exploited to image growing tumors. We found a lag-time prior to the onset of palpable tumors of around 20 days in all cohorts, after which the tumors formed in animals that had received Ba/F3+T766M and Ba/F3+T766M+H566F cells and with some additional delay in the Ba/F3+T766M+K946E cohort (tumor growth measurements by calipers; **Figure 8A**). Importantly, mirroring the growth pattern observed *in vitro*, Ba/F3+T766M tumors grew best throughout, followed by the double mutant tumors in the order Ba/F3+T766M+H566F > Ba/F3+T766M+K946E, while wild-type EGFR tumors did not establish or grow. The lack of tumor cell survival in the Ba/F3+WT cohort was already strongly suggested by day 21 when we observed no GFP reporter signals at the injection site (**Figure S8C-S8D**). The experimental endpoint was on day 41 when we imaged all animals by IVIS (**Figure 8B-8C)** to determine tumor fluorescence as an independent measure of tumor growth between cohorts. We found significant differences in fluorescence signals between groups in line with caliper measurements (*cf.* **Figure 8A**) except for Ba/F3+T766M *vs.* Ba/F3+T766M+H566F tumors. The latter was not surprising and demonstrated the limitations of epifluorescence imaging, which did not reflect accurately the signal differences between larger tumors due to limited tissue penetration and absorption within thicker tissues. Harvested tumors were first qualitatively imaged under daylight and by fluorescence (IVIS; **Figure 8D**), and then weighed (**Figure 8E)** upon which the significant growth differences between the Ba/F3+T766M *vs.* Ba/F3+T766M+H566F tumors were evident as were the other differences already seen *in vivo* in **Figure 8A**. Moreover, we subjected harvested tumor tissues to histology to demonstrate tumor morphology (by H&E/**Figure 8F**) and pan-EGFR expression (by anti-EGFR staining (**Figure 8G** *vs.* staining control/**Figure S8F**). All tumors were positive for EGFR with plasma membrane localization clearly visible in stained tissues (**Figure 8G**).

**Figure 8:**
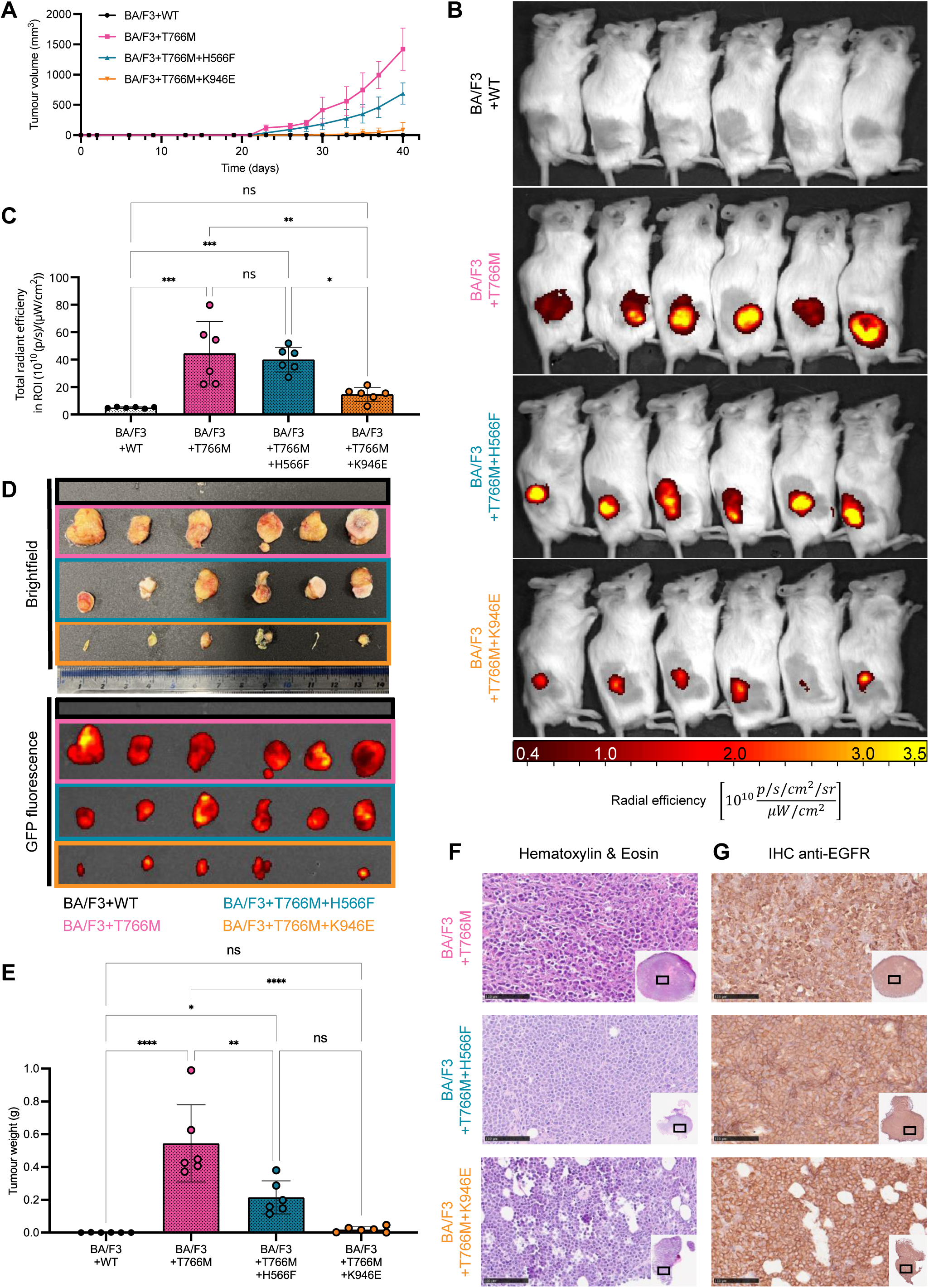
Tumors established from Ba/F3 cell lines with wild-type or indicated mutant EGFR showed different growth behavior in vivo. **(A)** NSG mice were used to establish tumors in their flanks (**STAR Methods)** and tumor establishment and cumulative growth was monitored using calipers. N=6 animals per cohort with error bars representing standard deviation (SD). Differences were tested for significance using 2-way ANOVA (alpha=0.05 and Tukey’s multiple comparison correction). **(B)** IVIS imaging of all animals from (A) at the experimental endpoint (day 41) showing obtained epi­fluorescence signals from animal tumors. **(C)** GFP fluorescence signal quantification of tumors corresponding to animals in (B). Statistical analysis by 1-way ANOVA with Tukey’s multiple comparison correction; error bars are SD. **(D)** Harvested tumors from those animals with established tumors were photographed under *(top)* daylight and *(bottom)* blue light illumination to detect fluorescence from GFP. No tumors had established in the cohort that had received Ba/F3+WT cells. **(E)** Tumor weights from harvested tumors. Statistical analysis by 1-way ANOVA with Tukey’s multiple comparison correction; error bars are SD. **(F)** Hematoxylin/eosin staining of tumors that did establish; scale bars are 100μm. **(G)** Immunohistochemistry (IHC) staining using a pan anti-EGFR antibody; corresponding background control staining is shown in Figure S8F. Scale bars are 100 μm.

Notably, the T766M+H566F mutant, which does not reduce basal phosphorylation compared with T766M alone or with T766M+K946E **(Figure 6L)**, also handicaps tumor growth, suggesting that oligomerization per se is conveying a growth advantage. An explanation is that T766M+H566F expressing Ba/F3 cells are unable to phosphorylate and activate AKT signaling to the same extent as T766M (**Figure S8G**).

## Discussion

We combined super-resolution FLImP imaging with *in silico* modelling and mutagenesis to identify the interfaces that assemble ligand-free oligomers. The <3 nm resolution achieved allowed us to determine the fingerprints of the known ligand-free dimer conformers and, crucially, show how interactions between these dimers involving *Lzip* transmembrane contacts promote the assembly of ligand-free hetero^conf^-oligomers. From this knowledge and the known shapes of the ligand-free dimers we derived a structural model of ligand-free hetero^conf^-oligomerization. The extracellular part of this model was thereafter validated by implementing a 2D version of the FLImP method.

The tethered conformation of EGFR’s ectodomain appeared during the evolution of vertebrates,^69^ and was suggested to have evolved to prevent crosstalk between the different EGFR homologs in the vertebrate EGFR family.^70^ Our MD simulations and FLImP results propose a new biological role for the tethered conformation, which is to regulate the formation of the H2H^ect^_dimer_, and through it, hetero^conf^-oligomer size, receptor activation, and, in the pathological context of T766M-EGFR, tumor formation.

Comparing model and data revealed that a previously orphan Bb2Bb^kin^_interface_ reported by X-ray crystallography^52^ plays a regulatory role in ligand-free hetero^conf^-oligomerization and activation. The role of the Bb2Bb^kin^_interface_ is to buttress the Asym^kin^_dimer_, leading to activation once the B2B^ect^/H2H^kin^_dimer_ cantilevers H2H^ect^_dimer_/2x^kin^_monomers_ into position to form the St2St^ect^/Asym^kin^_dimer_. This mechanism explains how the Asym^kin^_dimer_ can form in the absence of ligand-induced conformational changes that are typically required to overcome the activation barrier associated with the formation of the asymmetric interface between activator and receiver kinases.^47^ Importantly, our work shows that Ex20Ins mutations stabilize the Bb2Bb^kin^_interface_, providing a breakthrough in our understanding of these mutations, which so far had been studied only at the monomeric level.

Comparing models and data also suggested a second S2S^kin^_interface_ that we identified from a crystal contact in a structure of the H2H^kin^_dimer_.^56^ This interface allows the H2H^kin^_dimer_ to hijack kinase monomers, preventing formation of the stimulatory Bb2Bb^kin^_interface_, and down-regulating activation. The S2S^kin^_interface_ stabilizes the H2H^kin^_dimer_, providing an explanation for its autoinhibitory role. We propose that the regulation of the activation of ligand-free hetero^conf^-oligomers rests on balancing the interactions between the Asym^kin^_dimer_ and the H2H^kin^_dimer_ with the two newfound kinase interfaces.

Beyond its autoinhibitory role, the H2H^kin^_dimer_ also acts as a scaffold promoting the formation of larger hetero^conf^-oligomers. Thus, by stabilizing the H2H^kin^_dimer_, the T766M mutation indirectly underpins the formation of Asym^kin^_dimer_ units without the need to hijack monomers from Bb2Bb^kin^_interface_, which can thereby still buttress the Asym^kin^_dimer_ and amplify cell growth. This conveys a significant growth signaling advantage to the T766M receptor mutant.

We tested the effect of breaking the ligand-free hetero^conf^-oligomers *in vivo* on T766M-EGFR dependent cell growth by disrupting the tethered conformation and inhibiting the Bb2Bb^kin^_interface_. Disrupting the ectodomain tether associated with the H2H^ect^, which decreases oligomer size, reduces the size of T766M-EGFR driven tumors by half. Excitingly, disrupting the Bb2Bb^kin^_interface_, which has a deleterious effect on both oligomer size and phosphorylation, almost abolishes tumor growth. These results suggest that, counterintuitively, drugs that interfere with the tether, and drugs that can inhibit the Bb2Bb^kin^_interface_ might have an application in the treatment of NSCLC tumors in which the T766M mutation is dominant. Furthermore, these interfaces are away from the ATP-binding pocket mutational hotspot and inhibiting the Bb2Bb^kin^_interface_ has no detectable effect on ligand-bound dimerization, so one could in principle expect no adverse side effects derived from interfering with the ligand-bound state.

Targeting protein-protein interactions is a new direction in treating diseases and an essential strategy for the development of novel drugs.^71^ Given the low long-term efficacy of current treatments for NSCLC, the new structural understanding from this work of how T766M-EGFR amplifies cell growth suggests a possible route for more effective therapies. Notably, mutant and WT-EGFR share hetero^conf^-oligomer structure. It would be interesting to find out whether these principles apply to cancers driven by EGFR overexpression which currently have no therapy.

## Acknowledgements

We thank Drs Esther Garcia-Gonzalez, Jana Harizanova, Chris Tynan, Michael Hirsch, Jianguo Rao and Michalis Vrettas for technical support. We also thank Michael Hirsch for comments on the manuscript. This work has been funded by grant Ref: ST/S000682/1 from the Science and Technology Facilities Council UK and by a PhD studentship to RCHM, which was jointly funded by King’s College London and The University of Hong Kong. MLM-F, DJR, and BMD are grateful for significant computing resources and support provided by STFC Scientific Computing Department’s SCARF cluster and its Data Services, Research Infrastructure and Cloud Operations Groups, with funding from STFC’s Ada Lovelace Centre and IRIS eInfrastructure consortium. FLG and IG acknowledge the Swiss National Science Foundation and Bridge for financial support (project number: 200021_204795 and 40B2-0_203628). FLG and IG also acknowledge the Swiss National Supercomputing Centre (CSCS) for large supercomputer time allocations, project IDs: s1107, s1169, s1228.

## AUTHOR CONTRIBUTIONS

RSI, SRN, IG, BMD, and SKR contributed equally to this work. Conceptualization, RSI, IG, SKR, SRN, LCZ-D, PJP, DJR, BMD, DTC, FLG, and MLM-F; molecular biology, RSI and SKR; in vitro data acquisition, RSI, SRN, SKR, LCZ-D; FLImP automation, DJR, BMD, SRN; algorithm development and data analysis, DJR, BMD; MD simulations, IG and FLG; *in vivo* work RCHM, GOF, RSI, and SKR; writing – original draft, MLM-F; writing – review & editing, IG, LCZ-D, RSI, SRN, DJR, BMD, GOF, PJP, SKR, DTC, FLG, MLM-F; visualization, SRN, LZN-D, BMD, DJR, SKR, DTC, IG, RCHM, RSI; funding acquisition MLM-F, DJR, BMD, and FLG.

## STAR METHODS

### RESOURCE AVAILABILITY

#### Lead contact

Further information and requests for reagents and resources should be directed and will be fulfilled by the lead contact, Marisa Martin-Fernandez.

#### Materials availability

Reagents generated in this study will be made available on request, but we may require a payment and/or a completed Materials Transfer Agreement if there is potential for commercial application.

#### Data and code availability

- All data reported in this paper will be shared by the lead contact upon request.
- Original code will be made available on request, but we may require a completed Materials Transfer Agreement if there is potential for commercial application.
- Any additional information required to reanalyze the data reported in this paper is available from the lead contact upon request.

### EXPERIMENTAL MODEL AND SUBJECT DETAILS

#### Cell culture

All reagents for cell culture were purchased from ThermoFisher Scientific unless stated otherwise. All cells were grown at 37°C in a humidified atmosphere with 5% CO _2_ in air. Cell lines were verified negative for mycoplasma before use and tested routinely.

##### mAb-108 expressing hybridoma cells

mAb108 expressing hybridoma cells were purchased from ATCC (HB-9764) and cultured in phenol-red free high glucose DMEM media supplemented with 10% Fetal Bovine Serum (FBS) and 1 mM sodium pyruvate.

##### Generation of EGFR CHO stable cell lines

Chinese Hamster Ovary (CHO) cells expressing one of the following were generated: WT-EGFR, ED/RK EGFR, T766M EGFR, T766M + K946E EGFR or T766M + I942E. CHO cells were provided by PJP and were cultured in phenol-red free DMEM/F12 media supplemented with 10% FBS and 1X Penicillin/Streptomycin.

CHO cells were seeded at a density of 2 x 10^5^ cells per well in a 6-well dish, and transfected the following day with a mix of 0.5 μg of Super Piggybac Transposase expression vector and 5 μg of respective endotoxin-free EGFR plasmid DNA (in PB513B-1 vector) using Fugene HD at 1:3 DNA:Fugene HD ratio according to manufacturer’s instructions. The cells were trypsinised 48 h later and seeded in a 10 cm^2^ dish in fresh media containing 4 μg/mL Puromycin for 7-10 days for selection. Surviving clones of cells were pooled together and checked for EGFR expression by western blotting and confocal imaging. Puromycin was routinely added at 4 μg/mL final concentration to CHO cells stably expressing EGFR during culturing.

##### Generation of Ba/F3 stable cell lines

Ba/F3 cells were cultured in phenol-red free RPMI1640 media supplemented with 10% heat-inactivated FBS, 1X Penicillin/Streptomycin and 10 ng/mL recombinant murine IL3. Puromycin was added at 2 μg/mL final concentration to Ba/F3 cells stably expressing EGFR. To establish stable cell lines, Ba/F3 cell lines were electroporated using a Neon transfection kit and MicroPorator device. On the day of electroporation, cells were washed once in PBS and resuspended in buffer R at a final concentration of 1.5 x 10^7^ cells/mL. 100 μL cells (=1.5 x 10^6^ cells) were electroporated with a mix of 0.5 μg of Super Piggybac Transposase expression vector and 5 μg of respective endotoxin-free EGFR plasmid DNA (in PB513B-1 vector) at 1600 V, three pulses, 10 ms. Electroporated cells were immediately transferred to a well of a 6-well dish containing 2 mL pre-warmed Ba/F3 cell media without antibiotic. 48 h later cells were centrifuged and resuspended in fresh media with 2 μg/mL Puromycin for selection. Cells were maintained in media containing Puromycin at a density between 0.3 x 10^6^ to 1 x 10^6^ cells/mL for 7-10 days to obtain polyclonal cell populations stably expressing EGFR.

#### Mice

In this study, young adult male (6-7 weeks old, 24.6 ± 2.1 g) NOD.Cg-Prkdc^scid^ Il2rg^tm1Wjl^/SzJ mice (NSG) were used for all animal experiments. All mice were maintained within the King’s College London Biological Services Unit under specific pathogen-free conditions in a dedicated and licensed air-conditioned animal room (at 23±2°C and 40-60% relative humidity) under light/dark cycles lasting 12 h every day. They were kept in individually ventilated standard plastic cages (501cm^2^ floor space; from Tecniplast) including environmental enrichment and bedding material in the form of sterilized wood chips, paper stripes and one cardboard roll per cage. Maximum cage occupancy was five animals, and animals were moved to fresh cages with fresh environmental enrichment and bedding material twice per week. Sterilized tap water and food were available *ad libitum*; food was PicoLab Rodent Diet 20 (LabDiet) in the form of 2.5 x 1.6 x 1.0 cm oval pellets that were supplied at the top of the cages. For imaging, animals were anesthetized using isoflurane (1.5% (v/v) in pure O _2_). After imaging, mice were either left to recover from anesthesia (by withdrawal of anesthetic) in a pre-warmed chamber or sacrificed under anesthesia by cervical dislocation. Experimental design was based on the assumption that tumor growth differences in vivo are comparable to the effects observed upon IL3 withdrawal from cells in vitro. These data were used together with an α of 0.05 and a power of ≥90% to determine minimum cohort sizes. For longitudinal experiments, cohort sizes were then oversubscribed to hedge against potential adverse effects and resultant animal sacrifice, which if premature would endanger the whole study. Consequently, cohort sizes were *N*=6. The total number of animals used was 30. No adverse events were associated with the procedures performed in this study and animals put on weight in line with strain expectations (data from Charles River UK) throughout. Sentinel animals were kept on the same IVC racks as experimental animals and confirmed to be healthy after completion of the studies.

### METHOD DETAILS

#### Site directed mutagenesis

Point mutations in plasmids expressing EGFR were introduced using Quikchange Lightning Site-directed mutagenesis kit using primer pairs listed in key resources table manufacturer’s instructions.

#### Purification of mAb-108

For purification of antibody from cell culture media, cells were resuspended at a density of 1 x 10^6^/mL in high glucose DMEM media supplemented with 2% FBS and 1 mM sodium pyruvate in a final volume of 20 mL and cultured for 5 days. Following this, cell culture supernatant was collected by centrifugation and filtered by passing through a sterile 0.2 μm PES syringe filter. Tris pH 7.5 was added at a final concentration of 100 mM, and antibody purification was performed using a mouse TCS antibody purification kit. Purified antibody was quantified using a Nanodrop and stored at 4°C.

#### Ba/F3 IL3 independent growth assay

Ba/F3 cells stably expressing EGFR were washed once in PBS and resuspended in Ba/F3 media without IL3 and puromycin. Cells were grown in the absence of IL3 for 5 days following which they were seeded at a density of 20,000 cells (in 100 μL of media) per well in a white-bottomed 96-well plate in triplicate. As a positive control for cell growth, additional cultures of each cell line were maintained in complete media (including 10 ng/mL recombinant murine IL3). These cells were seeded at a density of 20,000 cells (in 100 μL of media +IL3) per well in a white-bottomed 96-well plate in triplicate. Cell viability was measured every day for 4 days, for the positive controls, or 10 days for the cells without IL3, using CellTiter-Glo Luminescent Assay using a CLARIOstar plus microplate reader according to manufacturer’s instructions.

#### FLImP sample preparation

Only the central 4 wells of ibidi glass-bottomed 8-well slides were used. The top left of these was coated with poly-L-Lysine (PLL) and the other 3 wells were coated with 1% BSA in PBS for at least 2 h. PBS without Ca^2+^ and Mg^2+^ was used throughout.

##### Affibody labeling

CHO cells stably expressing WT-EGFR or mutant EGFR were seeded 1.8 x 10^4^ cells per well and allowed to grow for 2 days. ΔC EGFR expressing CHO cells were grown in the presence of 50 ng/ml doxycycline.

Transient transfections in CHO cells were generated for the following EGFR mutants: H566F, Lzip3S, ED/RK + L680N, ED/RK + L680N + *Lzip*3S, K946E, insNPG, G564P, G564P + ED/RK, *Lzip*3A, T766M + LZip3S and T766M + I942E. Wild type CHO cells were seeded at 1.8 x 10^4^ per well. After one day cells were transfected using Fugene HD at 1:3 DNA:Fugene HD ratio and 600 ng DNA/well according to manufacturer’s instructions. Cells were left for 24 h prior to labeling.

All samples were left in low serum medium (0.1% FBS) for 2 h before rinsing with PBS. Samples were then chilled on ice at 4°C for 10 min in PBS and labeled with 8 nM Affibody-CF640R for 1 h on ice at 4°C. Cells were rinsed with PBS and fixed with 3% paraformaldehyde plus 0.5% glutaraldehyde for 15 min on ice and 15 min at room temperature.

##### Lapatinb/Erlotinib treatment

If cells were treated with Lapatinib or Erlotinib the drug was added at a concentration of 1 µM during low-serum-starving and while labeling with Affibody.

##### EGF labeling

For EGF labeling for 2D triangles CHO cells expressing T766M EGFR were fixed, after starvation, in 3% paraformaldehyde in PBS for 15 min at room temperature and rinsed. Cells were labeled with 20 nM EGF-CF640R for 1 h at room temperature and rinsed with PBS. The samples were fixed again for 15 min at room temperature this time with 3% paraformaldehyde plus 0.5% glutaraldehyde.

##### mAb-2E9 or mAb-108 treatment

Samples that were treated with the conformational selective monoclonal antibodies mAB-2E9 or - 108 had CHO expressing WT-EGFR cells serum-starved for 2 h with 50 ng/mL doxycycline. The samples were washed with PBS and chilled on ice at 4°C for 10 min. Samples were treated with 200 nM mAb by incubating on ice at 4°C for 2 h. Cells were fixed with 3% paraformaldehyde in PBS for 8 min on ice and 8 min at room temperature. Slides were then rinsed with PBS and labeled with 10 nM EGF-CF640R for 1 h at room temperature. The cells were fixed again in 3% paraformaldehyde plus 0.5% glutaraldehyde in PBS for 15 min at room temp.

##### All samples

Samples were rinsed with PBS after fixing and stained with 1 µg/mL Hoechst in PBS at room temperature for 10 min then rinsed again with PBS. Samples were stored at 4°C and prior to imaging slides were brought up to room temperature. Poly-L-lysine was removed from the top left well and PBS from the samples wells. A 1/1000 dilution of FluoSpheres, 0.1 µm, infrared (715/755) was prepared and vortexed and then a 1/50 dilution of the 1/1000 in PBS was prepared and vortexed and 300 µl was added per well. These were used as fiducials.

Samples were loaded onto the microscope such that the central dividing cross between the central 4 wells of the plate corresponded to position 0,0. The slide was warmed to 34°C prior to sample collection and the sample was brought into initial focus by the operator. From this point, sample acquisition was managed by the acquisition system described below.

#### FLImP data acquisition

Image acquisition was achieved using an Oxford Nanoimager S (ONI Oxford, UK) single molecule imaging microscope with 1.49N oil immersion objective, operating NanoImager software (Version: 1.7.3.10248 – ef4ff2c0) set up according to manufacturer’s instructions. Single frame FLImP acquisitions were obtained using 8 mW 640 nm diode laser power for 20 ms exposure, single frame Hoechst acquisitions were obtained using 8 mW 405 nm diode laser with 20 ms exposure and single frame GFP acquisitions were obtained using 8 mW 488 nm diode laser with 20 ms exposure. Emissions were collected between 665nm and 705nm for Hoechst and FLImP imaging, and 498 nm and 551 nm for GFP respectively.

The TIRF angle was set to 54.0° for FLImP acquisition and 52.0° for Hoechst or GFP channel acquisition. In the absence of active cooling, instrument temperature was maintained at 34 °C throughout. Microscope was seated on an optical table but owing to the excellent instrument stability, the table was not floated. The PythONI (Python API from ONI), included with the NanoImager software suite was used to facilitate rapid automation of data acquisition. This process is described briefly below:

##### Fiducial imaging

The top left well of each plate (starting position: x= 3 mm, y =3 mm) contained fiducials from which PSF properties were estimated. Once active, a 3 x 3 frame region of interest was selected (0.1mm inter-frame spacing) from which single frame acquisitions were taken. For each region of interest (ROI), focal plane was first established using ONI autofocusing software, before a focal polishing step was applied as described in the following section.

##### FLImP series acquisition

Upon navigating to the appropriate sample well. A 4 mm by 4 mm region of interest was then defined for a raster tile scan. ROI was chosen based on ONI instrument reach, remaining sufficiently far from the well boundaries as to avoid artifacts. On moving to each ROI, the following procedure was followed:

##### Focusing

While the inbuilt ONI autofocusing system performed reasonably at maintaining focal plane throughout acquisition, an additional focal polishing step was required for prolonged data acquisitions and was run at ∼20 min intervals. This algorithm has also since been used to find optimal focusing position from microscope “cold start”. Briefly, from a starting position approximating the focal plane a z-stack of images was recorded over +/-2μm with 200 nm step. A 3×3 pixel top-hat filter was employed to extract diffraction limited objects from the images. Each frame was then cross-correlated with a 2D Gaussian kernel approximating the experimentally determined PSF size. Next, by fitting and then finding the minimum of a second order polynomial an estimate of the focal plane could be extracted. Successive applications of this algorithm permitted discovery of the focal-plane and typically took no more than ten seconds per ROI.

##### ROI detection

For imaging of fiducials a collection of ∼600 FLImP series was acquired from each region of interest at 0.1mm intervals in x and y. This interval was sufficient to ensure results were not affected by bleaching from neighboring acquisitions and no further evaluation of labeling extent was required. For experiments containing cells, as fewer fields of view (FOV) were likely to contain FLImP suitable series, a larger search area was required.

The FOV of the ONI NanoImager is 0.05 mm by 0.08 mm, yielding a potential of ∼2280 ROIs per 3 x 3mm search area. Assuming average acquisition time of 1 minute per ROI and 800 MB, it would take over 100 hours to record FLImP series of the entire slide of which the majority would be of little biological relevance. To overcome this limitation, a method to rapidly evaluate the likely utility of each FOV was required.

A FOV likely to yield a good FLImP series comprises of a region that contains one or more cells that are sufficiently labeled to be likely to yield FLImP suitable track time series. A sufficient proportion of the ROI must be cell-free in order that sufficient fiducials are present to enable adequate drift correction. Here, cells are required to be nucleated or have an intact cytosol to ensure that cells have not become detached from the side leaving behind membrane patches or “ghosts”, which may have altered physiological status despite exhibiting some FLImP labeling. To achieve this objective, samples were counter-stained with the nuclear stain Hoechst, or cytoplasmically expressed GFP, depending on cell sample to be investigated.

While an image segmentation problem such as this may appear to lend itself to deep learning, and such techniques have recently been successfully employed for similar image segmentation problems,^73^ such approaches do have limitations, particularly the ongoing difficulty in clinical translation of technologies using deep learning owing primarily to the black-box nature of decision­making processes □. As such, a much more simple and tractable classical image segmentation approach was used to evaluate the suitability of each FOV yielding two tunable parameters: fraction of cell labeling (Hoechst or GFP) and fraction of FLImP labeling.

Briefly, the extent of Hoechst or GFP labeling was determined in each FOV to establish the presence of intact cells and/or cells that express the desired phenotype by removing background using Otsu thresholding^74^ followed by removal of small objects (fiducials etc.) smaller than 500 pixels (∼2.5 μm^2^). By averaging the resulting binary mask, the fraction of FOV that was Hoechst or GFP positive is returned. If this fraction cell labeling parameter is above sample specific critical threshold (i.e., > 0.1), then the second step of evaluation commences.

Here, the extent of FLImP labeling within the FOV is evaluated by first using Triangle method of thresholding^75^ to segment the extent of FLImP labeling from background. When used in conjunction with the binary mask for cell labeling area obtained above, the fraction of FLImP labeling within cells is returned. Finally, if the fraction of FLImP labelling is above a sample specific threshold (i.e., > 0.05), a FLImP series is recorded for the current FOV.

Failure to pass either of these checks results in the ONI moving to the next FOV. Single images of FOVs are recorded in each instance to enable post-acquisition user validation. The process of labeling extent evaluation typically took no more than 2 seconds per FOV. Finally, once an FOV was determined to be suitable a FLImP series acquisition was made comprising 1200 frames with 20 ms exposure time and 20 mW 640 nm laser power.

#### Western blot

CHO cells were seeded at a density of 2 x 10^5^ cells per well in a 6-well plate and transfected the following day using Fugene HD at 1:3 DNA:Fugene HD ratio according to manufacturer’s instructions. 24 h post-transfection, the cells were serum starved in media containing 0.1% FBS for 2 h at 37°C.

Cells were placed on ice, washed once in cold PBS and lysed directly in 6-well dishes in cell lysis buffer [50 mM Tris/HCl (pH 7.4), 1 mM EDTA, 1 mM EGTA, 50 mM sodium fluoride, 5 mM sodium pyrophosphate, 10 mM sodium β-glycerol 1-phosphate, 1 mM dithiothreitol, 1 mM sodium orthovanadate (New England Biolabs Inc), 0.27 M sucrose, 1% (v/v) Triton X-100, 1x Protease inhibitor]. Cell extracts were clarified by centrifugation, and the protein concentration was determined using Bradford assay (ThermoFisher Scientific). 10-20 µg of total cell lysate was resolved on an 8% Bolt Bris-Tris gel (ThermoFisher Scientific), and proteins were transferred to PVDF membrane using Mini Trans-Blot Cell system (Biorad). The PVDF membrane was blocked in 5% non-fat milk/TBS+0.1% Tween (TBST) for 1 h at room temperature and incubated overnight in 1:1000 dilution of primary antibody [EGFR D38B1, pEGFR Tyr1173 53A5, pEGFR Y1068 D7A5, EGFR polyclonal, pEGFR Y-12] or Actin-HRP 13E5 1:5000 dilution in 5% BSA/TBST at 4°C. After extensive washing in TBST, the membranes were incubated with 1:5000 dilution of the appropriate secondary antibody in 5% non-fat milk/TBST. Subsequently, membranes were incubated with Immobilon ECL Ultra Western HRP substrate solution and imaged on a Biorad ChemiDoc MP System imager. The images were quantified using Biorad Image Lab software where required.

#### Single particle tracking

Cells were seeded at a density of 1 x 10^5^ cells/dish on 1% BSA-coated 35 mm no. 1.5 (high tolerance) glass-bottomed dishes in 2 mL of media. At 18 h post seeding, transient transfections were performed using Fugene HD as described above, and cells were grown for further 24 h. Prior to imaging, cells were starved for 2 h at 37°C in 0.1% serum. Treatments with Erlotinib and Lapatinib were performed as described above. Cells were then rinsed twice with 0.1% serum pre-heated at 37°C and were labeled with a 1:1 mixture of 8 nM Alexa 488 Affibody/CF640R-Affibody for 7 min at 37°C. Cells were rinsed twice with low serum medium pre-heated at 37°C and promptly imaged. Single-molecule images were acquired using an Axiovert 200M microscope with a TIRF illuminator (Zeiss, UK), with a ×100 oil-immersion objective (α-Plan-Fluar, NA =1.45; Zeiss, UK) and an EMCCD (iXon X3; Andor, UK). The microscope is also equipped with a wrap-around incubator (Pecon XL S1). The 488 and 642 nm lines of a LightHub laser combiner (Omicron Laserage GmbH) were used to illuminate the sample and an Optosplit Image Splitter (Cairn Research) was used to separate the image into its spectral components as described previously^76^. The field of view of each channel for single-molecule imaging was 80 × 30 µm. Typically, for each condition, at least 30 field of views comprising one or more cells were acquired from a total of at least 3 independent biological replicates.

#### Confocal imaging

For all confocal experiments, cells were seeded at a density of 0.6^5^ cells/dish on 8-well Ibidi glass-bottomed multiwell slides, as above, and cultured for 48 h at 37°C. Prior to labelling, cells were starved 2 h at 37°C with serum-free media + 0.1% FBS, then rinsed twice in ice-cold 1x PBS and cooled down for 10 min on ice.

For anti-EGFR Affibody and EGF competition binding experiments, cells were then pre-treated with either 200 nM, 400 nM or 600 nM of ice-cold Anti-EGFR Affibody-CF640R in PBS or with mock treatment (just PBS) for 1 h at 4°C, rinsed 2x with ice-cold PBS and fixed with 3% PFA in PBS for 30 min at 4°C. After fixation, cells were rinsed again 3 timeswith room temperature 1x PBS to eliminate paraformaldehyde residues and labelled with 400 nM EGF-Alexa488 in PBS for 1h at RT, then rinsed twice with RT PBS, fixed with 3% PFA + 0.5% GA for 15 min at RT and rinsed 3 times with PBS.

For the mAb-2E9 binding experiments, after starvation cells were then pre-treated with 200 nM of mAb 2E9-AF488 in PBS or with mock treatment (just PBS) for 2 h at 4°C, rinsed 2x with ice-cold PBS and fixed with 3% PFA in PBS for 15 min at 4°C. For the EGF binding post-fixation test, cells were labelled with 200 nM EGF CF640R for 2 h at 4°C prior to fixation.

After fixation, cells were rinsed again with room temperature 1x PBS to eliminate paraformaldehyde residues and labelled with 200 nM EGF CF640R in PBS for 1 h at room temperature, then rinsed twice with PBS, fixed with 3% paraformaldehyde + 0.5% glutaraldehyde for 15 min at room temperature and rinsed with PBS.

All samples were stored in PBS at 4°C until the time of acquisition, and allowed to pre-warm at room temperature, wrapped in foil, before loading on the microscope.

In all cases, image acquisition was performed on an Elyra PS1, using Zen Black v2.3 SP1.

For the Anti-EGFR affibody and EGF competition binding experiment, the signal from Anti-EGFR affibody-CF640R was collected with 633 nm laser excitation (power = 2%) through a PMT detector in the 638-755 range, with gain = 550 and offset = 0, while the EGF-AF488 signal was collected with 488 nm laser (power 2%), through a PMT detector in the range 493-630 with gain = 700 and offset =0. Wavelengths were split using a 488/561/633 MBS filter and the pinhole size was set to 90μm. 1584 x 1584 pixel images were acquired at 1.32 μs pixel dwell time, switching laser line and detection between frames.

For the mAb-2E9 experiments in the confocal microscope, the signal from EGF-CF640R was collected with 633 nm laser excitation (power = 0.5%) through a GaASP detector in the 566-685 range, with gain = 700 and offset = 0, while the mAb 2E9-AF488 signal was collected with 488 nm laser (power 5%), through a PMT detector in the range 493-556 with gain = 700 and offset =0. Wavelengths were split using a 488/561/633 MBS filter and the pinhole size was set to maximum. 1024 x 1024 pixel images were acquired at 16.38 μs pixel dwell time, switching laser line and detection between frames.

##### Mice tumor models

Male NSG mice were used to establish subcutaneous tumor models (in right flanks) with indicated stable Ba/F3 cell lines. After acclimatization, mice were randomly allocated into four cohorts with six individuals each, shaved on their flanks, and then subcutaneously received each 2 x 10^6^ tumor cells suspended in 100 μL phosphate buffered saline (PBS). Tumor growth was followed by calipers and tumor volumes calculated using the formula: 0.5 × L × W^2^, wherein L represents tumor length and W its width. Tumor models were grown to compare tumor growth between cohorts. The experimental endpoint was defined by the time the human endpoint was reached for the cohort with the largest tumor growth, and then all animals were sacrificed.

##### In vivo imaging of tumor models

*In vivo* GFP fluorescence imaging of superficial tumor models was performed to visualize tumor growth in some animals per group over time and to quantify tumor growth differences in all animals at the experimental endpoint. The imaging device was an IVIS Spectrum *in vivo* imaging system (PerkinElmer, USA) equipped with excitation and emission wavelength bandpass filters of 465 ± 15 nm and 520 ± 10 nm, respectively. Regions of interest (ROI) were manually drawn including the whole tumor (or the injection sites where no tumors were visible) and used to calculate the radiant efficiency.

##### Tissue staining and histologic analysis

Formaldehyde-fixed paraffin-embedded (FFPE) tissues were prepared using standard methods. 3 µm tissue sections were cut using a microtome and adhered to poly- *L*-lysine slides, dried overnight at 37°C, de-waxed and subjected to antigen retrieval in a pressure cooker at pH 6.0. Morphologic analysis of tumor tissues was performed on hematoxylin- and eosin-stained sections. For antibody staining, sections were blocked (Dual Endogenous Enzyme Blocking Reagent) in 1% (w/v) BSA for 60 min at room temperature, incubated with indicated primary antibodies at 4°C overnight before being stained with a horse radish peroxidase-conjugated secondary antibody (2 µg/mL in TBS) for 60 min at room temperature. Samples were developed using the Liquid DAB+ Substrate Chromogen System and counterstained with hematoxylin before mounting. Slides were scanned using a Nanozoomer (Hamamatsu, Japan) with images being analyzed and processed by ImageJ v1.54.

#### MD simulations and structural modelling

##### Ectodomain mutations – H2H^ect^ simulations

To simulate the behavior of the H2H^ect^ dimer, we used the model of the asymmetric dimer seen in the crystal packing of the PDB entry 4KRP^77^ after removing 9G8-NB and adding the TM helix that we used in our previous work.^43^ The G564P and H566F mutations in each monomer were introduced to the WT structure using MODELLER.^78^ The models of the WT and each mutant were N-glycosylated at N151, N172, N328, N337, N389, N420, N504, N544, N579 with the core glycan (Man3GlcNAc2-Asn) and one site (N32) with the “Pauci-mannose” glycan type (Man3GlcNAc2(Fuc)-Asn) as this site has been reported to be fucosylated.^79^ The *in-silico* glycosylation was carried out using the CHARMM-GUI Glycan Modeler tool and the CHARMM carbohydrate force field was used in the performed simulations.^80, 81^ The glycosylated ECD was embedded in a membrane whose lipid composition was chosen such that it mimics the lipid composition seen in the membrane of CHO cells, i.e. with a ratio of 1-palmitoyl-2-oleoyl-sn-glycero-3-phosphocholine (POPC), cholesterol (CHOL), and n-palmitoyl-sphingomyelin (PSM) of POPC : CHOL : PSM = 30 : 11 : 2.^82^ A symmetric 250 Å x 250 Å lipid bilayer patch was generated using CHARMM-GUI’s input generator.^83, 84^ The generated model was parameterized using the CHARMM36m force field^85^ at pH 7.4 and CHARMM36 parameters were used for the lipids and glycans as provided by CHARMM-GUI. We made sure to maintain the same protonation state of the protonated residues across all systems. The systems were solvated with modified TIP3P-CHARMM water molecules.^86^ The generated model was parameterized using the CHARMM36m force field^85^ atpH7.4andCHARMM36parameterswereusedforthelipidsand glycans as provided by CHARMM-GUI. We made sure to maintain the same protonation state of the protonated residues across all systems. The systems were solvated with modified TIP3P-CHARMM water molecules,^86^ and Na^+^ and Cl^-^ ions were added to reach neutrality and the final ion concentration of 0.15 M.

All simulations were run using the GROMACS v.2020.3 package.^87^ The potential energy of each system was first minimized using the steepest-descent algorithm. Each of the minimized systems underwent a six-step equilibration protocol in the NVT and NPT ensembles, as provided by the CHARMM-GUI, where the restraints on the lipid, glycan and protein atoms were gradually relaxed. During each step of the equilibration, the temperature and pressure were kept constant at 310 K and 1 bar, respectively, using the velocity-rescale thermostat^88^ and the Parrinello-Rahman barostat^89^. All bond lengths to hydrogen atoms were constrained using the LINCS algorithm,^90^ while van der Waals interactions were treated with a cut-off of 12 Å. Electrostatic interactions were computed using the particle mesh Ewald method^91^ with the direct sum cut-off of 12 Å and the Fourier spacing of 1.6 Å. Post equilibration, the production run for each of the WT and mutants was 2.5 μs long, where no restraints were applied.

##### Kinase mutations – Bb2Bb^kin^_dimer_ simulations

The simulations of EGFR kinase starting from the Bb2Bb^kin^_dimer_ were based on the dimers seen in the crystal packing of the kinase domain of the PDB entry 3VJO.^52^ The co-crystalized ligand was removed from both monomers, and the missing atoms were built with MODELLER^78^. Each dimer consisted of the sequence G696-A1022 (in the numbering with the 24-aa tag). The two non-naturally occurring R938E and K946E mutations, as well as the NSCLC D770-N771insNPG (insNPG) mutation on the αC/β4 loop, were introduced in each monomer of the WT dimer structure with MODELLER^78^. In the case of the insNPG, the structure of each monomer of the resulting modelled dimer was then compared to the crystal structure of insNPG in complex with a covalent inhibitor (PDB ID 4LRM) to ensure that the overall conformation of each monomer and the environment around the point of the mutation are in accordance with the experimentally derived structure. For the unbiased MD simulations, each simulated system was parameterized using the CHARMM36m force field^85^ at pH 7.4. The protonation states of the residues were determined by PlayMolecule^92^, which left all the residues in their usual charge states. The systems were solvated with modified TIP3P-CHARMM water molecules^85^ in a dodecahedral box with periodic boundary conditions, while Na^+^ and Cl^-^ ions were added to reach neutrality and the final concentration of 0.15 M.

All simulations were run using the GROMACS v.2020.3 package^87^. Prior to the production simulations, the energy of each system was minimized through steepest-descent energy minimization. Each system was equilibrated using the following protocol: the initial velocities for the atoms were taken from Maxwell distribution at 300 K, and the system was simulated for 5 ns at the NVT ensemble using a velocity rescaling thermostat^88^ and position restraints on heavy atoms (1000 kJ mol^-1^ nm^-2^), followed by 10 ns in the NPT ensemble under constant pressure (1 bar) using the Berendsen barostat^93^, followed by 5 ns using the Parrinello-Rahman barostat.^89^ All bond lengths to hydrogen atoms were constrained using the LINCS algorithm,^90^ while van der Waals interactions were treated with a cut-off of 12 Å. Electrostatic interactions were computed using the particle mesh Ewald method^91^ with the direct sum cut-off of 12 Å and the Fourier spacing of 1.6 Å. Each production run was 4 μs long.

### QUANTIFICATION AND STATISTICAL ANALYSIS

#### Single molecule tracking

##### Feature detection and tracking

All single-molecule time series data (for FLImP and single molecule tracking) were initially analyzed using the multidimensional analysis software described previously^94^. Briefly, this software performs frame-by-frame Bayesian segmentation to detect features, and then performs a least-squares Gaussian profile fit to locate detected features to sub-pixel precision, then links these features through time to create tracks using a simple proximity-based algorithm. For multichannel data (for colocalization analysis) the software determines cubic polynomial registration transformations from images of fluorescent beads and performs feature detection and tracking independently in each channel before applying the transformations to transform all tracked positions to a common channel.

##### Track colocalization analysis

The colocalization event duration analysis was performed in the same way as in.^43^ Briefly, single particle tracks were extracted from the two imaging channels and registered as described above. The duration (T_ON_) of individual events in which a track in one channel moves within a pixel (160 nm) of a track in the other channel and then they move apart again was calculated. To reduce the impact of localization error on these results a temporal Gaussian smoothing filter of FWHM 4 frames (200 ms) was applied to the position traces before the colocalization analyses.

#### FLImP data analysis

##### Drift determination

An automatic algorithm for the identification of fiducial markers and exclusion of outliers was used to measure drift in FLImP videos so it could be accounted for in later analysis. Briefly, a population of 200 fiducial markers was first identified by selecting the brightest objects from the video that persisted for at least 80% of total runtime. Then, a modified cross-validation based approach was employed where the population of 200 selected fiducials were randomly divided into three bins with a 20:40:40 split. Here, the first bin represents a discard bin, excluded from further analysis that attempts to capture outlier measurements. Drift was then calculated independently for tracks split between the remaining two bins before scoring the drift correction quality by recording the residual sum of squares (RMS) between the independent partitions. Here, tracks were subdivided at positional change points. After 200 iterations of this cross-validation scheme, the partition with the smallest RMS (bins2 and 3) were pooled for determination of a final FLImP video drift measure. This process is outlined in more detail in the **Supplemental file.**

##### Track selection

Each FLImP video typically returned between 1,000 and 10,000 track objects of which only a small subset was suitable for FLImP analysis. Previously, identification of FLImP suitable tracks was a laborious process, requiring trained operators to manually trawl track lists from each FLImP series in order to identify tracks that may be suitable for downstream FLImP fitting processes. In the present study a sequential filtering approach to track selection was employed, which involves passing a population of FLImP tracks through a series of pre-determined filters to rapidly and automatically identify tracks suitable for further FLImP analysis. As tracks with increasing numbers of fluorophores represent nested sets, temporal subset tracks may be suitable for further FLImP analysis, even if the entire track is not. Filters were organized so that the most computationally resource intensive filters were located towards the end of the pipeline, to minimize resource requirement and accelerate this aspect of the process. A detailed description of each filter is provided in the Supplemental file. Filter parameters were developed heuristically on training datasets independent of those presented in the results section of this manuscript and were then held constant throughout. The result is a list of tracks suitable for FLImP analysis, with temporal ranges corresponding to 1 and 2 (and for 2D FLImP 3) fluorophores simultaneously fluorescence identified. This process is outlined in more detail in the **Supplemental file**.

##### FLImP localization fit

As described in^95^ the FLImP fitting process determines the locations of the two molecules in each track and their uncertainties by fitting a model of two overlapping, photobleaching Gaussian fluorophore PSF profiles to the ROI of the spot during the frames identified by the track selection procedure (described above). Briefly, a 7-parameter least square fit is performed, where the parameters are the intensity and *x* and *y* location of the two fluorophores and their common PSF size. Which fluorophores to include in the model at each time point is identified as part of the track selection procedure. The drift of the sample is accounted for in the fitting process, by assuming the fluorophores move in time following the drift while remaining fixed with respect to one another. The fitted profile of other nearby features in the ROI detected by the feature detection is remove from the ROI images before this FLImP fit is performed to remove their contribution. By repeating the fit 2400 times, each time resampling with replacement from the *n_x_ ×n_y_ ×n _t_* intensity values (the unravelled ROI-time image volume), we obtain an empirical estimate of the uncertainties in the parameters. From this we can obtain an empirical probability distribution, or posterior given the data, for the fitted parameters and anything which can be calculated from them, such as the fluorophore separation. We showed in^95^ how this approach provides robust estimates including of the uncertainties without need for an explicit model for image noise.

##### 1D decomposition

With the assumption that there is a fixed set of molecular structures in our sample each FLImP measurement should correspond to one of the fixed set of discrete pair-wise separations between the fluorescent labels in the samples. We perform a Bayesian parameter estimation using the Metropolis Hastings Markov Chain Monte Carlo sampling algorithm and average Bayesian Information Criterion (*AVBIC*) to optimally and objectively determine the number of discrete separations and their values required to explain the population of FLImP measurements for each condition. It accounts for ambiguity in correspondence between each measurement and proposed separation components (*assignment sampling*) and allows for the presence of a population of spurious measurements not sampled from the structure (*clutter*). We devise a new measure of the separation between molecules, *r_Δx_*, which does not suffer from the problems of the typically used L2 norm between uncertain localizations at short separations (**Supplemental File).** This new measure allows unbiased estimation of separations down to zero separation. Localization errors can make posteriors with negative *r_Δx_* for short separation measurements. We therefore use its absolute value, |*r_Δx_*|, in some places as appropriate.

In the decomposition. plots, e.g., Figures 1D, 1E:

- The grey background curve is the sum of the individual FLImP measurement posteriors of. |*r_Δx_*|
- The coloured peaks are the posteriors of each individual separation component from our MCMC sampling, with the area under each weighted by the number of measurements assigned to it (its abundance).

We define also the abundance-weighted separation posterior, P_w_(|*r_Δx_* ||*D*), or marginalized, weighted probability distribution, the sum of the coloured peaks. This ignores the assignment of each component and just asks how much evidence the decomposition gives for the presence of each separation in the sample.

For more detail see the 1D FLImP decomposition section in the **Supplemental File.**

##### Posterior comparisons and bootstrap resampling

To compare and contrast the decomposition of different FLImP datasets, we compare the abundance-weighted posteriors, P_w_(|*r_Δx_* ||*D*), between samples. To take some account of the finite number of FLImP measurements in each dataset, which is typically *n* = 100 measurements after the measurement quality filter has been applied, we use bootstrap resampling with replacement. For each dataset we created 20 datasets each also of size *n* by bootstrap resampling with replacement from the *n* measurements in the true dataset. We repeat the Metropolis-Hastings decomposition process separately on each of these, for the number of components, *N*, determined from the *AV BIC* test on the true (unresampled dataset). For each of these we also calculate P_w_(|*r_Δx_* ||*D*). The variation between these curves at each *x* gives a measure of the uncertainty due to the finite number of measurements comprising the distribution. In (add type of plot) plots we show P_w_(|*r_Δx_* ||*D*) for the true data as a solid line and an error band showing the 20%–80% percentiles of the bootstrap sample distribution of P_w_(|*r_Δx_* ||*D*) as a function of |*r_Δx_*|. This enables comparison between conditions to consider the significance of difference between distributions.

##### Multidimensional scaling analysis (MDS)

Distances between each pair of abundance-weighted separation posteriors were measured using the Wasserstein metric,^55^ whereby identical distributions would have a Wasserstein metric of zero and more different distributions would exhibit larger positive values. As the Wasserstein metric satisfies the triangle inequality,^96^ these distances were used to construct a distance matrix between all pairs of distributions before the Multidimensional scaling (MDS)^97^ dimensionality reduction technique was applied to this distance matrix to produce a simplified visualisation of the similarity of the conditions investigated. SCREE plot evaluation via heuristic breakpoint detection suggests most of the variability in the dataset could be described by two principal components and conditions that produce more similar distributions of separations would appear more closely together in the plot. As the Wasserstein metric does not provide information regarding the direction of the change (whether oligomers are getting larger or smaller between conditions), plots were overlay with a vector denoting the directionality of increasing oligomer size, defined as an increasing fraction of the FLImP distribution that comprised separations greater than 20 nm). An estimate of the uncertainty in the position of each condition within the MDS plot was obtained by including 1D-MCMC FLImP posteriors derived from 20 bootstrap resamplings of the 1D-FLImP datasets for each condition in the calculation of distance matrix and subsequent MDS analysis, fixing the number of components in each case ***a priori*** to that obtained when fitting the entire dataset. This analysis permitted the addition of confidence ellipses to the MDS plots that represent the 95% Confidence interval for each condition. Here, the highlighted centre of each ellipse is taken as the 1D-MCMC FLImP posteriors derived from fitting the entire (non-bootstrapped) dataset.

##### 2D FLImP triangle pooling

2D FLImP is an extension of 1D FLImP whereby FLImP measurements are extended to include three fluorophores per measurement object and the FLImP localisation fit extended to fit a sum of 1, 2 or 3 overlapping PSFs at intervals identified by the track selection procedure. As the number of fluorophores present within FLImP track objects represent a nested set, meaning the same track selection criteria as is used for 1D FLImP can be trivially extended to the 2D case containing three or more fluorophore objects and 2D FLImP tracks are presently rarer than 1D FLImP measurements. 2D FLImP imaging returns a population of triangles, providing information about the separation of a group (three or more) fluorescently labelled locations in the sample of interest. Our interpretation assumes that multiple FLImP measurements can be treated as a representative sample from all the possible separation combinations between labelled locations in the experimental sample. Distances between each pair of 2D FLImP abundance-weighted separation posteriors were measured using the Wasserstein metric,^55^ whereby identical distributions would have a Wasserstein metric of zero and more different distributions would exhibit larger values. As the Wasserstein metric satisfies the triangle inequality.^96^ Assuming there is a finite, discrete and precise set of separations between labelled locations in our structure(s), we wish to inter from our 2D FLImP measurements the number of these separations, their value and uncertainties and relative position in 2D space. Optimally exploiting the many highly quantitative 2DFLImP measurements and using the revised separation measure, described elsewhere, should give more robust, precise, and unbiased structural information down to zero separation. [D] The Distance matrix of triangle-relatedness was used to construct a dendrogram with optimal cuts determined using Bayesian Hierarchical clustering approach as described In,^98^ The resulting grouped triangles were optimally aligned before pooling as illustrated in [E].

##### Measurement quality filtering

Datasets of FLImP measurements for a particular condition typically contain many hundreds or even thousands of individual single molecule track measurements with a range of localization error and other qualities. Beyond the filtering for suitable tracks for FLImP analysis described above in the Track Selection section, after the FLImP fit to each track selection has been performed we can further narrow down the dataset to good quality results by applying a set of filters based on properties of the fitted measurements. By removing low resolution, poor quality results we speed up decomposition and other analysis of the FLImP measurements to increase the resolution for the decomposition without compromising it. We filter them as follows:

- Only keep measurements whose separation between determined fluorophore locations is *r* < 80nm. We are not interested in longer separations.
- Only keep measurements for which the 69% confidence interval (CI) in *r_Δx_ <* 8nm. This means only high localization precision measurements are used. Note this filter uses the *r_Δx_* CI which is not correlated with separation, rather than the |*r_Δx_*| separation which is. This ensures this filter does not bias the separation distribution which results.
- Measurements where the localization posterior of either fluorophore has too high an asymmetry are rejected as this may be a sign of a poor measurement. The ratio of the minor axis to the major axis of the (*x,y*) location posterior covariance ellipse must be > 0.5 for both spots in the measurement.

- Finally, of the measurements which pass the above filters we keep the 100 highest resolution measurement, with resolution defined as the 69% *r_Δx_* CI.

#### Analysis of confocal images

Colocalization analyses were carried out in images of 600 μm optical slices and performed using Huygens software (Scientific Volume Imaging)

For the mAb 2E9 binding experiments, pixel-wise intensity or intensity ratio distributions were extracted from the data using Huygens (SVI). Data were loaded as pandas-data frames using Pandas 1.3.5 and analyzed using Pandas, Scipy 1.7.3 and Sckit Posthocs 0.6.7 through Jupyter notebooks 6.4.1 in Python 3.7.13.

The non-parametric Kruskal-Wallis statistical test was performed and T-test post-hoc analysis with Bonferroni multiple comparison correction was applied to calculate P Values.

Python packages Matplotlib 3.5.1 and Seaborn 0.11.2 were used to generate the plots.

#### Statistical analysis for mice studies

Prism software version 9 (GraphPad, La Jolla, USA) was used to calculate all statistical parameters as indicated. Generally, p-values were calculated using significance levels of α = 0.05. In-text numbers indicate means of pooled data ± standard deviation (SD) unless otherwise stated.

#### Key resources table

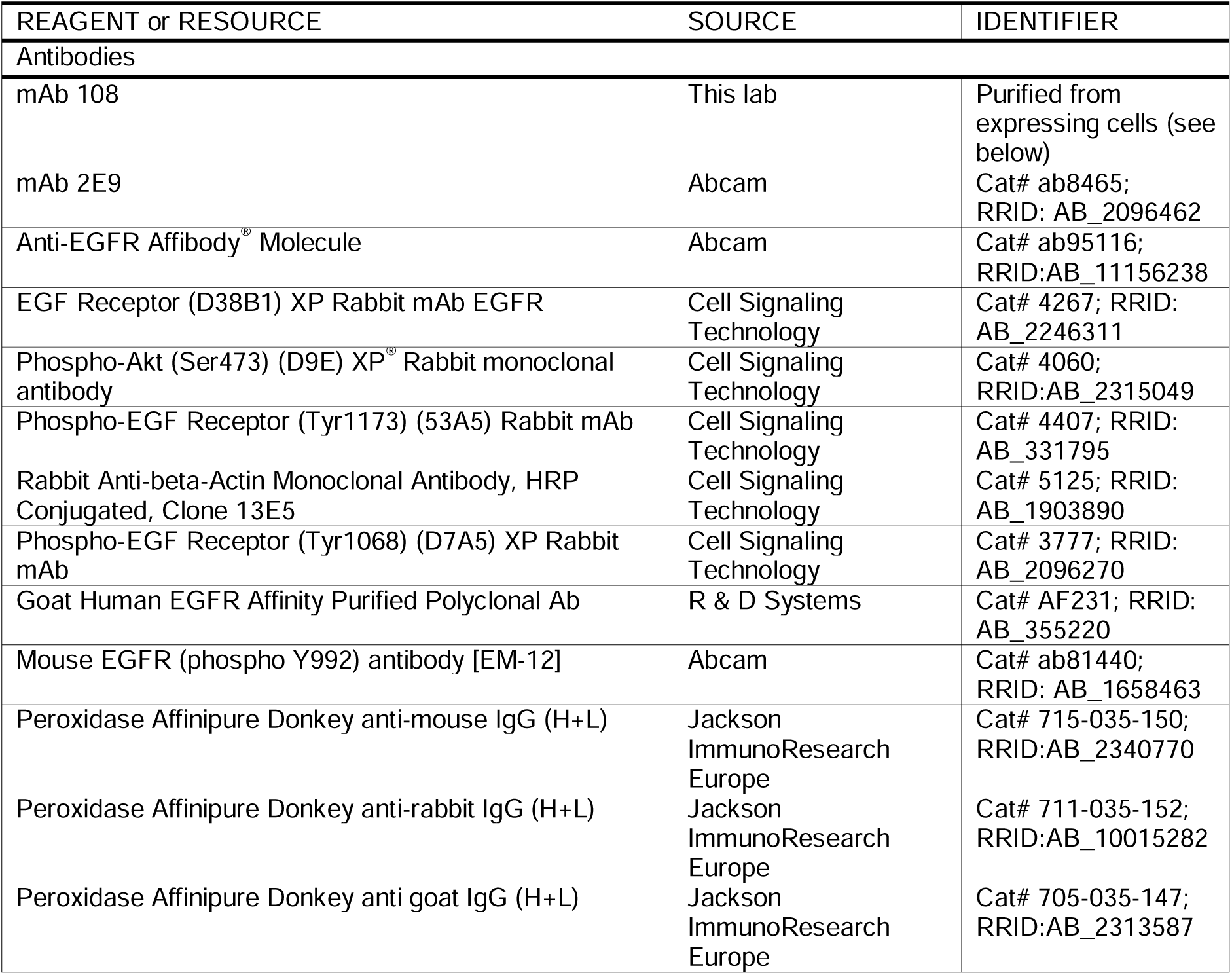
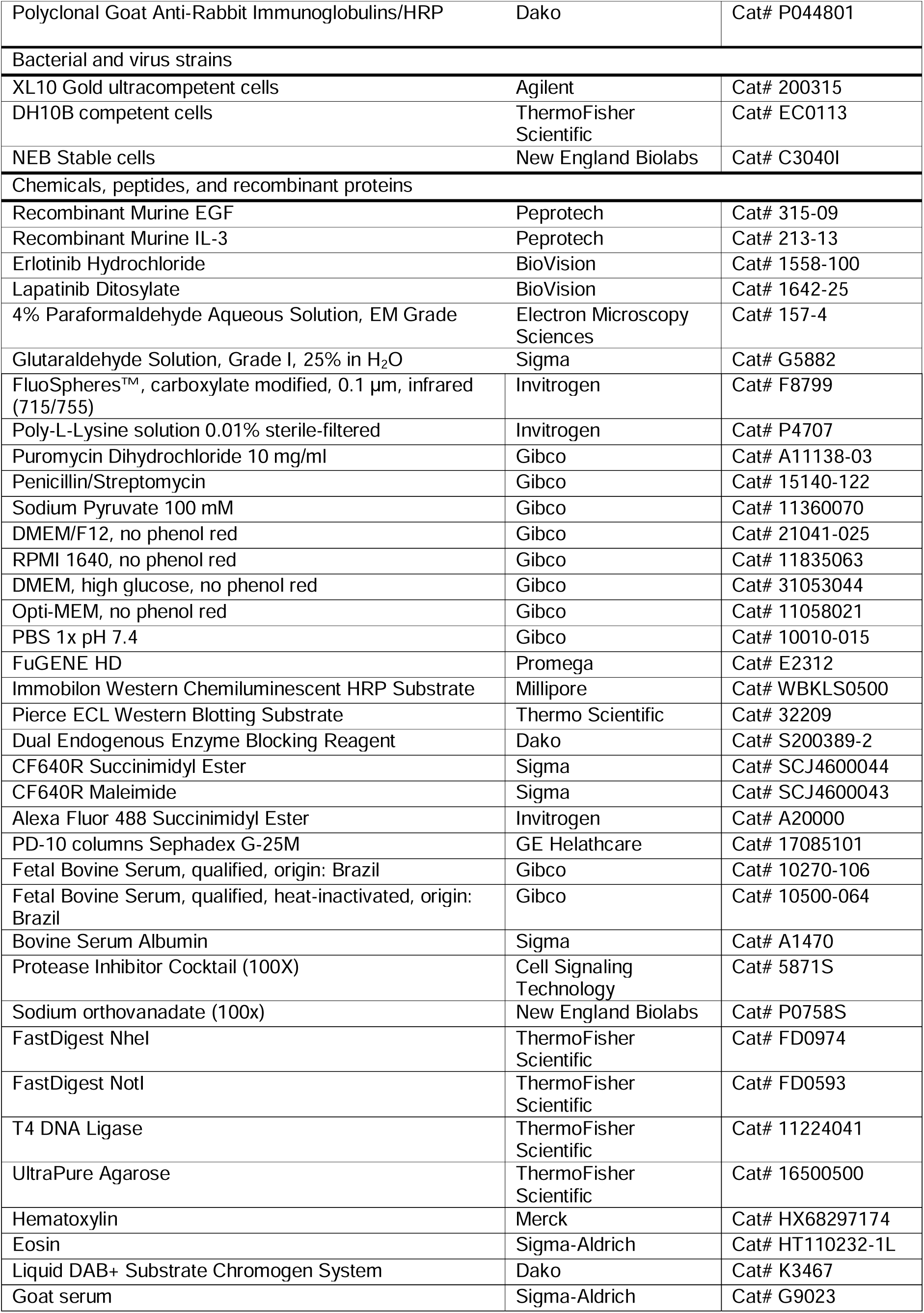
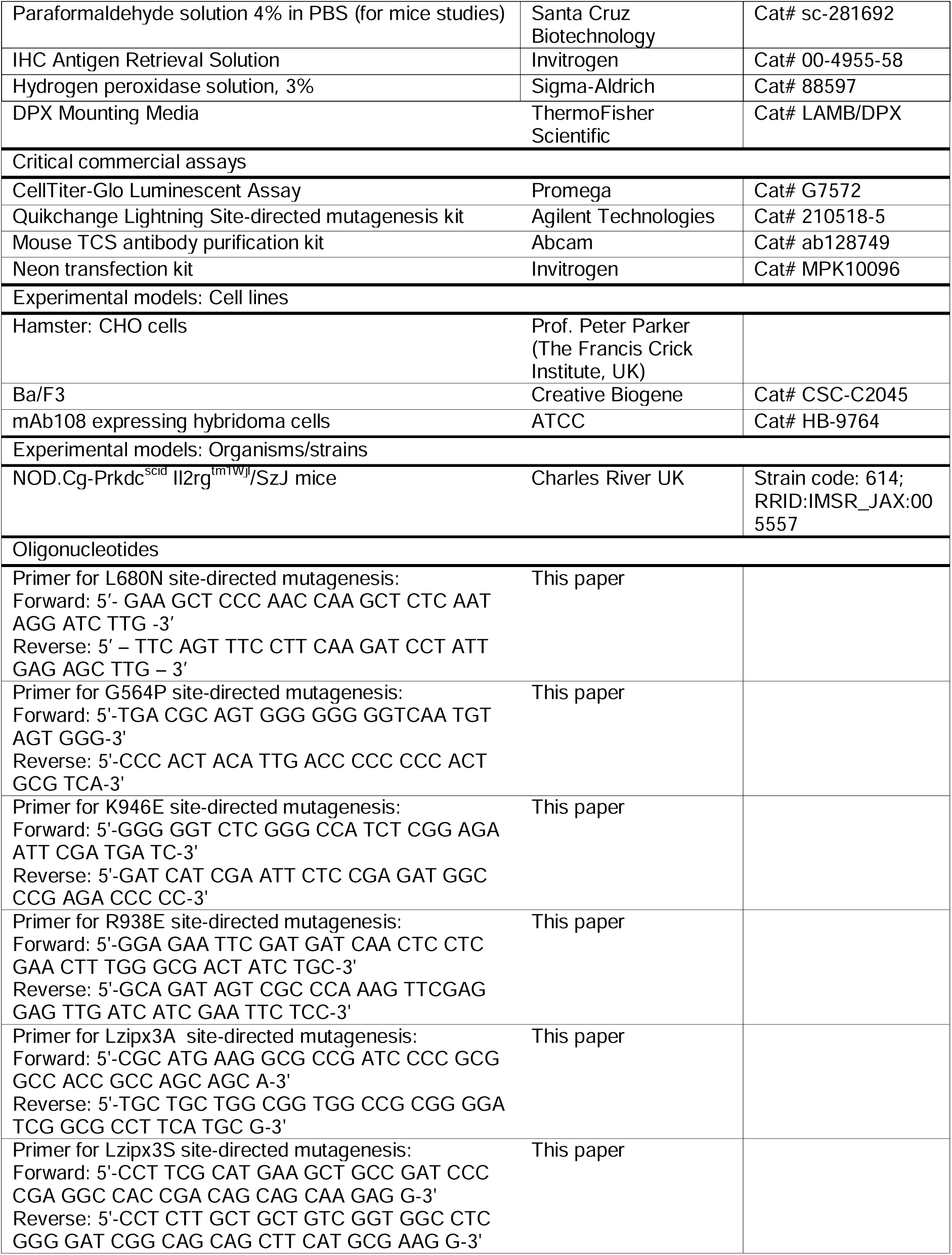
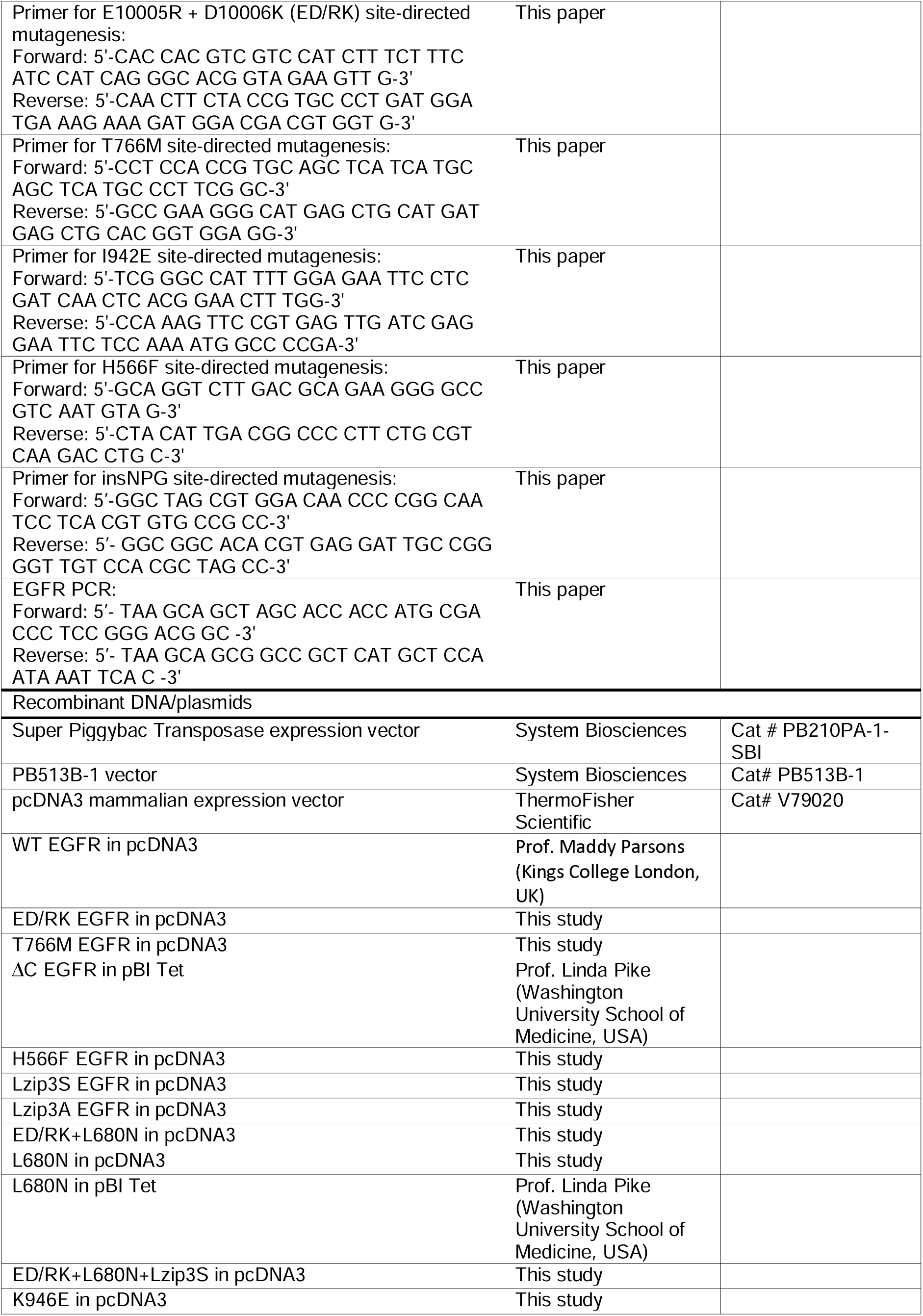
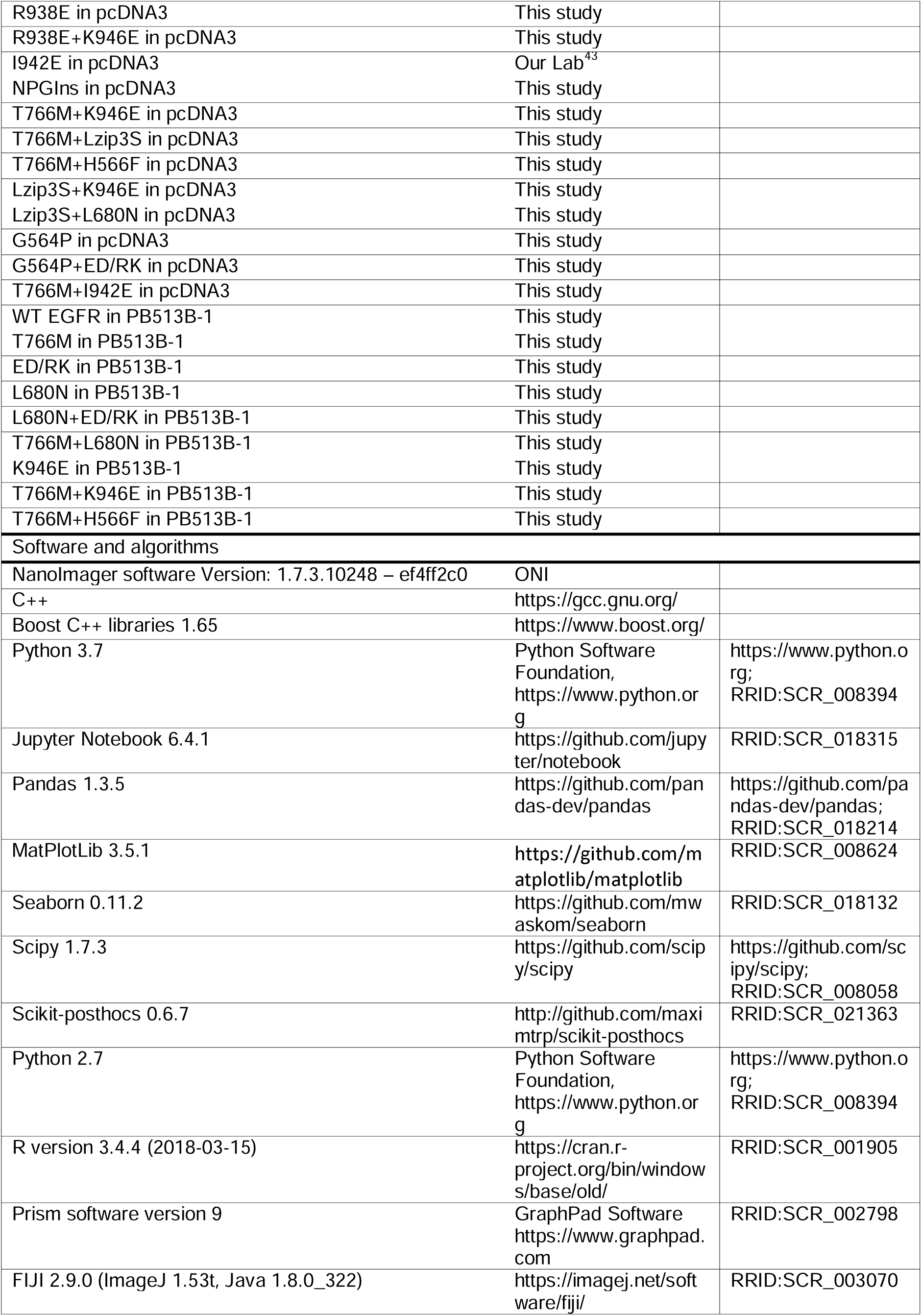
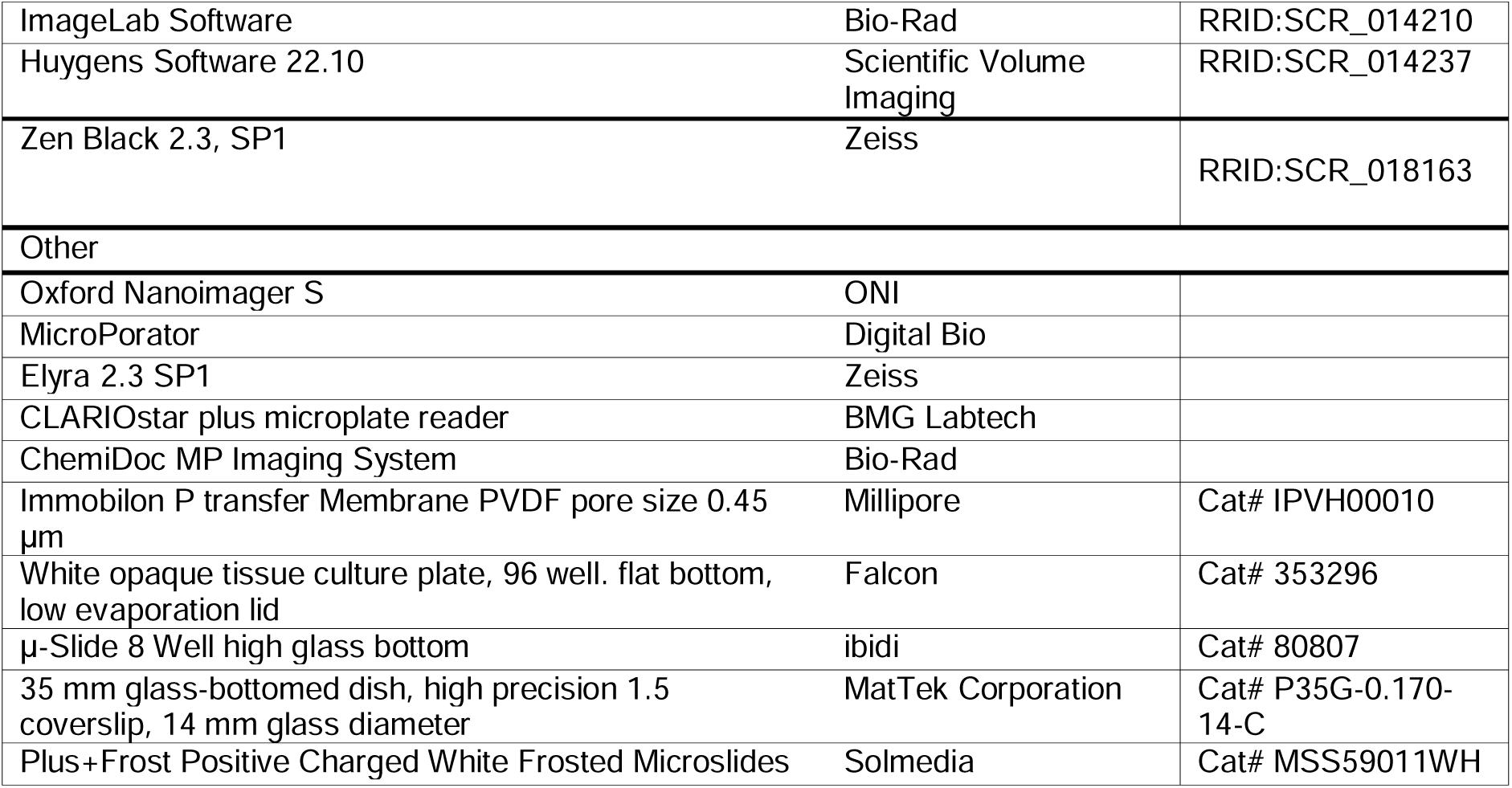

