## Supplemental information for "The T766M-EGFR lung cancer mutation promotes tumor growth by exploiting newfound mechanisms assembling ligand-free EGFR oligomer structures"

**Contents:**

**Figure S1:** Controls using single particle tracking (SPT) and confocal microscopy, related to Figure 2

**Figure S2:** Effects of tether disrupting mutations on the H2H^ect^_dimer_ and B2B^ect^_dimer_ seen in the MD simulations, related to Figure 3

**Figure S3:** Experimental results for G564P-EGFR and Lzip3A-EGFR, related to Figure 3 and Figure 4

**Figure S4:** Steps for construction of the ligand-free EGFR hetero-oligomer, related to Figure 5

**Figure S5:** 2D FLImP data and analysis for T766M-EGFR, related to Figure 5

**Figure S6:** Mutations interfere with T766M-EGFR oligomer formation, related to Figure 5 and Figure 7

**Figure S7:** Modelling and simulations of the Bb2Bb^kin^_dim_ and modelling of a “zig-zag” tetramer, related to Figure 6 and Figure 7

**Figure S8:** Growth controls in Ba/F3 cells and data supplementing the mice study, related to Figure 8

**FLImP data analysis document**

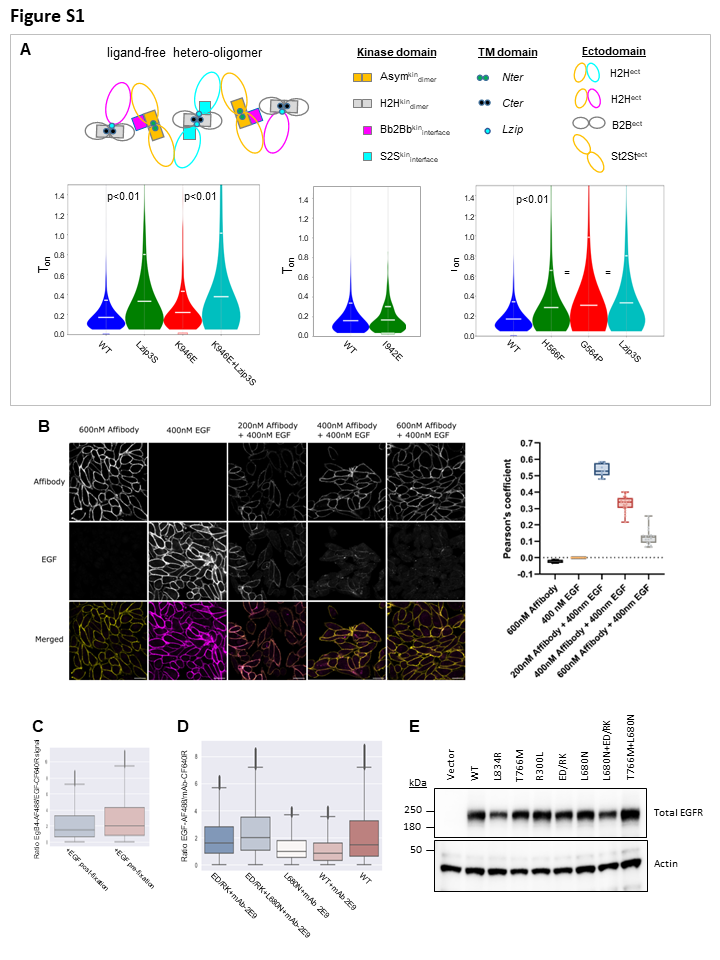

**Figure S1: Controls using single particle tracking (SPT) and confocal microscopy, related to Figure 2**

(A) *(Top)* Model of ligand-free EGFR oligomers derived from FLImP data. The proposed oligomer is assembled by 10 interfaces of different strengths, i.e., that buried different surface areas.

(*Bottom*) Use of two-colour SPT of cell surface receptors in transfected live CHO cells^1^ to check for artifacts due to cell fixation. Cells were labeled with a 1:1 mixture of anti-EGFR Affibody-Alexa 488 and Affibody-CF640R (**STAR methods**). Two colour SPT reports the incidence of pairwise particle interactions and their duration (T_ON_). T_ON_ therefore reflects the combined strength of the interactions holding the oligomer together, and thus should reflect changes in the dynamics of oligomer disassembly when mutations are introduced. Although it is hard to predict how the breaking of a large/complex oligomer in two portions might translate in changes in T_on_, a few assumptions appear reasonable:

(*Bottom* *left panel*) We would expect that the *Lzip*3S mutations will increase T_on_ when *Lzip* contacts participate in oligomer assembly. The rationale is that oligomers will more often break via the weakest assembling interaction, in this case, transmembrane contacts. Among the latter, the *Lzip* interface is not integral to any given dimer species. We therefore speculated that, in our time observation window, which is from zero to a maximum dictated by the lifetime of the fluorophores employed, the observed pairwise T_on_ should be biased towards the short duration of the *Lzip* interfaces. Thus, if the *Lzip*3S mutations are introduced, the resultant mutant oligomers will have to break via stronger interfaces, e.g., between ectodomains or kinase domains, that would be predicted to last longer. The increase in T_ON_ observed when the *Lzip*3S mutations are introduced is consistent with our speculations.

(*Bottom* *middle panel*) A second assumption safe to make refers to the I942E mutation, which strengthens the S2S^kin^_interface_. Because I942E-EGFR oligomers will still break via weak *Lzip* interfaces, I942E should not increase T_ON_, as shown by the data.

(*Bottom* *right panel*) A third assumption safe to make is that since the tether-disrupting mutations and the *Lzip*3S mutations have a similar effect on the conformation of the H2H^ect^/2x^kin^_monomers_, we would expect that the T_ON_ values for these mutations would be similar among each other, and in turn different to WT-EGFR. This is what we found. The agreement between these predictions and the two-colour SPT results backs previous results^2^ in which we could not detect that the cell fixation procedure introduced artefacts.

(B) (*Left*) Representative confocal images of cells labeled with Affibody-CF640R (top) at 4°C before fixation and EGF-Alexa488 (middle) after fixation and merged (bottom panels). Scale bars: 20 μm.

(*Right*) Numerical analysis of the overlaid colocalization images. A decrease in the Pearson colocalization coefficient clearly shows that Affibody and EGF compete for the same EGFR binding sites.

(C) Comparison of the degree of labeling with EGF-CF640R before fixation at 4^o^C and after cell fixation. Results show the fixative does not impede the binding of EGF-CF640R after cells were fixed.
(D) EGF-binding comparisons on different cell lines pre-treated with 200 nM mAb 2E9 prior to fixation and EGF labeling: A previous quantitative ^125^I-EGF binding analysis proposed that Erlotinib promotes inside-out high affinity EGF binding.^3^ In addition, Erlotinib promotes the Asym^kin^_dimer_^3^. Together, these results suggest that the St2St^ect^/Asym^kin^_dimer_ binds EGF with high affinity. To test this possibility, we labeled mAb-2E9, which selects for high affinity EGF binding^4^, with Alexa Fluor 488, labeled with 200 nM mAB-2E9-AF488, fixed them and added 200 nM EGF-CF640R, measuring the degree of EGF binding still possible after mAb-2E9 treatment by ratioing the two intensity values. The results show that mAb-2E9 blocks as much EGF binding to cells expressing WT-EGFR as to cells expressing the L680N-EGFR mutant. L680N is an N-lobe mutation of the kinase that inhibits the St2St^ect^/Asym^kin^_dimer_. In contrast, binding is not reduced when the St2St^ect^/Asym^kin^_dimer_ is present but the B2B^ect^/H2H^kin^_dimer_ has been inhibited by the ED/RK mutation. Interestingly, the results indicate that mAb-2E9 does not block the binding of EGF to the H2H^ect^/2X^kin^_monomers_.
(E) Western blot showing that the stably expressing CHO cells lines used express comparable levels of WT and receptor mutants to experiment in (**D**). Western blots from whole cell lysates of untreated CHO stable cell lines expressing different EGFR mutants, normalised for total protein content and probed with Anti-EGFR, show an equal expression level when comparing cells expressing WT-EGFR, ED/RK-EGFR and L680N-EGFR. Quantification by densitometry confirms this. The quantified intensity (in arbitrary units) of WT-EGFR peak is 23,490.61 +/- 12.036.31 (mean +/- S.D.), for ED/RK-EGFR is 22,191.42 +/- 12.137.93 and L680N-EGFR is 24,478.90 +/- 13,234.20 (mean +/- S.D.). Actin expression levels confirm equivalent total protein loading.

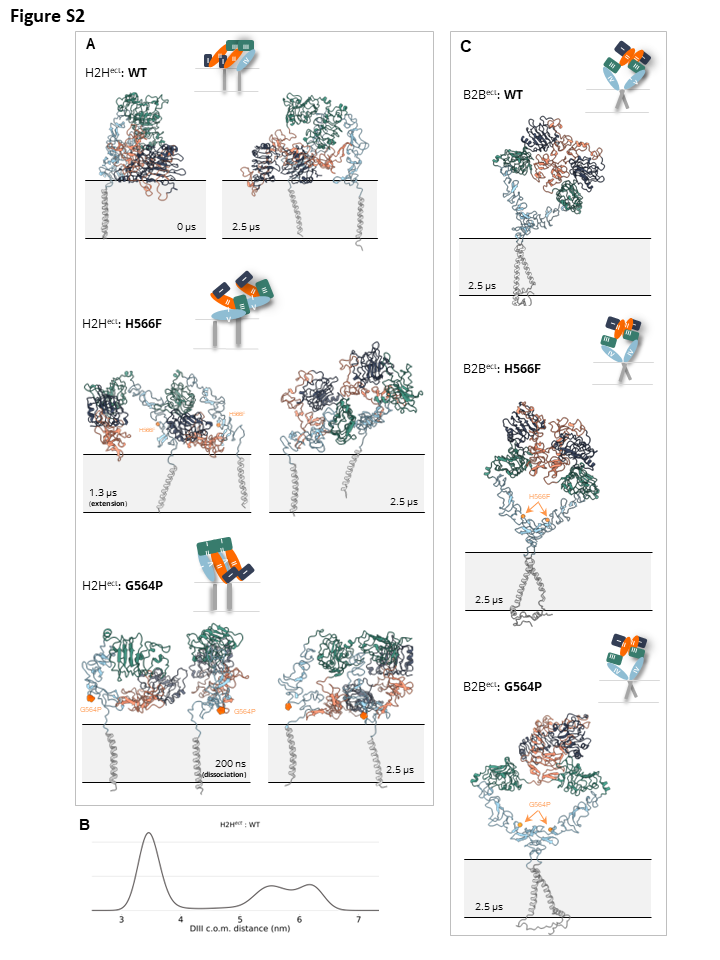

**Figure S2: Effects of tether disrupting mutations on the H2H^ect^_dimer_ and B2B^ect^_dimer_ seen in the MD simulations, related to Figure 3**

(A) Snapshots extracted from different time points of a 2.5 μs long simulation of the H2H^ect^_dimer_ and their TM domains in the lipid bilayer. The glycans bound to the ectodomain of each monomer are not shown for clarity. A cartoon representation of the dimer in the last frame of the simulation of each variant is drawn. (*Top*) Over the course of the simulation, the ectodomains (ECDs) of WT proved flexible, transitioning to a state where the ECDs oriented themselves parallel to each other and vertically to the membrane. (*Middle*) Early in the simulation, the H566F mutation removes the hydrogen bonds that H566 forms with the backbone and sidechains of T250, Y251, and Q252 from the WT DII-DIV interface. The loss of these interactions compromises the tethered conformation, and the H566F mutant adopts an open conformation transiently. Despite the transient opening, the H566F mutant is found in an H2H^ect^_dimer_ in most parts of the simulation. Unlike the WT, though, in this tethered conformation, DI has moved moves away from the membrane and closer to DIII, making the EGF binding site less accessible in both monomers. The destabilisation of the DII-DIV interface upon mutation is expected to increase the population of extended conformations that are supposed to bind EGF with higher affinity. (*Bottom*) The introduction of a Pro through the G564P mutation significantly alters the backbone configuration in the region of the DIV, which interacts with the dimerization arm on DII, thus, disrupting several intramolecular interactions that lead to the dissociation of the two monomers.

**(B)** Distribution of the centre of mass (c.o.m.) distance of the DIII domains where the fluorescent affibody binds over the course of the 2.5 μs long MD simulation of the H2H^ect^ of the WT-EGFR.

**(C)** Snapshots extracted from the last frame of a 2.5 μs long simulation of the B2B^ect^_dimer_ and their TM domains in the lipid bilayer. The glycans bound to the ectodomain of each monomer are not shown for clarity. A cartoon representation of the dimer in the last frame of the simulation of each variant is drawn. Although both monomers of the H2H^ect^_dimer_ H566F and G564P underwent larger-scale DII oscillations that triggered untethering, the monomers of the B2B^ect^_dimer_ of both variants stay in contact, and the dimers are stable over the course of the 2.5 μs long simulations.

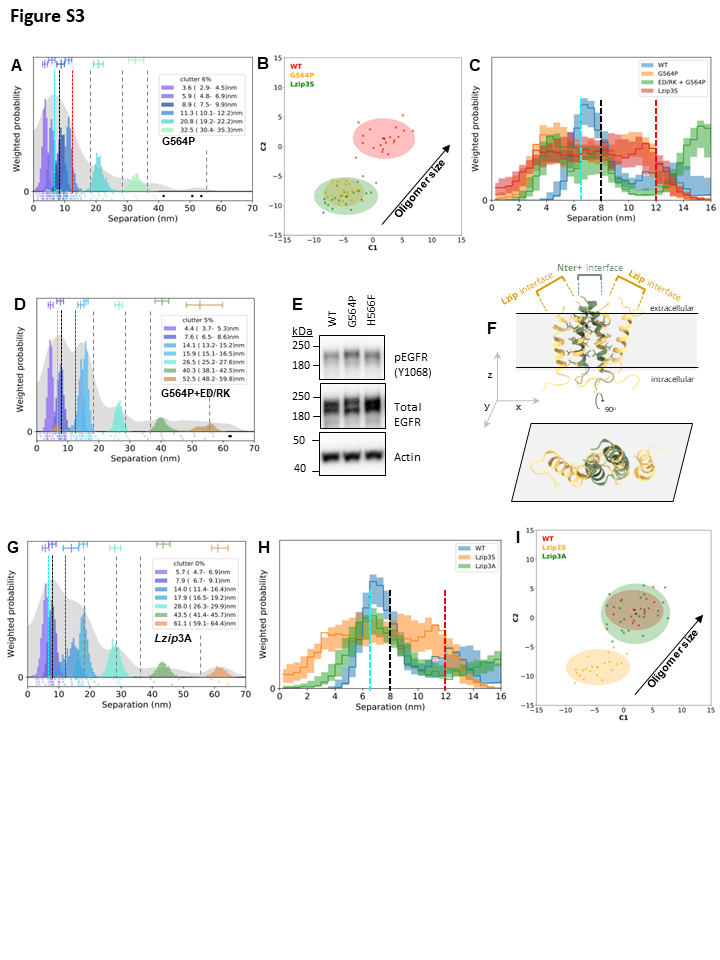

**Figure S3: Experimental results for G564P-EGFR and *Lzip3*A-EGFR, related to Figure 3 and Figure 4**

**(A, D, G)** FLImP analysis as in Fig 2.

**(B, I)** Wasserstein MDS analysis (**STAR methods**). The similarities or dissimilarities between the 21 separation sets associated to different conditions (one main FLImP decomposition plus 20 Bootstrapped with resampling decompositions) are compared; in this case, those associated to the separation sets for G564P-EGFR, Lzip3S-EGFR, and WT+EGFR. The axes in the plot are Component 1 (C1) and Component 2 (C2). C1 represents the dimension that captures the largest amount of variance in the data, while C2 represents the second-largest amount of variance that is orthogonal to C1. The centres of the 95% confidence ellipses mark the mean positions of the main FLImP decompositions. The crosses mark the positions of individual bootstrapped separation sets.

**(C, H)** Comparisons as in Figure 1(F). Note the similarity between the data for G564P-EGFR and Lzip3S-EGFR. We investigated whether the increase in separation density at ~10-12 nm was related to the B2B^ect^/H2H^kin^_dimer_ by combining the G564P and ED/RK mutations. Results show that inhibiting the latter decreases the density at this interval.

**(E)** Western blot comparing WT, G564P and H566F EGFR phosphorylation in C-terminal residue Y1086 in transfected CHO cells.

(F) (*Top*) Hexamer formed by two *Lzip* dimers interacting with a *Nter* dimer. (*Bottom*) Orthogonal projection on xy-plane.

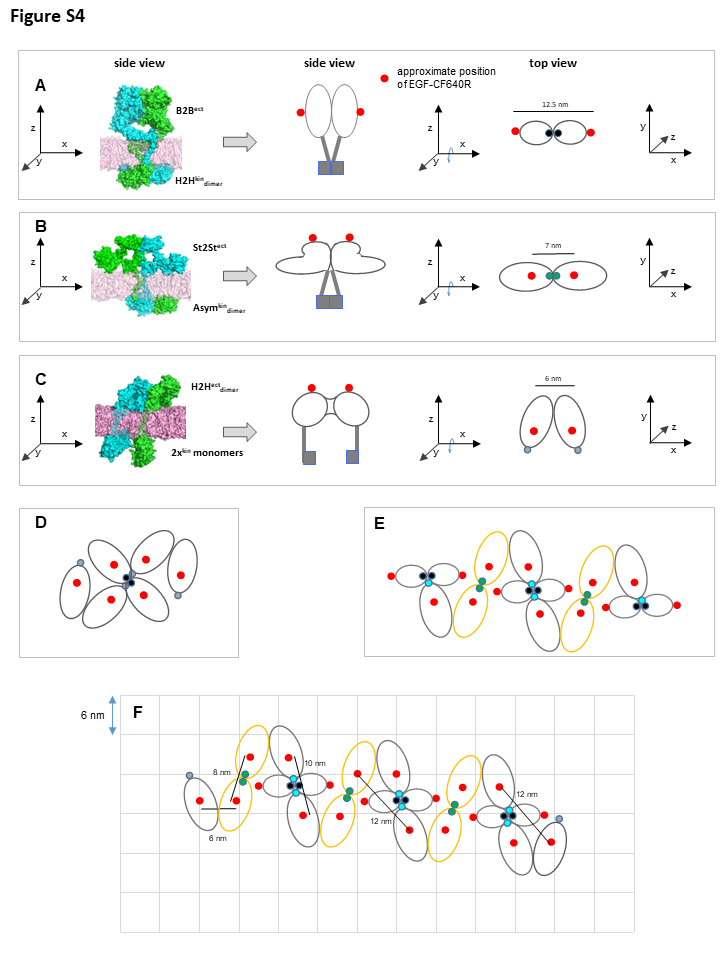

**Figure S4: Steps for construction of the ligand-free EGFR hetero^conf^-oligomer, related to Figure 5**

(A) A B2B^ect^/H2H^kin^_dimer_^5^ is simplified into a cartoon of approximate dimensions. The red dots mark the positions where we would expect the fluorescent derivative (EGF-CF640R) to bind after cells are fixed. The cartoon model of the B2B^ect^/H2H^kin^_dimer_ is then rotated around the x-axis. A top view projection of the dimer model is shown.
(B) As (A) for the St2St^ect^/Asym^kin^_dimer_^1^
(C) As (A) for the H2H^ect^/2x^kin^_monomers_^1^
(D) The attempt to connect H2H^ect^/2x^kin^_monomers_ and the St2St^ect^/Asym^kin^_dimer_ resulted in steric clashes that prevented the formation of hetero-oligomers as large as those reported by the data.

**(E, F)** Hetero^conf^-oligomers large enough to match the data could be formed by connecting H2H^ect^/2x^kin^_monomers_ and the B2B^ect^/H2H^kin^_dimer_, as shown. The model predicts the distances experimentally found by 1D and 2D FLImP. Note that in E and F the intracellular portion is not shown for simplicity.

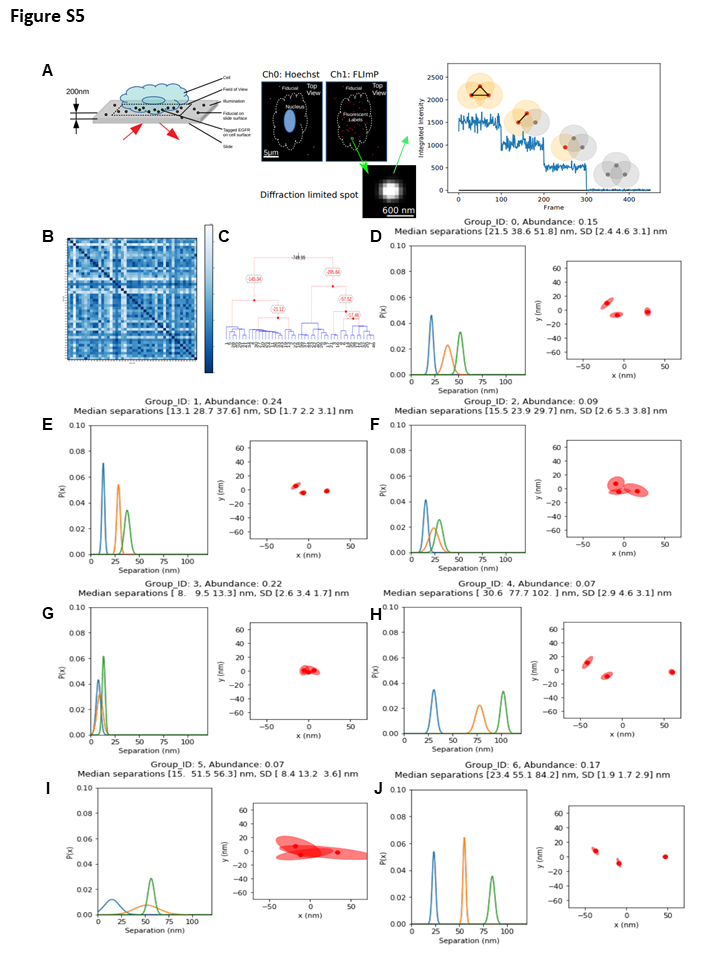

**Figure S5:** **Experimental 2D FLImP data and analysis for T766M-EGFR, related to Figure 5**

**(A)** CHO cells were imaged using total internal reflection microscopy. The cells were fixed and labeled with EGF-CF640R and Hoechst^6^, the latter to verify that data was collected on cells and not glass. Images of labelled cells were acquired, and the intensity of individual spots plotted as a function of time. Spots that decay in three steps (containing at least 3 fluorophores) were selected for further analysis.

**(B)** 2D FLImP imaging (**STAR methods**) returned a population of 46 triangles, providing information about the separations between groups of three fluorescently labeled locations in the sample of interest. Distances between each pair of 2D-FLImP posteriors were measured using the Wasserstein metric^7^, whereby identical 2D-histograms would have a Wasserstein metric of zero (blue) and more different histograms would exhibit larger values (white). As the Wasserstein metric satisfies the triangle inequality^8^, these could be used to assemble a distance matrix of triangle-relatedness where the smaller the values, the more similar the triangles.

**(C)** In turn, this was used to construct a dendrogram with optimal cuts determined using a Bayesian Hierarchical Clustering approach as described in Heller et al, ICML 2005^9^. The resulting cuts optimally grouped the 46 triangles into 7 distinct supra-triangle groups.

**(D-J)** Group members were optimally aligned before pooling and measuring separations as illustrated. Red ellipses represent 95% confidence intervals.

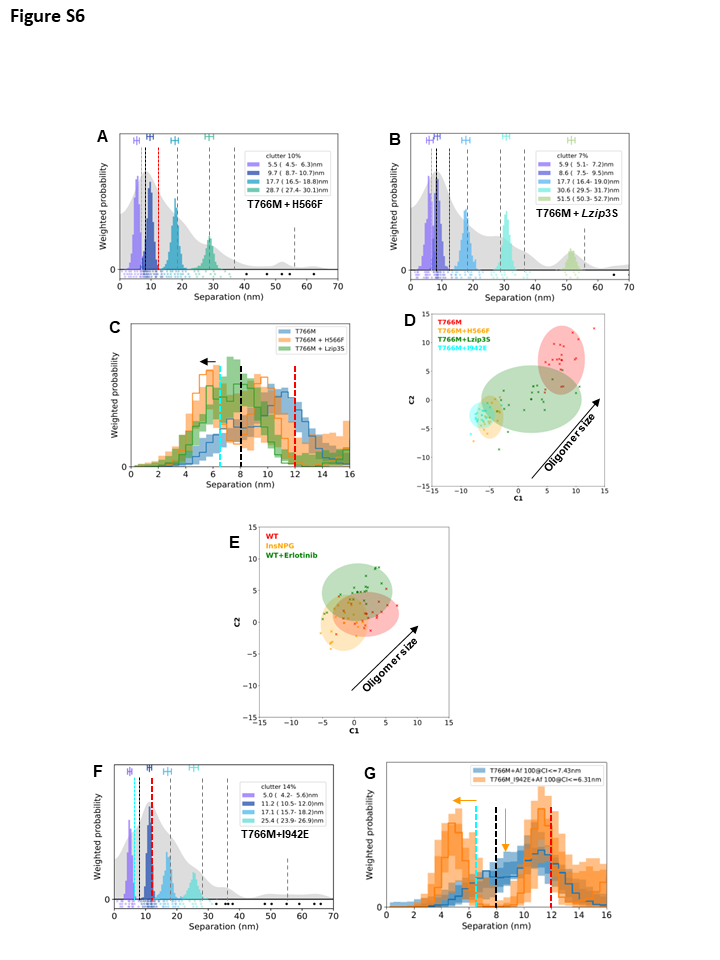

**Figure S6: The H566F, Lzip3S and I942E mutations interfere with T766M-EGFR oligomer formation, related to Figure 5 and Figure 7**

**(A, B, F)** FLImP analysis as in Fig 2.

**(C, G)** Comparisons as in Figure 1(F). Horizontal arrows mark shifts in position associated with the H2H^ect^/2x^kin^_monomers_ induced by the mutations. Vertical arrow marks a reduction in density associated with the St2St^ect^/Asym^kin^_dimer_.

**(D, E)** Wasserstein MDS analysis (**STAR methods**). The similarities or dissimilarities between the 21 separation sets associated with different conditions (one main FLImP decomposition plus 20 Bootstrapped with resampling decompositions) are compared; in this case associated with the separation sets for G564P-EGFR, Lzip3S-EGFR and WT+EGFR. The axes in the plot are Component 1 (C1) and Component 2 (C2). C1 represents the dimension that captures the largest amount of variance in the data, while C2 represents the second-largest amount of variance that is orthogonal to C1. The centre of the 95% confidence ellipses marks the positions of the main FLImP decompositions. The crosses mark the positions of individual bootstrapped separation sets.

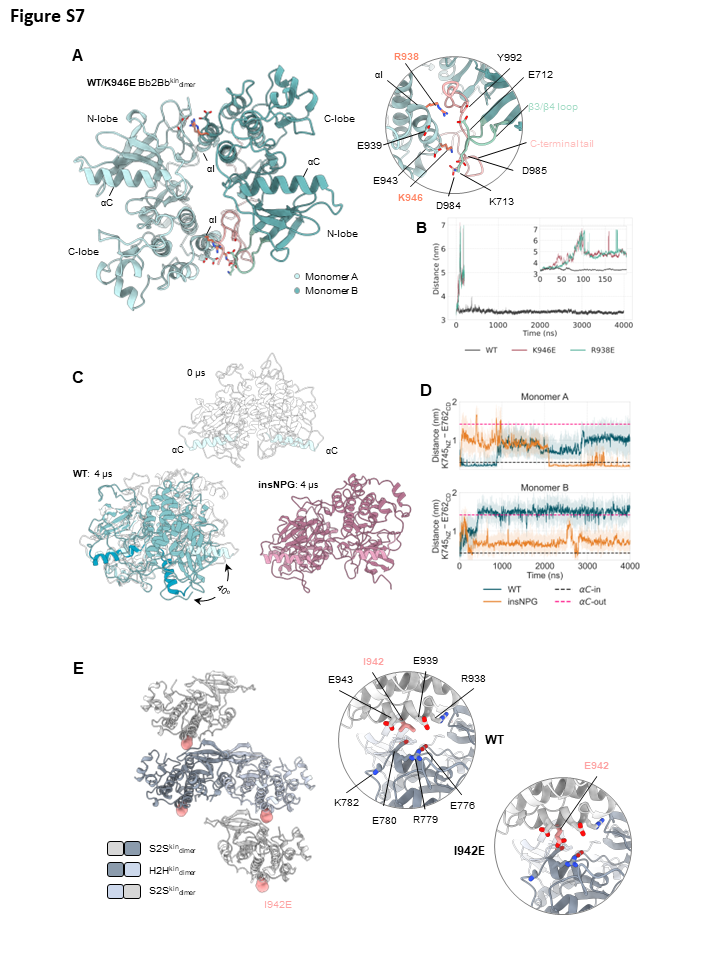

**Figure S7: Modelling and simulations of the Bb2Bb^kin^_dim_ and modelling of a “zig-zag” tetramer, relevant to Figure 6 and Figure 7.**

**(A)** (*Left*) Cartoon representation of a Bb2Bb^kin^_dimer_ (described in^10^) formed predominately through N-to-C lobe interactions (PDB ID: 3VJO^11^). (*Right*) Close-up on the interface of the Bb2Bb^kin^_dimer_ around the region where the K946E and R938E mutations lie. Both R938E and K946E are located on the αΙ-helix and are surrounded by residues that belong to flexible regions, namely the C-terminal tail and the β3/β4 loop. The presence of both positively and negatively charged residues in the vicinity of R938E and K946E already suggests that the introduction of a charge-reversal mutation within the two N-lobe/C-lobe interfaces is expected to have severe effects on the stability of the dimer.

**(B)** Time series of the center-of-mass distance between monomers A and B over the course of the 4 μs long simulations of each variant, showing the detachment of the two monomers upon K946E or R938E mutation. Within the first 100 ns, the R938E and K946E dimers broke apart, unlike the WT, which remained intact for the entire 4 μs of simulation. In the case of K946E, although K713 of the β3/β4 loop interacts at the beginning with K946E and E943, as soon as K946E gets close enough to D984 and D985 of the C-terminal tail, the repulsive potential between the negatively charged side chains disrupts the connection of the monomers in one of the N-lobe/C-lobe interfaces, pushing the two monomers away. After that, it takes only a few ns for the monomers to become flexible enough to disrupt the connection on the second interface and eventually break apart. Interestingly, K946 itself does not seem to facilitate stable inter-monomer interactions as its sidechain mostly interacts with E943 of the αI-helix. Regarding the R938E mutant, the shorter and negatively charged sidechain of R938E prevents it from maintaining inter-monomer interactions seen between R938 and the sidechains of E712 and N996 in the WT.

**(C)** Snapshots at the beginning (0 μs) and the end (4 μs) of the simulations of a WT and insNPG Bb2Bb^kin^_dimer_. The relative position of the two monomers of the insNPG remains intact over the course of the simulation, compared to the WT, in which the one monomer rotates as a rigid body about 40^o^ with respect to its initial position.

**(D)** Time series of the K745-D855 distance of each monomer of the Bb2Bb^kin^_dimer_. This distance was used as a proxy of the position of the αC-helix in the “αC-in” or “αC-out” conformation and shows the increased tendency of either monomer of the insNPG to sample αC-in conformations, even in the absence of ATP. An indicative distance of the two residues in the αC-in (PDB ID 2GS6^10^) and αC-out (PDB ID 2GS7^10^) conformation is shown with a dashed line as a reference distance.

**(E)** (*Left*) Model of a “zig-zag” tetramer formed by two monomers forming two S2S^kin^_dimer_ around an H2H^kin^_dimer_. In the crystal lattice of the kinase domain of the activator-impaired V948R EGFR (PDB ID 5CNO^12^), two different dimers can be observed; one in which the interaction between the dimers is mediated primarily by interactions between the AP-2 helix in the C-terminal tail in one kinase and the N-lobe of the other (termed H2H^kin^_dimer_), and a second one where the β2-sheet and αD-helix of the one monomer interact with the αI- and αE-helices of the other monomer (termed S2S^kin^_dimer_). The decreased phosphorylation upon I942E mutation (Figure 6D), which lies on the αI-helix of the one monomer and can form salt-bridges with R779 (αD-helix) and K782 (αG-αF loop) of the other monomer, highlights the biological relevance of the second dimer, which had been disregarded from the literature so far. The presence of these two kinds of dimers in the crystal lattice of the V948R EGFR made us speculate on the existence of a “zig-zag” kind of tetramer in cells composed of an H2H^kin^_dimer_ and two monomers attached to it in an S2S^kin^_dimer_ way. What is more, in this type of tetramer, the two monomers that form the H2H^kin^_dimer_ are found in a Src-like inactive conformation, while the two monomers attached to the H2H^kin^_dimer_ can adopt an active or inactive conformation, as their αC-helix and A-loop are not part of the S2S^kin^_dimer_ interaction interface. (*Right*) Close-up on the interface of the S2S^kin^_dimer_ around the region where the I942E lies.

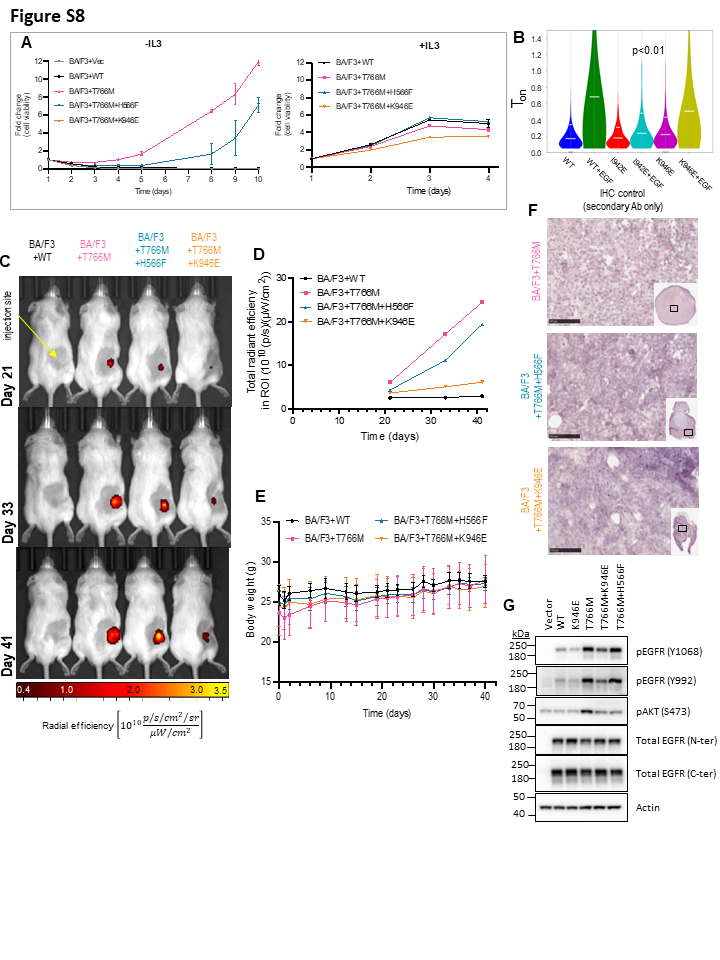

**Figure S8: Data supplementing the *in vivo* study, relevant to Figure S8.**

(A) (*Left*) The cell viability over 10 days of Ba/F3 cell stable lines expressing WT-EGFR, T766M-EGFR, T766M+K946E-EGFR, T766M+H566F-EGFR or empty vector only, in the absence of IL3. Cells were deprived of IL3 5 days prior to cell counting and seeding. Measurements of ATP levels in lysed cells were taken on the day of seeding (day 1) and up to 10 days afterwards, using the cell titre glo assay. Fold change reports the change in luminescence detected (mean +/- SD). IL3 independent proliferation is clearly only possible for the T766M-EGFR and T766M+H556F-EGFR expressing cell lines. The latter to a lesser extent. BA/F3 cells expressing T776M+K946E-EGFR, WT-EGFR or empty vector only do not proliferate without IL3. (*Right*) Positive controls showing more uniform growth among all conditions in the presence of IL3.
(B) Two-colour single particle tracking comparing T_on_ values. The data show that whilst the K946E mutation does not change the T_ON_ value either of the ligand-free or ligand-bound state, the I942E significantly decreases the T_ON_ value of the ligand-bound state.
(C) One selected tumor-bearing animal per cohort was serially imaged by IVIS on days 21, 33, and 41 after tumor inoculation. Areas around injection sites were shaved, hence the different patterns in the fur of the animals.

**(D)** Quantitative epi-fluorescence data from the animals shown in (C) as a function of time.

**(E)** Cumulative body weights of animals in all cohorts throughout the experiment indicates expected growth and no obvious adverse effects that may have been reflected in body weight loss. There were also no significant differences in animals bearing different tumor types. Error bars represent standard deviation (SD).

**(F)** Immunohistochemistry control staining of tumor sections from the same tumours as in Fig.8 with all reagents except for the specific anti-EGFR antibody. Micrographs show background staining. Scale bars are 100 μm.

**(G)** BA/F3 cells stably expressing different EGFR mutants were grown in the absence of FBS and IL3 for 2 hours followed by western blotting to assess basal EGFR and AKT phosphorylation as a measure of their activity. The impairment of the T766M+H566F mutant to sustain BA/F3 cell growth can be explained by its inability to activate a downstream signaling pathway in BA/F3 cells. In the absence of IL3 and growth factors, we found that T766M+H566F expressing BA/F3 cells were unable to phosphorylate and activate AKT signaling to the same extent as T766M.

### **FLImP data analysis**

#### **Drift determination**

Method to determine drift from a given set of tracks:

- Split tracks at points where their movement between subsequent frames is in the top 5% quantile for that frame. This will reduce the contribution from any mis-tracking, for example a track jumping between close together/overlapping spots.
- Throw out any resulting tracks which last fewer than 20 frames.
- Determine average motion of this resulting set of tracks. This is done by iteratively populating bins of aligned track positions, $\underset{̲}{\mathbf{x}}(t)=[x_{1}(t),x_{2}(t)...x_{N_{t}}(t)]$, $\underset{̲}{\mathbf{y}}(t)=[y_{1}(t),y_{2}(t)...y_{N_{t}}(t)]$, as a function of time index, $t$. $\underset{̲}{\mathbf{x}}(t)$ and $\underset{̲}{\mathbf{y}}(t)$ are initialised to the longest track, and then working through the remaining tracks in order of decreasing length, the next track is shifted in $x$ and $y$ to have the same temporal mean position as $\underset{̲}{\mathbf{x}}(t)$ and $\underset{̲}{\mathbf{y}}(t)$ at their common time points and then its shifted locations are pooled with those in $\underset{̲}{\mathbf{x}}(t)$ and $\underset{̲}{\mathbf{y}}(t)$. The final $\underset{̲}{\mathbf{x}}(t)$ and $\underset{̲}{\mathbf{y}}(t)$ will have multiple values of $x$ and $y$ at each time point from the individual aligned tracks, which are finally averaged at each time point to give an average motion, or drift, of all tracks.

To further mitigate the effect of misidentified beads, and errors in tracking (for example in crowded fields), we use a cross validation approach to identify outlier bead tracks and discard them from our drift calculation. This is performed as follows:

- Identify beads as the 200 tracks lasting as least 80% of duration whose time-averaged brightness is highest. Work only with these from here.
- Repeat 200 times:
  - Randomly reject a proportion $f_{rej}$ (=1/3) of the bead tracks, to and randomly split remaining $1-f_{rej}$ of beads into two equal partitions of tracks, $A$ and $B$.
  - Calculate drift as above for each of these partitions separately.
  - Calculate RMS residual between partition drift curves after aligning them to have the same mean location in x and y.
- Choose the partition set which gave the lowest RMS residual and pool $A$ and $B$ into one set of bead tracks. The rejected $f_{rej}$ tracks from this partition are likely to have the poorest/worst outlier tracks determined by the cross validation.
- Calculate the drift again for the pooled bead tracks to determine the final drift curve. This was repeated 100 times resampling with replacement which tracks to include to enable frame-by-frame uncertainties to be estimated.

#### **Track selection**

Each FLImP series typically returned between 1,000 and 10,000 track objects of which only a small fraction was suitable for FLImP analysis. FLImP suitable tracks were defined by the following criteria:

1. In the absence of active fluorophores, track background (the track ROI intensity in the 20 frames beyond the time at which the final fluorophore in a track goes dark) has uniform, zero intensity
2. Successive fluorophores have additive and approximately equal intensities
3. Except between sequential frames where level transitions have occurred, fluorophores are stationary in xy position, and their intensity remains constant over time. During transitions, xy positions shift by <0.5 pixels (~60nm).
4. Fluorophore intensity levels can be interrupted when fluorophores enter transitory non-radiative decay (so-called dark states) including single frame blinking events.
5. A minimum of 5 measurements is required per level for FLImP-fitting

Previously, identification of FLImP suitable tracks was a laborious process, requiring trained operators to manually trawl track lists from each FLImP series to identify tracks that may be suitable for downstream FLImP fitting processes. Recent attempts to automate similar time series analysis for single molecule imaging have relied on Bayesian grouping of localization.^13^ While such techniques are indispensable for assigning probabilistic identities to large groups of localizations, as the present 2D FLImP technique typically involves fewer than 10 fluorophores per track object, this enabled simpler, and substantially faster, sequential filtering approach to be used. Such an approach was chosen over emerging 1D convolutional neural networks (1D-CNNs)^14^ owing to the tractability of the decision-making process and the lack of a sufficiently large manually labeled training dataset. That said, the outputs of the following sequential filtering approach have been written with this future development in mind by accelerating the generation of labeled training datasets and will be the subject of future work. The authors anticipate that progressing to a 1D-CNN approach is likely to increase the efficiency of FLImP suitable track identification, but at least in the short term this will come at the price of explainability which may limit the ultimate clinical utility of FLImP derived technologies should this path be taken.

The sequential filtering approach to track selection involves passing a population of tracks through a series of filters to rapidly and automatically identify tracks suitable for further FLImP analysis. As tracks with increasing numbers of fluorophores represent nested sets, track subsets may be suitable for further FLImP analysis, even if the entire track is not. As such, a successful automated FLImP suitable track identification algorithm is required to evaluate track suitability at different granularities.

Filters are organized so that the most computationally resource intensive filters are located towards the end of the pipeline, to minimize resource requirement and accelerate this aspect of the process. As such, filtering of a typical FLImP series containing 5000 track objects is typically completed within as little as 60 seconds using a typical single CPU core and lends itself to parallelization. Furthermore, the track selection process has been written so that it can be applied to tracks containing any up to 9 fluorophores (the anticipated limit for a non-Bayesian approach). A detailed description of each filter is provided in the following sections. Filter parameters were developed heuristically on training datasets independent of those presented in the results section of this manuscript and were then held constant throughout.

1. Label levels - Tracks were first divided into crude intensity levels using the R implementation of the dynamically programmed Optimal k-means clustering algorithm for one dimensional data^15^ and the level with the lowest median intensity assigned background (Level 0) and levels with increasing median intensity labeled in ascending numerical order.
2. Ensure zero background - Remove tracks where the best-fit spot intensity at final location in the track is not zero for the 20 frames after the final time point as determined using a Two One-Sided Test (TOST) test.^16^ The limits of the test were set using 10\% of the 95th percentile of track intensity and (ɑ=0.05).
3. Remove short levels - Next, levels were removed from tracks that contained fewer than 5 frames.
4. Ensure adequate level intensity separations - Remove tracks with median level intensity separations that are not 2*(level) fold different from the intensity separation between level 1 and background (threshold +- 0.25). This is because we expect fluorophores to have very similar and additive intensities within tracks (but this can vary between tracks).
5. Short inter-level sub-level removal - Sub-levels, defined as sections of track that occur between level-change events whose median intensity is not an integer multiple of the lowest intensity level are identified and those containing fewer than 5 frames are removed from further analysis. These may correspond to either blink states that extend over multiple frames or level transitions that are spread over multiple frames.
6. Intra-level sub-level removal - Intra-level sub-levels denote the existence of intensity or positional transitions that can be observed within a single level. These were isolated using R change point implementation of the PELT (Pruned Exact Linear time) algorithm to independently detect potential changes of state within each fluorophore level.^17^
7. Intra-level sub-level positional filter - Intra-level sub-levels are further filtered using Median absolute deviation (MAD)^18^ in intensity and x,y positions to remove sub-tracks with MAD >10% of median level intensity and 0.125 pixels respectively as measurements such as these were deemed likely to be too imprecise for FLImP fitting. Background levels were excluded from this filter.
8. Remove sub-levels with significant intensity gradients and low occupancy - Intra-level and sub-levels containing fewer than 5 frames or with an occupancy (proportion of frames where a spot was detected within the level) <80% were then removed, before all intra-level sub-levels with a significantly non-zero gradient (TOST test >5% gradient where ɑ=0.05). Background level was excluded from this filter.
9. Intra-level sub-level median intensity equivalence - Ensure all intra-level sub-levels have the same median intensity +/- 12.5% using a TOST test. This is to ensure levels interrupted by brief transitions between states (i.e. blink states) return to near identical starting conditions so can be classed as the same object. Such level merging increases the efficiency of the track selection process.
10. Inter-level positional shift filter - Remove all sub-tracks that show excessive x or y positional shifts (median positional shift >0.125 pixels for any sub-track within a level). Excluding background levels.
11. Minimum frames and levels filter - As quite a few filters have been applied to sub-tracks within the data, ensure that all remaining tracks still contain sufficient frames to be useful for bootstrapping and that all desired sequential levels are still sufficiently represented, depending on the maximum number of levels currently under evaluation. For example, level Background + 1 + 2 for two spot tracks or level Background + 1 + 2 + 3 for three spot tracks).
12. Spatial neighborhood filter - Remove frames from tracks that are within ~2 PSF (approximately 5 pixels) of another detected fluorophore at any time point. This is because these neighboring spots objects are likely to contribute to inhomogeneity in the background of the spot of interest (which is currently assumed to be constant that can vary between frames). This is achieved by calculating the cross-nearest neighbor distance for each frame in the selected track for every detected object in the frame.
13. Inter-level minimum frames filter - Remove sequential sub tracks that contribute fewer than 5 frames to the track dataset as these were found to be of low quality (i.e., temporally sparse, from small levels etc).
14. Identify and remove blinking events - Blinking events, typically single or double frames events that exhibit intensities that are substantially lower than the ordinary intensity distribution of a level. Assuming fluorophore intensity is drawn from a sum of normal distributions for each fluorophore present (n-spots * N(μ,σ)), blinking events can be considered as outliers from this distribution. As such, a modified Z-score,^19^ can be used to readily identify and remove outlying intensity, (and localization x and y values) from remaining tracks using the threshold of z<3.
15. Maximum level filtering and minimum frames per level filtering - Tracks are filtered so that only those containing the desired number of levels are retained. As quite a few filters have been applied to sub-tracks within the data, ensure that all remaining tracks still contain sufficient frames (5 per level) to be useful for bootstrapping and that all desired sequential levels are still sufficiently represented. (level 1 + 2 or level 1 + 2 + 3).
16. Split-level matching - To recombine split levels, we need confidence that they are drawn from the same location, here defined as returning a TOST test result with limits of +/- 0.1 pixels of x and y where ɑ = 0.05. In rare instances where split levels cannot be merged (i.e., two fluorophores simultaneously switch on and off at the same point), the fluorophore containing the most frames is used.
17. Level ordering filter - Split levels of the type lvl1-lvl2-lvl1 within a track can be indicative of two fluorophores (A+B) behaving in one of two ways. A-AB-A or A-AB-B. In the second case, it would be undesirable to merge the two lvl1 subsets. As such, tracks are filtered such that only those with sequentially decreasing levels are selected.
18. MAD population intensity filter - A limitation of estimating the intensity of a single fluorophore within each track independently is that this method will not distinguish between situations where multiple fluorophores go dark at once. For example, the two step tracks; for the two-fluorophore system: AB-B-0 and the four-fluorophore system: ABCD-AB-0 may both exhibit 2N-N-0 intensity steps. While the second case is likely to be rare, this eventuality can only be eliminated by considering the fluorophore intensity at the population scale. Here, the intensity of each fluorophore is pooled and tracks excluded when fluorophores were found to possess MAD values exceeding 2.5 of the fluorophore population.

#### **Separation measure**

The difference in position, or separation, between locations is normally represented by the L2 norm between the locations. When those locations are uncertain, the probability distribution for their separation also has uncertainty and can take forms which are problematic for measuring short separations in particular. Measuring separations between uncertain locations is a common task in single molecule imaging. In the simplest case, where the uncertainty in each location can be represented by the same axisymmetric 2D normal distribution, the probability distribution for the L2 separation will be a Rician distribution. In the extreme case that the locations are coincident, i.e., the true separation is zero, the Rician distribution will have zero probability at separation zero. In general, using the L2 norm means low sensitivity and possible biases in measurement of separations of order the location uncertainty or less. In traditional 1D-FLImP (which used Ricean modelling of the L2 norms) this meant separations $<$8nm could not be accurately quantified.

We have used an alternative measure in some areas our new FLImP analysis which we call $r_{\Delta x}$. $r_{\Delta x}$ is calculated as follows:

1. A pair of emitters, A and B, have locations in space $\underset{̲}{\mathbf{R}_{A}}$ and $\underset{̲}{\mathbf{R}_{B}}$ described by 2D probability distributions $\Pr(\underset{̲}{\mathbf{R}_{A}})$ and $\Pr(\underset{̲}{\mathbf{R}_{B}})$ respectively (e.g., from the 2400 FLImP bootstrap location samples for each emitter in the track object.
2. The $\underset{̲}{\mathbf{L}}$ direction is defined as the direction from the estimated location of A, $\langle\underset{̲}{\mathbf{R}_{A}}\rangle$ to the estimated location of B, $\langle\underset{̲}{\mathbf{R}_{B}}\rangle$, i.e. $L=(\langle\underset{̲}{\mathbf{R}_{B}}\rangle-\langle\underset{̲}{\mathbf{R}_{A}}\rangle)/|\langle\underset{̲}{\mathbf{R}_{B}}\rangle-\langle\underset{̲}{\mathbf{R}_{A}}\rangle|$.
3. The separation measure used, $r_{\Delta x}$, between two points $\underset{̲}{\mathbf{r}_{A}}$ and $\underset{̲}{\mathbf{r}_{B}}$ chosen from emitter posteriors $\Pr(\underset{̲}{\mathbf{R}_{A}})$ and $\Pr(\underset{̲}{\mathbf{R}_{B}})$ respectively is the vector between those two points resolved parallel to $\underset{̲}{\mathbf{L}}$, i.e. $r_{\Delta x}=(\underset{̲}{\mathbf{r}_{B}}-\underset{̲}{\mathbf{r}_{A}}).\underset{̲}{\mathbf{L}}$. $r_{\Delta x}$ can be positive or negative, and has the desirable property that in the event that $\Pr(\underset{̲}{\mathbf{R}_{A}})$ and $\Pr(\underset{̲}{\mathbf{R}_{B}})$ are axisymmetric Gaussians the probability distribution of $r_{\Delta x}$ between A and B, $\Pr(r_{\Delta x})$, will be a Gaussian centred on $|\underset{̲}{\mathbf{R}_{B}}-\underset{̲}{\mathbf{R}_{A}}|$. It does not suffer from the insensitivity or bias at short separations suffered by the L2 norm and can in principle even measure zero separations.
4. Normal distribution-based approximations to $\Pr(r_{\Delta x})$ are used in our 1D-FLImP decomposition which allows fast evaluation.

#### **1D decomposition**

##### FLImP measurements as posteriors

We have $n$ spots (tracks) (labeled $i=1..n$) for each of which we have measured a single separation from the FLImP localization fit. We interpret the bootstrap distribution for separation $x$ from spot $i$ given its data $D_{i}$ as a posterior for the separation, $\text{Pr}\left( x|D_{i} \right)$. From Bayes’ Theorem,

$$\text{Pr}\left( x|D_{i} \right)=\frac{\text{Pr}\left( D_{i}|x \right)\text{Pr}\left( x \right)}{\text{Pr}\left( D_{i} \right)}$$

If we limit $x$ to the range 0 to $x_{max}$ with a uniform prior, then in this range we have

$$\text{Pr}\left( x \right)=\frac{1}{x_{max}}$$

We can then write the likelihood for $D_{i}$ given $x$,

$$\text{Pr}\left( D_{i}|x \right)=\text{Pr}\left( x|D_{i} \right)\text{Pr}\left( D_{i} \right)\left( x_{max} \right)=a_{i}\text{Pr}\left( x|D_{i} \right)$$

Equation 1

where $a_{i}=x_{max}\text{Pr}\left( D_{i} \right)$.

##### Decomposition model

For our model we assume the structure being measured has $N$ distinct components, $k=1..N$, each component having a different separation, $\underline{\mathbf{x}}=\{x_{1},..,x_{N}\}$. Each FLImP separation corresponds to one of these components or is a spurious “clutter” measurement. This assignment for each measurement is denoted by $\underline{\mathbf{K}}=\{k_{1},..k_{n}\}$, $k_{i}=1..N$, where measurement $i$ corresponds to component $k_{i}$ with separation $x_{k_{i}}$, and $k_{i}=0$ means measurement $i$ is clutter.

Assuming the model components are precise, and the separation of clutter components is uniformly distributed in the $x$ domain, then the probability distribution of separation for component $k$ is

$$\text{Pr}\left( x|k \right)=\left\{ \begin{matrix} \delta\left( x-x_{k} \right) & \text{if }k>0 \\ 1/\Delta x & \text{if }k=0\text{ and }x\text{ is within the domain} \\ 0 & \text{if }k=0\text{ and }x\text{ is not within the domain} \end{matrix} \right.$$

Equation 2

where $\delta$ is the Dirac delta function and $\Delta x$ is the size of the domain of $x$.

We assume that the probability of the measurement process yielding a clutter measurement is $P_{c}$, so that typically this should be the proportion of measurements which is clutter. We assume that the non-clutter components are all equally probable.

We therefore have model parameters $\{\underline{\mathbf{K}},\underline{\mathbf{x}},P_{c}\}$.

##### Model likelihood and posterior

Each spot measurement is completely independent, so that the likelihood for the entire set of measurements $\underline{\mathbf{D}}=\{D_{1},..,D_{n}\}$ given $\underline{\mathbf{x}}$ and $\underline{\mathbf{K}}$ is

$$\text{Pr}\left( \underline{\mathbf{D}}|\underline{\mathbf{K}},\underline{\mathbf{x}} \right)=\prod_{i=1}^{n} \text{Pr}\left( D_{i}|k_{i},\underline{\mathbf{x}} \right)$$

We have $\text{Pr}\left( D_{i}|x \right)$ from **Equation 1** and $\text{Pr}\left( x|k \right)$ from **Equation 2** which we use to calculate $\text{Pr}\left( D_{i}|k_{i},\underline{\mathbf{x}} \right)$. $P_{c}$ does not figure here as once $\underline{\mathbf{x}}$ and each $k_{i}$ are given the likelihood of the data is known.

$$\begin{matrix} \text{Pr}\left( D_{i}|k_{i},\underline{\mathbf{x}} \right) & = & \int_{-\infty}^{\infty} \text{Pr}\left( D_{i}|x \right)\text{Pr}\left( x|k_{i} \right)dx \\ & & \\ & = & \left\{ \begin{matrix} \int_{-\infty}^{\infty} \text{Pr}\left( D_{i}|x \right)\delta\left( x-x_{k_{i}} \right)dx & = & \text{Pr}\left( D_{i}|x=x_{k_{i}} \right) & \\ & = & a_{i}\text{Pr}\left( x_{k_{i}}|D_{i} \right) & \text{if }k_{i}>0 \\ & & & \\ \int_{-\infty}^{\infty} \text{Pr}\left( D_{i}|x \right)/\Delta xdx & = & \int_{-\infty}^{\infty} a_{i}\text{Pr}\left( x|D_{i} \right)/\Delta xdx & \\ & = & a_{i}/\Delta x & \text{if }k_{i}=0 \end{matrix} \right. \end{matrix}$$

We abbreviate $L_{k_{i}}=\text{Pr}\left( D_{i}|k_{i},\underline{\mathbf{x}} \right)/a_{i}$ so that

$$\begin{matrix} L_{k_{i}} & = & \left\{ \begin{matrix} \text{Pr}\left( x_{k_{i}}|D_{i} \right) & \text{if }k_{i}>0 \\ 1/\Delta x & \text{if }k_{i}=0 \end{matrix} \right. \end{matrix}$$

Equation 3

so that

$$\text{Pr}\left( \underline{\mathbf{D}}|\underline{\mathbf{K}},\underline{\mathbf{x}} \right)=\prod_{i=1}^{n} \text{Pr}\left( D_{i}|k_{i},\underline{\mathbf{x}},P_{c} \right)=\prod_{i=1}^{n} a_{i}L_{k_{i}}=A\prod_{i=1}^{n} L_{k_{i}}$$

where $A=\prod_{i=1}^{n} a_{i}$.

From Bayes theorem we have:

$$\text{Pr}\left( \underline{\mathbf{K}},\underline{\mathbf{x}}|\underline{\mathbf{D}} \right)=\frac{\text{Pr}\left( \underline{\mathbf{D}}|\underline{\mathbf{K}},\underline{\mathbf{x}} \right)\text{Pr}\left( \underline{\mathbf{K}},\underline{\mathbf{x}} \right)}{\text{Pr}\left( \underline{\mathbf{D}} \right)}=\frac{\text{Pr}\left( \underline{\mathbf{D}}|\underline{\mathbf{K}},\underline{\mathbf{x}} \right)\text{Pr}\left( \underline{\mathbf{x}} \right)}{\text{Pr}\left( \underline{\mathbf{D}} \right)}=\frac{\text{Pr}\left( \underline{\mathbf{x}} \right)}{\text{Pr}\left( \underline{\mathbf{D}} \right)}A\prod_{i=1}^{n} L_{k_{i}}$$

as the prior $\text{Pr}\left( \underline{\mathbf{K}},\underline{\mathbf{x}} \right)=\text{Pr}\left( \underline{\mathbf{K}} \right)\text{Pr}\left( \underline{\mathbf{x}} \right)=\text{Pr}\left( \underline{\mathbf{x}} \right)=\prod_{k=1}^{N} \text{Pr}\left( x_{k} \right)=\prod_{k=1}^{N} \pi_{x_{k}}$, $\underline{\mathbf{K}}$ is only constrained through $\text{Pr}\left( \underline{\mathbf{K}}|P_{c},\underline{\mathbf{D}} \right)$ (see below) so $\text{Pr}\left( \underline{\mathbf{K}} \right)=1$ and we assume a uniform prior on $x$ within the domain

$$\text{Pr}\left( x_{k} \right)=\pi_{x_{k}}=\left\{ \begin{matrix} 1/\Delta x & \text{if }x\text{ in domain} \\ 0 & \text{if }x\text{ outside domain} \end{matrix} \right.$$

Our model for clutter gives a probability for the assignments, $\text{Pr}\left( \underline{\mathbf{K}}|P_{c},\underline{\mathbf{D}} \right)$. This is composed of two components as follows.

First, the number of measurements assigned to clutter, $n_{c}=\sum_{i=1}^{n} \delta_{k_{i},0}$ where $\delta$ is the Kronecker delta, will be binomially distributed with probability $P_{c}$, i.e.

$$\text{Pr}\left( n_{c} \right)=\frac{n_{c}!}{n!\left( n-n_{c} \right)!}P_{c}^{n_{c}}\left( 1-P_{c} \right)^{n-n_{c}}$$

Secondly, the assumption that the clutter is uniformly distributed in the $x$ domain can be expressed as a multinomial prior on the number of measurements assigned to clutter in a set of $B$ equal-sized bins $b=1..B$ in the $x$ domain, $\underline{\mathbf{n}_{c_{b}}}=\{n_{c_{1}}..n_{c_{B}}\}$. We have $n_{c_{b}}=\sum_{i=1}^{n} \delta_{k_{i},0}F\left( b,\hat{x_{i}} \right)$ where

$$F\left( b,x \right)=\left\{ \begin{matrix} 1 & \text{if }\left( b-1 \right)\Delta x/B\leq x<b\Delta x/B \\ 0 & \text{otherwise} \end{matrix} \right.$$

and $\hat{x_{i}}$ is $\left\langle\text{Pr}\left( r_{\Delta x}|D_{i} \right) \right\rangle$, the mean $r_{\Delta x}$ for measurement $i$. We use $B=20$ bins. So

$$\text{Pr}\left( \underline{\mathbf{n}_{c_{b}}} \right)=\frac{n_{c}!}{\prod_{b=1}^{B} n_{c_{b}}!}\prod_{b=1}^{B} \left( 1/B \right)^{n_{c_{b}}}$$

This gives:

$$\text{Pr}\left( \underline{\mathbf{K}}|P_{c},\underline{\mathbf{D}} \right)=\left\{ \begin{matrix} \text{Pr}\left( n_{c} \right) & \text{if }n_{c}=0 \\ \text{Pr}\left( n_{c} \right)\text{Pr}\left( \underline{\mathbf{n}_{c_{b}}} \right) & \text{if }n_{c}>0 \end{matrix} \right.$$

The posterior probability can be separated into two components,

$$\text{Pr}\left( \underline{\mathbf{K}},\underline{\mathbf{x}},P_{c}|\underline{\mathbf{D}} \right)=\text{Pr}\left( \underline{\mathbf{K}},\underline{\mathbf{x}}|\underline{\mathbf{D}} \right)\text{Pr}\left( \underline{\mathbf{K}},P_{c}|\underline{\mathbf{D}} \right)$$

.

We can write

$$\text{Pr}\left( \underline{\mathbf{K}},P_{c}|\underline{\mathbf{D}} \right)=\text{Pr}\left( \underline{\mathbf{K}}|P_{c},\underline{\mathbf{D}} \right)\text{Pr}\left( P_{c}|\underline{\mathbf{D}} \right)=\text{Pr}\left( \underline{\mathbf{K}}|P_{c},\underline{\mathbf{D}} \right)\text{Pr}\left( P_{c} \right)=\text{Pr}\left( \underline{\mathbf{K}}|P_{c},\underline{\mathbf{D}} \right)\pi_{P_{c}}$$

since the clutter probability does not depend on the data. For the clutter fraction prior we assume a beta distribution $\text{Pr}\left( P_{c} \right)=\pi_{P_{c}}=\text{Beta}\left( P_{c},\alpha,\beta\right)$ with shape parameters $\alpha=1$, $\beta=9$ which peaks at $P_{c}=0.0$ and has $\left\langle\pi_{P_{c}} \right\rangle=0.1$.

We therefore have the full posterior

$$\text{Pr}\left( \underline{\mathbf{K}},\underline{\mathbf{x}},P_{c}|\underline{\mathbf{D}} \right)=\frac{\text{Pr}\left( \underline{\mathbf{K}}|P_{c},\underline{\mathbf{D}} \right)\pi_{P_{c}}\text{Pr}\left( \underline{\mathbf{D}}|\underline{\mathbf{K}},\underline{\mathbf{x}} \right)\text{Pr}\left( \underline{\mathbf{x}} \right)}{\text{Pr}\left( \underline{\mathbf{D}} \right)}=\frac{\text{Pr}\left( \underline{\mathbf{K}}|P_{c},\underline{\mathbf{D}} \right)\pi_{P_{c}}}{\text{Pr}\left( \underline{\mathbf{D}} \right)}A\prod_{k=1}^{N} \pi_{x_{k}}\prod_{i=1}^{n} L_{k_{i}}$$

Equation 4

##### Sampling from the posterior

Our model parameters are separations $\underline{\mathbf{x}}$, clutter probability $P_{c}$ and assignments $\underline{\mathbf{K}}$. We wish to draw a representative set of samples of these from the posterior $\text{Pr}\left( \underline{\mathbf{K}},\underline{\mathbf{x}},P_{c}|\underline{\mathbf{D}} \right)$. We do this using the Metropolis-Hastings Markov Chain Monte Carlo (MCMC) sampling approach, sampling parameters in turn as follows:

1. Initialise parameters at iteration $t=0$. Each component of $\underline{\mathbf{x}}\left( t=0 \right)$ is sampled independently from $\pi_{x_{k}}$, $P_{c}\left( t=0 \right)$ is sampled from the clutter prior $\pi_{P_{c}}$ and each component $k_{i}$ of assignment $\underline{\mathbf{K}}\left( t=0 \right)$ is sampled independently from $0..N$ with a uniform distribution.
2. For each iteration $t\to t+1$:
   a. Metropolis-Hastings sample $P_{c}\left( t+1 \right)$ given $\underline{\mathbf{x}}\left( t \right)$ and $\underline{\mathbf{K}}\left( t \right)$ . The proposal distribution for $P_{c}\left( t+1 \right)$ is $\mathcal{N}\left( P_{c}\left( t \right), \sigma_{P_{c}}^{2} \right)$ limited to the range $0..1$ with $\sigma_{P_{c}}=0.1$.
   b. Jointly Metropolis-Hastings sample $\underline{\mathbf{x}}\left( t+1 \right)$ and $\underline{\mathbf{K}}\left( t+1 \right)$ given $P_{c}\left( t+1 \right)$ by Metropolis-Hastings sampling each component $x_{k}$ of $\underline{\mathbf{x}}$ ($k=1..N$) in turn, jointly with $\underline{\mathbf{K}}$. Perform $j=1..N$ sub-iterations yielding in turn updated parameters $\underline{\mathbf{x}}'\left( j \right)$ and $\underline{\mathbf{K}}'\left( j \right)$, starting with $\underline{\mathbf{x}}'\left( 0 \right)=\underline{\mathbf{x}}\left( t \right)$ and $\underline{\mathbf{K}}'\left( 0 \right)=\underline{\mathbf{K}}\left( t \right)$, with each iteration as follows:
      i. Proposal for $\underline{\mathbf{x}}'\left( j \right)$ is ${\underline{\mathbf{x}}}^{*}$, with component $x_{k=j}^{*}$ from proposal distribution $Q\left( x_{k=j}^{*}|x_{k=j}\left( t \right) \right)$ and components $x_{k\neq j}^{*}=x'_{k}\left( j-1 \right)$. $Q\left( x'|x \right)=0.5*P_{\mathrm{DATA}}\left( x' \right)+0.5*\left( w_{1}\mathcal{N}\left( x'-x, \sigma_{1}^{2} \right)+w_{2}\mathcal{N}\left( x'-x, \sigma_{2}^{2} \right) \right)$ with $\left( w_{1},\sigma_{1},w_{2},\sigma_{2} \right)=\left( 0.7,2\text{nm},0.3,5\text{nm} \right)$. $P_{\mathrm{DATA}}\left( x \right)=\frac{1}{n}\sum_{i=1}^{n} \text{Pr}\left( x|D_{i} \right)$ is the normalised sum of sample data posteriors (see **FLImP measurements as posteriors**).
      ii. Proposal ${\underline{\mathbf{K}}}^{*}$for $\underline{\mathbf{K}}'\left( j \right)$ has each component directly sampled from **Equation 6** given the proposed ${\underline{\mathbf{x}}}^{*}$.
      iii. Metropolis-Hastings sample to accept/reject a move from $\underline{\mathbf{x}}'\left( j-1 \right)$, $\underline{\mathbf{K}}'\left( j-1 \right)$, $P_{c}\left( t+1 \right)$ to $\underline{\mathbf{x}}'\left( j \right)$, $\underline{\mathbf{K}}'\left( j \right)$, $P_{c}\left( t+1 \right)$. Set $\underline{\mathbf{x}}'\left( j \right)$, $\underline{\mathbf{K}}'\left( j \right)$ to $\underline{\mathbf{x}}'\left( j-1 \right)$, $\underline{\mathbf{K}}'\left( j-1 \right)$ if the move is rejected, and to ${\underline{\mathbf{x}}}^{*}$, ${\underline{\mathbf{K}}}^{*}$ if it is accepted.
   b. After the sub-iterations we keep $\underline{\mathbf{x}}\left( t+1 \right)=\underline{\mathbf{x}}'\left( N \right)$ and $\underline{\mathbf{K}}\left( t+1 \right)=\underline{\mathbf{K}}'\left( N \right)$.

Jointly sampling $\underline{\mathbf{x}}$ components and $\underline{\mathbf{K}}$ this way ensures a sufficient acceptance rate by proposing changes in assignment commensurate with changes in component locations.

Each new $\underline{\mathbf{x}}$ sample is sorted in ascending order.

Sampling is performed until convergence, and samples are thinned to keep every 20th sample (autocorrelation tests on sample chains reveal this thinning to be sufficient). The first 50% of samples are discarded to exclude the burn-in period. Two independent chains of samples are run, each initialized separately, and convergence is checked periodically after each chain has drawn the same number of samples comparing the second 50% of thinned samples in each chain using the Gelman-Rubin test. The test is initially performed after 2000 iterations (giving 100 post-burnin thinned samples). If the test fails sampling resumes to increase the iterations by 50% (i.e., to 3000, 4500... iterations). The sampling fails if convergence isn’t reached after 40000 iterations (10000 post-burnin thinned samples). The chains are considered to have converged if $\left| 1-\hat{R} \right|<0.001$ for each of $x_{k}$ ($k=1..N$) and $P_{c}$ compared separately between the two chains (A and B), where $\hat{R}=\sqrt{\frac{\hat{\sigma}^{2}}{W}}$ is the univariate Gelman-Rubin statistic^20^ for each parameter, and

$$W=\frac{\sigma_{A}^{2}+\sigma_{B}^{2}}{2}$$

$$B=n_{iter}\left( \left( \mu_{A}-\mu\right)^{2}+\left( \mu_{B}-\mu\right)^{2} \right)$$

$$\hat{\sigma}^{2}=\frac{n_{iter}-1}{n_{iter}}W+\frac{1}{n_{iter}}B$$

with $n_{iter}$ samples used from each chain, the variance of the samples from each chain is $\sigma_{A/B}^{2}$ respectively, the mean of the samples in each chain is $\mu_{A/B}$ and $\mu=\left( \mu_{A}+\mu_{B} \right)/2$. Of the two chains for each $N$ we adopt the one with the lowest $AVBIC$ (see **Model selection**) for further analysis (they are almost identical in all cases given the stringent convergence criterion).

We use $0..\Delta x$ as the domain of our separation measurement, with $\Delta x$ set so that all of the probability in $P_{\mathrm{DATA}}\left( x \right)$ is in the domain.

We use $x=\left| r_{\Delta x} \right|$ (see **Separation measure**) for our separation measure in the decomposition. For computational efficiency we replace each measurement posterior $\text{Pr}\left( r_{\Delta x}|D_{i} \right)$ with a Gaussian approximation (from its mean and variance).

##### Assignment proposal distribution

While the assignments do have a joint posterior due the inclusion of clutter probability in the model, we ignore this and treat each measurement separately when calculating the proposal distribution for $\underline{\mathbf{K}}$. This should be a good approximation for the target distribution for typical $P_{c}$. We calculate the probability $\text{Pr}\left( k_{i}|D_{i},\underline{\mathbf{x}},P_{c} \right)$ of a particular assignment $k_{i}$ for measurement $i$. From Bayes theorem:

$$\text{Pr}\left( k_{i}|D_{i},\underline{\mathbf{x}},P_{c} \right)=\frac{\text{Pr}\left( D_{i}|k_{i},\underline{\mathbf{x}},P_{c} \right)\text{Pr}\left( k_{i}|\underline{\mathbf{x}},P_{c} \right)}{\text{Pr}\left( D_{i}|\underline{\mathbf{x}},P_{c} \right)}=\frac{\text{Pr}\left( D_{i}|k_{i},\underline{\mathbf{x}},P_{c} \right)\text{Pr}\left( k_{i}|\underline{\mathbf{x}},P_{c} \right)}{\sum_{k_{i}=0}^{N} \text{Pr}\left( D_{i}|k_{i},\underline{\mathbf{x}},P_{c} \right)\text{Pr}\left( k_{i}|\underline{\mathbf{x}},P_{c} \right)}$$

Equation 5

We assume that all non-clutter assignments have equal prior probability giving priors for $k$

$$\text{Pr}\left( k|P_{c},N \right)=\pi_{k}=\left\{ \begin{matrix} P_{c} & \text{if }k=0 \\ \left( 1-P_{c} \right)/N & \text{if }k=1..N \end{matrix} \right.$$

We therefore have from Equation 5 and Equation 3

$$\text{Pr}\left( k_{i}|D_{i},\underline{\mathbf{x}},P_{c} \right)=\frac{a_{i}L_{k_{i}}\pi_{k_{i}}}{\sum_{k_{i}=0}^{N} a_{i}L_{k_{i}}\pi_{k_{i}}}=\frac{L_{k_{i}}\pi_{k_{i}}}{\sum_{k_{i}=0}^{N} L_{k_{i}}\pi_{k_{i}}}$$

Equation 6

##### Model selection

To choose an appropriate number of components to fit to a particular dataset we repeat the Metropolis-Hastings parameter estimation for $N$ from 1 to 9 and then score each fit with the average Bayesian Information Criterion, $AVBIC$, over the parameter samples^21^

$$AVBIC=n_{par}\ln n-\frac{1}{n_{samples}}\sum_{i_{sample}=1}^{n_{samples}} 2\ln\text{Pr}\left( \underline{\mathbf{K}}|P_{c},\underline{\mathbf{D}} \right)\pi_{P_{c}}\text{Pr}\left( \underline{\mathbf{D}}|\underline{\mathbf{K}},\underline{\mathbf{x}} \right)\text{Pr}\left( \underline{\mathbf{x}} \right)$$

where $n_{par}=1+N+n$ is the total number of parameters being optimised for $P_{c}$, $\underline{\mathbf{x}}$ and $\underline{\mathbf{K}}$ respectively and $\text{Pr}\left( \underline{\mathbf{K}}|P_{c},\underline{\mathbf{D}} \right)\pi_{P_{c}}\text{Pr}\left( \underline{\mathbf{D}}|\underline{\mathbf{K}},\underline{\mathbf{x}} \right)\text{Pr}\left( \underline{\mathbf{x}} \right)$ is from Equation 4. The Bayesian Information Criterion is an approximation to the hard-to-calculate Bayesian evidence for comparing different model fits while avoiding overfitting, and the average BIC modifies this to apply to MCMC results. We choose the smallest $N$ for which $AVBIC$ is at least $10$ lower than that for all smaller $N$, i.e. the simplest decomposition (fewest components) which is demanded by the data according to $AVBIC$.

##### Using the Metropolis-Hastings posterior samples

The Metropolis-Hastings process produces a representative set of samples from the full posterior distribution, which will explore uncertainties in all parameters, including in the locations of the separation components and correspondence between measurements and individual components. Before sorting, the components in $\underline{\mathbf{x}}$ from one sample to the next need not correspond to one another (it is a Markov chain of samples). Even after sorting in separation they may not correspond to one another from one sample to the next. When visualising and taking estimates and confidence intervals we wish to reorder these components between samples so that they correctly correspond as well as possible. This is done by reordering the components in $\underline{\mathbf{x}}$ independently in each Metropolis-Hastings sample to best match the order in the best (most probable) sample, ${\underline{\mathbf{K}}}_{best sample}$, as follows.

1. Use the measurement assignments ${\underline{\mathbf{K}}}_{best sample}$, determine the normalised sum of sample data posteriors (see **FLImP measurements as posteriors**) separately for each component, i.e. ${P_{\mathrm{DATA}}}_{,k}\left( x \right)=\frac{\sum_{i=1}^{n} \delta_{k_{i},k}\text{Pr}\left( x|D_{i} \right)}{\sum_{i=1}^{n} \delta_{k_{i},k}}$ where $\delta$ is the Kronecker delta.
2. For each sample, consider all possible permutations of ${\underline{\mathbf{x}}}_{sample}$, scoring each permutation ${\underline{\mathbf{x}}}_{sample}'$ with a probability $P'=\prod_{k=1}^{n} {P_{\mathrm{DATA}}}_{,k}\left( x_{k}' \right)$. The best permutation for this sample will be that which maximises $P'$.
3. To allow that some peaks are less clear than others, this process was repeated, but finding for each sample the reordered subset of $N-N_{ignore}$ components of ${\underline{\mathbf{x}}}_{sample}$ which best matches the narrowest $N-N_{ignore}$ components of ${P_{\mathrm{DATA}}}_{,k}\left( x \right)$, and sorting the ignored components by $x$. This is performed for $N_{ignore}=1,2,..N-2$.
4. Steps 2 and 3 yield several proposed relabelings of components between samples (for $N_{ignore}=0,1,2,..N-2$). Whichever set yields components whose posterior sample distributions in $x$, $\text{Pr}\left( x_{k} \right)$ least overlap with one another is chosen as the best relabellinglabelingted for interpretation of results. The overlap between components $j$ and $l$ is defined as

$$Overlap_{j,l}=\int_{0}^{\Delta x} n_{j}\text{Pr}\left( x_{j}|D \right)n_{l}\text{Pr}\left( x_{l}|D \right)dx$$

- where $n_{j}$ and $n_{l}$ are the number of measurements assigned to these components in the most probable sample ($n_{best,k}=\sum_{i=1}^{n} \delta_{k_{best sample,i},k}$, so weighting overlap by abundance of contributing components). The overlap to be minimised is the summed overlap of all pairs of components, $Overlap=\sum_{j=1}^{N} \sum_{l=j+1}^{N} Overlap_{j,l}$.

After this process we have posterior samples from which clear estimates for separate components can be obtained when the data supports this, and where some components are not clearly resolved this will be clear with large ranges of $x$ for some components.

We are most interested in the distribution of separations revealed by these decompositions, and the proportion of measurements attributed to each, and extract various estimates from the posterior distribution samples. Our plots, such as Figure 1(D), show the posterior distributions of $x$ weighted by $f_{median,k}$, the median proportion of measurements assigned to each component, $f_{median,k}\text{Pr}\left( x_{k} \right)$, coloured separately for each component. The grey background in each plot is $P_{\mathrm{DATA}}\left( x \right)$. In the legend for each plot is the median proportion of measurements assigned to clutter, the median $\text{Pr}\left( x_{k}|D \right)$ and limits of its most compact 68% confidence interval. Underneath each plot is a point showing the individual FLImP measurement locations ($\left| \text{Pr}(r_{\Delta x} \right|D_{i})|$, coloured according to their model assignment, with clutter assignments in black.

We also define the sum of abundance-weighted posteriors of each component, $P_{w}\left( x|D \right)=\sum_{k=1}^{N} f_{median,k}\text{Pr}\left( x_{k}|D \right)$. This ignores the assignment of each component and just asks how much evidence the decomposition gives for the presence of each separation in the sample.
